# Distinct myeloid precursors and their interaction with mesenchymal cells orchestrate the spreading of psoriatic disease from the skin to the joints

**DOI:** 10.1101/2025.08.14.665302

**Authors:** Maria G. Raimondo, Hashem Mohammadian, Mario R. Angeli, Stefano Alivernini, Vladyslav Fedorchenko, Kaiyue Huang, Richard Demmler, Peter Rhein, Cong Xu, Yi-Nan Li, Raphael Micheroli, Zoltán Winter, Aleix Rius Rigau, Charles A. Gwellem, Alina Soare, Markus Luber, Hannah Labinsky, Jiyang Chang, Claudia Günther, Ursula Fearon, Douglas J. Veale, Francesco Ciccia, Jürgen Rech, Michael Sticherling, Tobias Bäuerle, Jörg H. W. Distler, Mariola S. Kurowska-Stolarska, Matthias Mack, Arif B. Ekici, Adam P. Croft, Oliver Distler, Hans M. Maric, Caroline Ospelt, Juan D. Cañete, Maria A. D’Agostino, Georg Schett, Simon Rauber, Andreas Ramming

## Abstract

Psoriatic disease initially affects the skin, but later extends to the joints. Herein, we describe a two-step process that orchestrates spreading of inflammation from the skin to the joints. Induction of psoriatic skin disease in photoconvertible mice, followed by sequencing and computational characterization of skin-derived cells in the joints, identified a unique population of CD2^+^ MHC-II^+^ CCR2^+^ myeloid precursors that built up a skin-derived myeloid cell compartment in the joints. Single-cell cross-species reference mapping and mitochondrial variant tracing showed an orthologue human cell population. Interactome analyses in the joints showed that in a second step, resident regulatory CD200^+^ fibroblasts critically regulate the priming of the CD2^+^ MHC-II^+^ CCR2^+^ myeloid precursors, which subsequently regulate the IL-17 expression in T cells. Hence, spreading of inflammation requires a distinct migratory myeloid precursor population and a permissive local tissue environment, similar to tumour metastasis.

**Graphical abstract:** 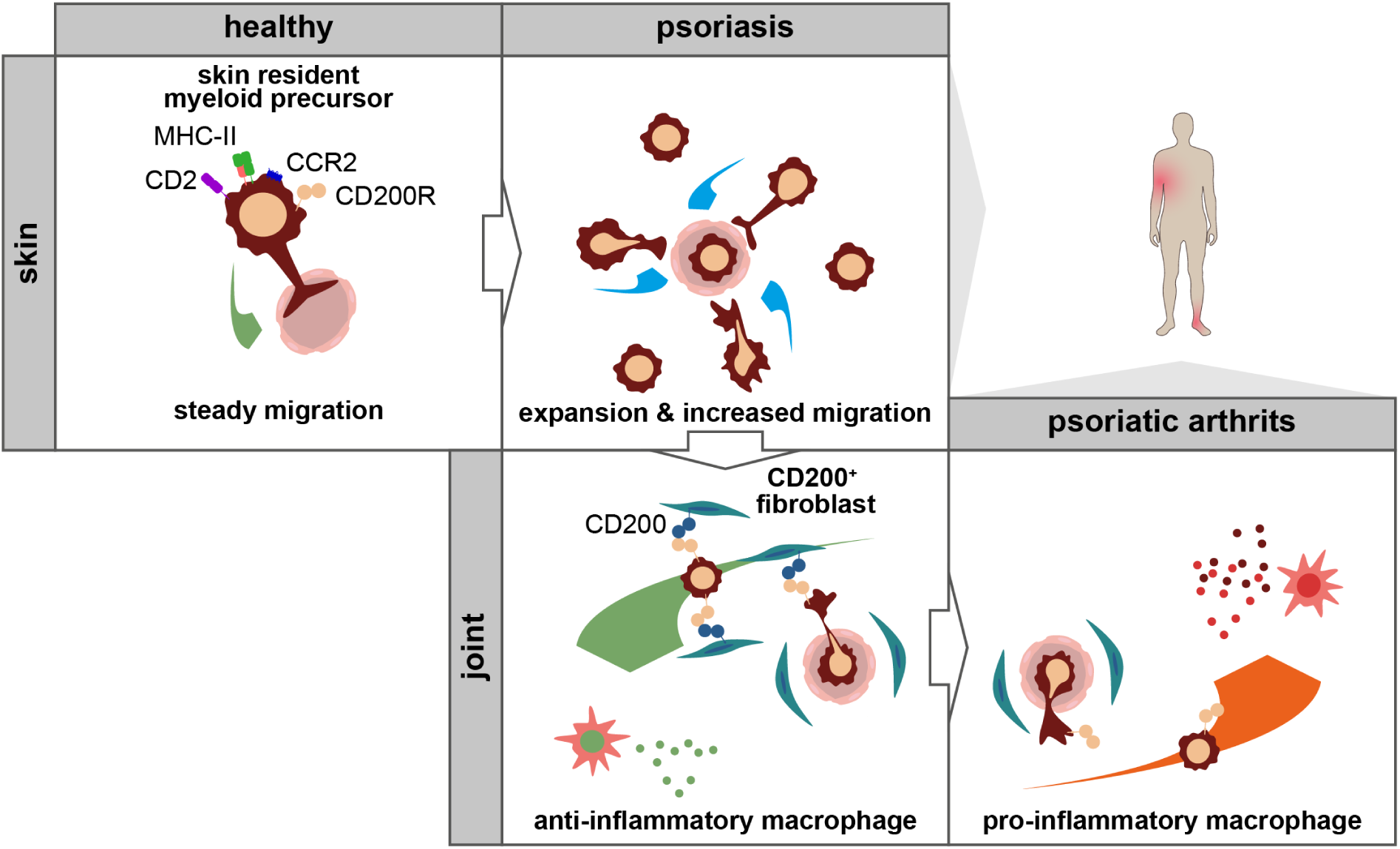

## INTRODUCTION

Psoriatic disease is an immune-mediated inflammatory disease (IMID) that presents with characteristic skin and joint inflammation ^1^. Skin inflammation (psoriasis) precedes joint inflammation (psoriatic arthritis, PsA) in about 80 % of patients ^2^. Approximately 30 % of the patients with psoriasis go on to develop arthritis ^3^, suggesting a directed spatio-temporal link between these two target organs of psoriatic disease. However, how the disease spreads from the skin to the joints is largely unknown.

At the molecular level, psoriatic disease has been linked to the increased activity of the interleukin (IL)-23/-17 signalling and tumour necrosis factor (TNF) pathways. For example, induced systemic overexpression of IL-23 results in psoriasis-like skin and joint disease in autoimmune-prone B10.RIII mice ^4^. Similarly, mice carrying a doxycycline-inducible human TNF transgene develop a psoriasis-like skin inflammation with musculoskeletal involvement ^5^. Consequently, neutralization of these cytokines has been successfully implemented in the clinical setting to alleviate the skin and joint manifestations of psoriatic disease ^6^. However, such therapeutic approaches do not specifically help to understand how inflammation spreads from the skin to the joints in psoriatic disease.

Genome-wide association studies (GWAS) have described genetic differences associated with psoriasis and PsA, such as certain variants of HLA-C and HLA-B, respectively ^7^. However, no molecular mechanism or biomarker has yet been identified to reliably differentiate patients who will develop joint disease from those who will not ^8^. The sequential progression of the disease from skin to joints suggests a mechanistic axis between these two organs. Recent studies in patients with PsA have demonstrated the recirculation of skin-derived tissue-resident memory T cells into the blood ^9, 10^. Clonal expansion of tissue-resident T cells poised to produce IL-17 from lesional to non-lesional psoriatic skin has also been demonstrated ^11, 12, 13^. The mechanism of immune cell recirculation is not well understood, but for monocytes it is thought to be dependent on blockade of the junctional adhesion molecule JAM3 ^14^. It remains unclear whether other immune cell subsets can migrate by this mechanism and whether immune cells can migrate between distinct peripheral tissue compartments such as skin and joint. A systematic unbiased analysis of cell migration in psoriatic disease has not been undertaken. Therefore, a mechanistic model of spreading of inflammation between organs remains to be developed.

Here, we demonstrate the recirculation of skin-derived pro-inflammatory CD2^+^ MHC-II^+^ CCR2^+^ myeloid precursors into the joints in psoriatic disease. While this recirculation contributes to the spreading of psoriatic disease to the joints, it also requires the local mesenchymal permissive microenvironment in the joints that overrides the CD200 immune checkpoint in order to establish full-blown inflammation in the joints.

## RESULTS

### IL-23-induced psoriasis triggers leukocyte migration from the skin to the joints

To disentangle the skin-joint axis in psoriatic disease, we modified the established model of PsA using a hydrodynamically delivered *Il23* vector in B10.RIII mice ^4, 15^ by introducing IL-23 overexpression (IL-23OE) into the more resistant inbred strains BALB/c and C57BL/6 aiming for a model system that would eventually be resistant to arthritis. We found typical symptoms of psoriasis, such as scaling and epidermal hyperplasia, as early as 3 days after induction, and the symptoms remained stable to a similar extent in both strains for up to 21 days (**Figures 1a**, **ED1a** and **ED1d**). We then examined the ankle joints for signs of inflammation using histological haematoxylin and eosin (H & E) staining and magnetic resonance imaging (MRI). We observed early signs of inflammation on day 7 in BALB/c, but not in C57BL/6 mice (**Figures ED1b** and **ED1c**). On day 21, inflammatory arthritis, including dactylitis, tendinitis, enthesitis and synovitis, was observed by histology and MRI, again only in BALB/c but not in C57BL/6 mice (**Figures 1b**, **1c** and **ED1e** - **ED1g**). In addition, both strains exhibited systemic bone loss, presumably due to elevated osteoclast activation (**Figures ED1h** and **ED1i**) ^15^. However, osteoproliferative lesions were only found in animals with inflammatory arthritis (**Figure ED1j**). Thus, we have established a model system that mimics the dichotomy of human psoriasis and enables the identification of molecular differences associated with spreading or non-spreading of inflammation from the skin to the joints.

**Figure 1:**
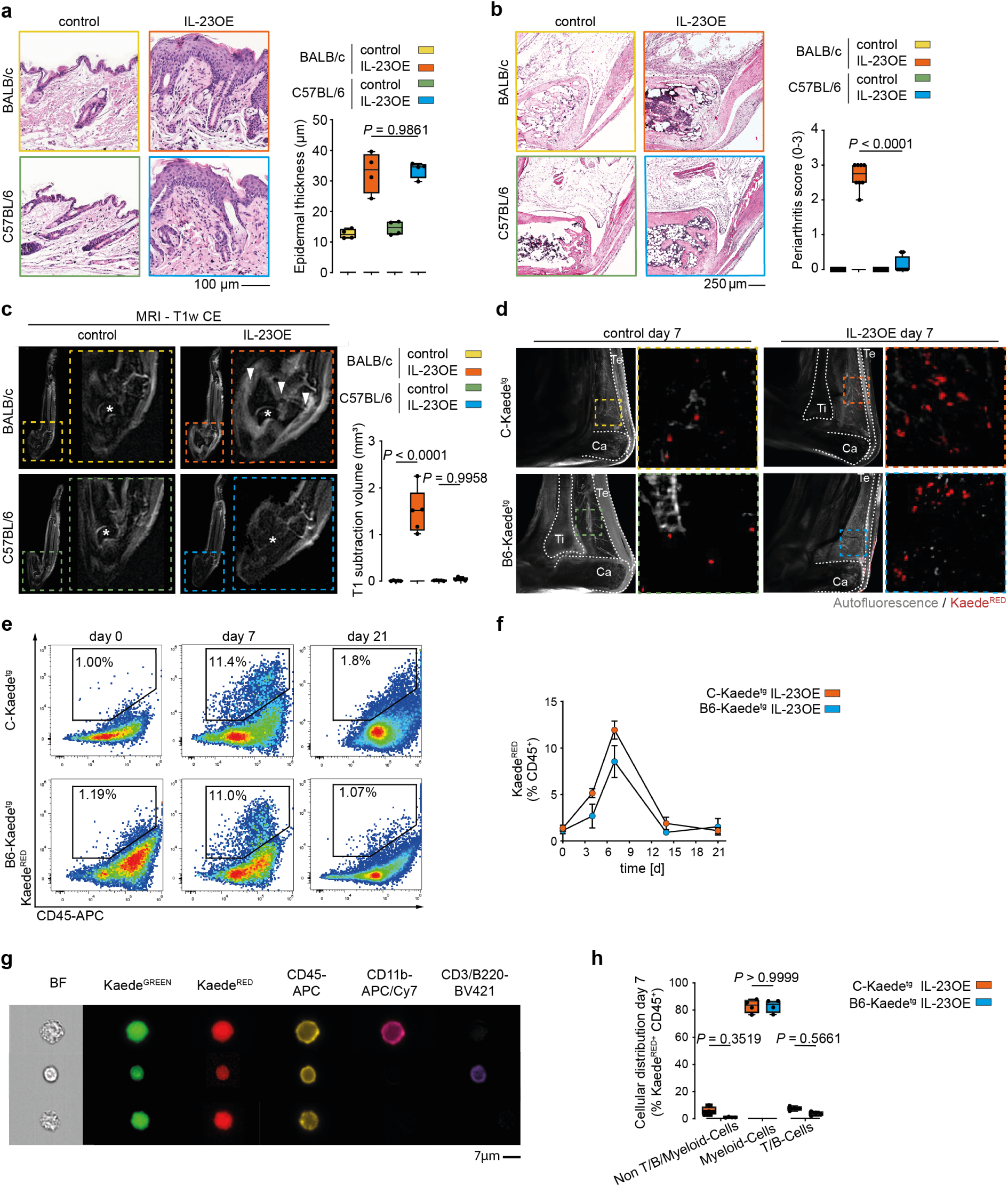
CD11b^+^ myeloid cells migrate from the skin to the joints in a model of psoriatic disease. (**a**) Representative micrographs of haematoxylin & eosin (H & E)-stained skin sections of hind paws of BALB/c and C57BL/6 mice at day 21 with and without IL-23 overexpression (OE); quantification of epidermal thickness at day 21. Graph shows median, quartiles and min-max; *N* = 4 per condition; *P*-values were calculated by one-way ANOVA with Tukeýs *post hoc* test. (**b**) Representative micrographs of H & E-stained ankle sections of BALB/c and C57BL/6 at day 21 with and without IL-23OE; quantification of arthritis at day 21. Graph shows median, quartiles and min-max; *N* = 8 per condition; *P*-values were calculated by one-way ANOVA with Tukeýs *post hoc* test. (**c**) Representative micrographs of magnetic resonance imaging (MRI)-scanned ankles of BALB/c and C57BL/6 at day 21 with and without IL-23OE used for the quantification of arthritis at day 21. Arrowheads indicate inflammation. Stars indicate the talar bone. Graph shows median, quartiles and min-max; *N* = 5 per condition; *P*-values were calculated by one-way ANOVA with Tukeýs *post hoc* test. (**d**) Representative micrographs of light sheet fluorescence microscopy of Kaede^tg^ ankles from BALB/c (C-Kaede^tg^) and C57BL/6 (B6-Kaede^tg^) background strains at day 7 with and without IL-23OE and after photoconversion of cells localized in the skin. Arrowheads indicate accumulations of photoconverted Kaede^RED^ cells. Graphical drawing of tibia (Ti), calcaneus (Ca) and Achilles tendon (Te). (**e**) Representative flow cytometry plots for the quantification of Kaede^RED^ skin-derived cells in the joint. (**f**) Quantification of Kaede^RED^ skin-derived cells in the joints. Graph shows mean and standard error of the mean; *N* = 4 per timepoint and condition. (**g**) Representative micrographs of imaging flow cytometry for the typing of Kaede^RED^ skin-derived cells in the joint at day 7; bright field (BF). (**h**) Quantification of CD45^+^ Kaede^RED^ skin-derived cell types in the joints at day 7. Graph shows mean and standard deviation; *N* = 4 per condition; *P*-values were calculated by one-way ANOVA with Tukeýs *post hoc* test.

To follow the recirculation of cells originating from the skin, we used photoconvertible Kaede^tg^ mice ^16^ and light sheet microscopy to detect trafficking of skin-derived cells to the joints (**Figure ED2a**). IL-23OE induced robust cell migration from the skin into the ankle joints with the cells localising to Kager’s fat pad and the synovium (**Figures 1d** and **ED2b**). Unexpectedly, this migration from the skin to the joints was similar in the arthritis-developing BALB/c strain and the arthritis-resistant C57BL/6 strain.

To further quantify this cellular migration, we used flow cytometry of dissociated ankle tissues and quantified the migration of photoconverted skin-derived Kaede^RED^ cells over time. Migration was restricted to CD45^+^ leukocytes and was observed as early as at day 4, peaked at day 7 and declined thereafter (**Figures 1e**, **1f** and **ED2c**). To further investigate the nature of the skin-derived migrating Kaede^RED^ CD45^+^ leukocytes in more detail, we used imaging cytometry to assess the phenotype of these cells. As expected, we found CD3^+^ T-cells and B220^+^ B-cells in the ankle joint, but also CD11b^+^ (CD3^-^/B220^-^) myeloid cells disseminated from the skin (**Figure 1g**). Unexpectedly, myeloid cells accounted for approximately 85 % of the migrating cells in both strains (**Figure 1h**). The uniform cytosolic distribution of Kaede^GREEN^ and Kaede^RED^ protein also ruled out that phagosomal scavenge transport of Kaede^RED^ protein among myeloid cells was responsible for the Kaede^RED^ signal in the joint.

### Leukocyte trafficking on the skin-joint axis is dominated by myeloid cells

Having observed that cellular migration from psoriatic skin was not predictive of subsequent development of arthritis, we aimed to obtain a detailed profile of the skin-derived and joint-invading leukocytes. Therefore, we generated a comprehensive single-cell RNA sequencing (scRNAseq) dataset of skin- and joint-resident Kaede^GREEN^ leukocytes (CD45^+^), stromal cells (CD45^-^) and Kaede^RED^ migrated leukocytes (CD45^+^) (**Figure ED3a**). Using uniform manifold approximation and projection (UMAP) for visualization, we again found the majority of Kaede^RED^ migrated leukocytes in the joint tissue belonging to the myeloid phagocyte cluster (**Figures 2a** and **ED3b**). In a subsequent clustering of this compartment, we were able to identify five different clusters in agreement with previous reports ^17, 18, 19, 20^, namely MHC-II^-^ lining layer macrophages, *Hmox1*^+^ macrophages, *Cd163*^+^ *Lyve1*^+^ anti-inflammatory macrophages, *Aqp1*^+^ macrophages and MHC-II^+^ (*H2^+^)* phagocytes, containing a diverse cellular composition. Three of the five subclusters, namely *Hmox1*^+^ macrophages, *Cd163*^+^ *Lyve1*^+^ anti-inflammatory macrophages and *H2*^+^ phagocytes, were significantly enriched in Kaede^RED^ migrated cells (**Figure 2b**). However, there was no difference in the abundance of the three migrated subclusters between the two mouse strains, confirming the observation that migration alone is not sufficient to trigger the spread of inflammation from the skin to the joints (**Figure 2c**).

**Figure 2:**
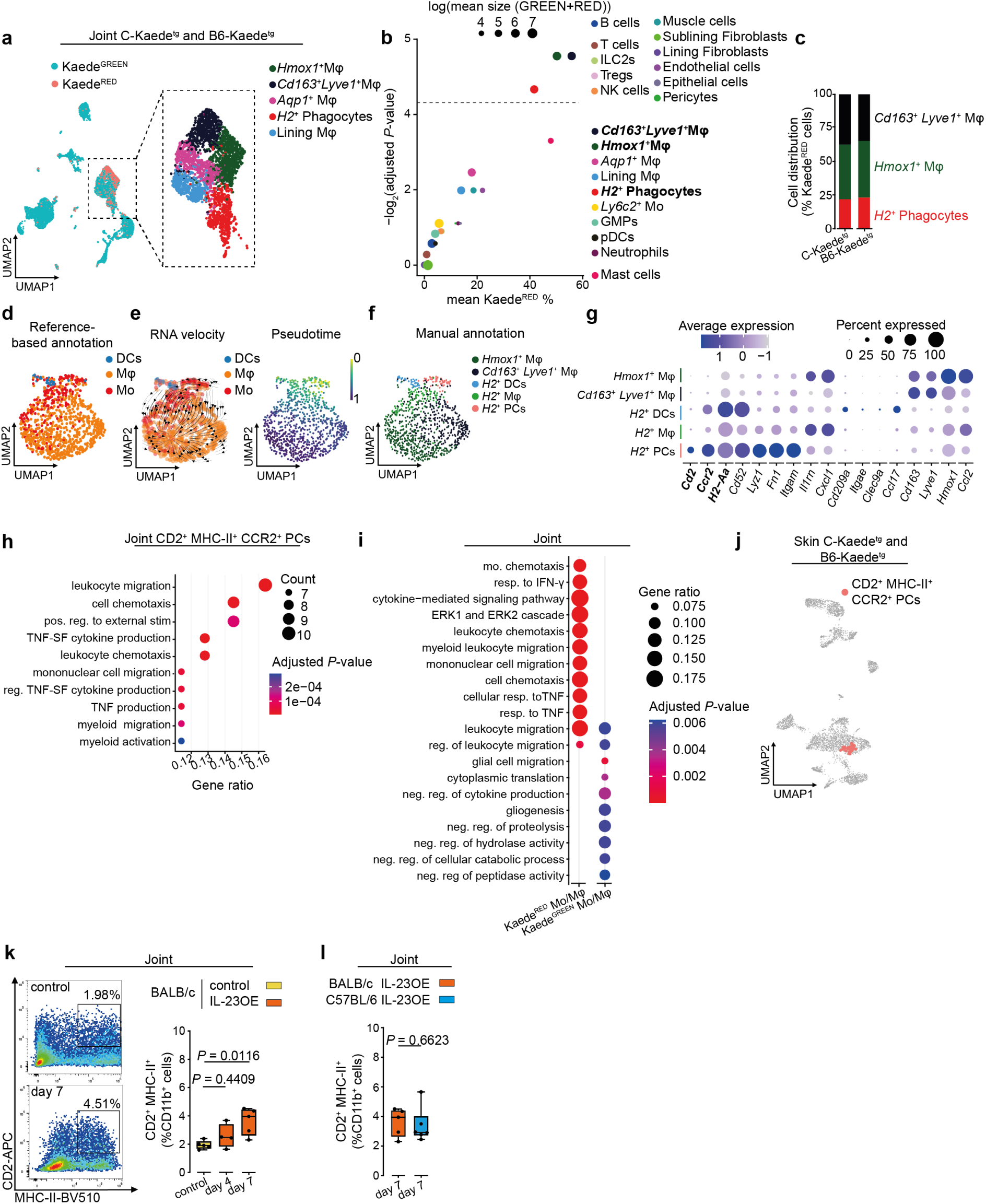
Skin-derived myeloid cells are precursors that contribute to the pool of mononuclear phagocytes in the joint. (**a**) UMAP plot of the hash-tag identified ankle joint cells in the scRNAseq dataset from Kaede^tg^ mice on BALB/c and C57BL/6 background on day 7 of the IL-23 overexpression (OE) psoriasis model. Kaede^GREEN^ and Kaede^RED^ cells are highlighted (left). Sub-cluster visualization of myeloid phagocytes with apparent highest abundance of Kaede^RED^ cells (right). (**b**) Scatter plot corresponding to **a** showing the log transferred cell abundance and the Kaede^RED^ percentage in each of the identified clusters versus the Benjamini and Hochberg (BH)-adjusted *P*-values calculated using quasibinominal models. The dashed line indicates the significance threshold (BH-adjusted *P*-value <0.05). (**c**) Bar plot comparing the ratio of clusters of myeloid cells with highest Kaede^RED^ percentage between strains. (**d**) Reference-based annotation of each single cell of the Kaede^RED^ myeloid cells based on the ImmGen’s mononuclear phagocyte dataset (GSE122108) identified by singleR. (**e**) Velocity streams (left) and pseudo-time (right) identified by CellRank of the Kaede^RED^ myeloid cells visualized over the UMAP plot. (**f**) UMAP visualization of Kaede^RED^ myeloid cells with subclusters of *H2*^+^ phagocytes after manual annotation. Abbreviation: precursors (PCs). (**g**) Expression of the most relevant marker genes among each myeloid cluster from **f**. (**h**) Significantly enriched GO-BP terms in the CD2^+^ MHC-II^+^ CCR2^+^ myeloid precursors cluster, when compared to all other cell types in the joint. Selection criteria: BH-adjusted *P*-value < 0.05. (**i**) Comparison of the significantly enriched GO-BP terms between Kaede^RED^ and Kaede^GREEN^ myeloid cells. Selection criteria: BH-adjusted *P*-value < 0.05. (**j**) Identified CD2^+^ MHC-II^+^ CCR2^+^ myeloid precursors using UCell, highlighted on the UMAP plot of the hash-tag identified skin leukocytes in the scRNAseq dataset from Kaede^tg^ animals with BALB/c and C57BL/6 background strains on day 7 of the IL-23OE. (**k**) Representative flow cytometry plots for the quantification of CD2^+^ MHC-II^+^ CCR2^+^ skin-derived myeloid precursors in the joint (left) and their quantification in BALB/c over time. Graph shows median, quartiles and min-max.; *N* = 5, *N* = 4 and *N* = 5 per timepoint and condition; *P*-values were calculated by one-way ANOVA with Tukeýs *post hoc* test. (**l**) Quantification of CD2^+^ MHC-II^+^ CCR2^+^ skin-derived myeloid precursors in the joint of BALB/c and C57BL/6 on day 7. Graph shows median, quartiles and min-max.; *N* = 5 and *N* = 6 per timepoint and condition; *P*-values were calculated by Mann-Whitney *U*-test.

### CD2^+^ MHC-II^+^ CCR2^+^ myeloid precursors are skin-derived joint-invading cells

To better characterize the Kaede^RED^ myeloid compartment, we first performed a reference-based annotation using the Immunological Genome Project (ImmGen) mononuclear phagocytes dataset (GSE122108) ^21, 22^. This approach allowed for a clustering-independent annotation of each single cell, which identified monocytes, macrophages and dendritic cells (DCs) (**Figure 2d**). We then examined the transcriptional dynamic and differentiation potential of Kaede^RED^ phagocytes. Transcriptional turnover estimated by RNA velocity when combined with transcriptional similarity using CellRank ^23^ identified several macro-states among these cells (**Figure 2e** and **ED3c**). However, CellRank identified *H2*^+^ precursors as the only initial state with the potential to differentiate not only into *H2*^+^ phagocytes, but also into *Hmox1*^+^ and *Cd163*^+^ *Lyve1*^+^ macrophages (**Figure ED3d**), which perfectly aligned with CellRank’s assisted pseudo-time estimation. We therefore extended our initial annotation of *H2*^+^ phagocytes and subclustered them into *H2*^+^ DCs, *H2*^+^ macrophages and *H2*^+^ myeloid precursors (**Figure 2f**), which also overlapped precisely with the reference-based annotation. Among the *H2*^+^ phagocytes, the *H2*^+^ myeloid precursors could be distinguished by the expression of *Cd2* and *Ccr2* (**Figure 2g**). We will therefore refer to them as CD2^+^ MHC-II^+^ CCR2^+^ myeloid precursors. Consistent with our findings, gene ontology (GO) biological processes (GO-BP) enrichment analysis revealed a significant contribution of migration-associated pathways when comparing CD2^+^ MHC-II^+^ CCR2^+^ myeloid precursors with others subsets in the joints (**Figure 2h**). Similarly, the comparison of migrated Kaede^RED^ with resident Kaede^GREEN^ phagocytes revealed a higher abundance of chemotaxis- and migration-related terms in the Kaede^RED^ population (**Figure 2i**). We also identified a pure Kaede^GREEN^ cluster of potential precursors (**Figure ED3b**), *Ly6c2*^+^ monocytes ^17^, which were characterised by a high metabolic activity based on GO-BP enrichment analysis (**Figure ED3e**).

Given the CellRank analysis on the likelihood of differentiation and the GO-BP enrichment analysis, we hypothesized that a CD2^+^ MHC-II^+^ CCR2^+^ myeloid precursor population would be found in the skin. To explore this possibility, we used the transcriptional profile of joint CD2^+^ MHC-II^+^ CCR2^+^ myeloid precursors to score the skin myeloid compartment by UCell ^24^ and identified CD2^+^ MHC-II^+^ CCR2^+^ myeloid precursors among the monocytes in the skin **(Figures 2j** and **ED3f)**. CD2^+^ MHC-II^+^ CCR2^+^ myeloid precursors in the skin showed a significant enrichment of GO-BP terms associated with cellular migration **(Figure ED3g)**.

To test cellular trafficking of CD2^+^ MHC-II^+^ CCR2^+^ myeloid precursors between skin and joints independently of the Kaede^tg^ system, we developed a flow cytometry strategy to detect CD2^+^ cells among CD11b^+^ MHC-II^+^ cells in the joints and the skin (**Figure ED3h**). Compared to untreated mice, IL-23OE induced a significant expansion of CD2^+^ MHC-II^+^ myeloid precursors in skin and joints in BALB/c mice on days 4 and 7 (**Figures 2k** and **ED3i**). Notably, no difference was observed between BALB/c and C57BL/6 inbred strains (**Figure 2l**). We did not observe a significant increase in CD2^+^ MHC-II^+^ myeloid precursors in the bone marrow and small intestine following IL-23OE (**Figures ED3j** and **ED3k**). Given that both organs are affected by IL-23OE ^15, 25^, this finding supports the concept that the skin is the source of these cells in the joint.

### Leukocyte trafficking is conserved between mice and humans

We next asked whether similar migration mechanisms exist in humans. Therefore, we integrated scRNAseq datasets from synovial tissues (healthy *N* = 3, PsA *N* = 5, E-MTAB-11791 ^26^), identified myeloid cells, subclustered them into eleven distinct cellular identities by unsupervised clustering and labelled them according to their characteristic gene expression profile (**Figures 3a**, **ED4a** and **ED4b**). We then aimed to find biologically relevant human-mouse orthologous populations within the myeloid compartment. Therefore, we used scANVI (single-cell annotation using variational inference) ^27^ integration and reference mapping, which showed optimal results in a benchmarking comparison ^28^, to map our mouse annotation (**Figure 2f**) to the human synovial myeloid subclusters (**Figure 3a**). In addition to mouse and human DC subsets, we observed the highest degree of co-occurrence between mouse CD2^+^ MHC-II^+^ CCR2^+^ myeloid precursors and human *CCR2*^+^ monocyte subclusters (**Figure 3b**). The human *CCR2*^+^ monocytes expressed *CD2* and *HLA-DR* in agreement with the mouse CD2^+^ MHC-II^+^ CCR2^+^ myeloid precursors (**Figures ED4b** and **ED4c**). The reverse detection of human myeloid precursor signatures among mouse myeloid cells using UCell ^24^ confirmed the high degree of similarity between human and mouse CD2^+^ MHC-II^+^ CCR2^+^ myeloid precursors (**Figure ED4d**).

**Figure 3:**
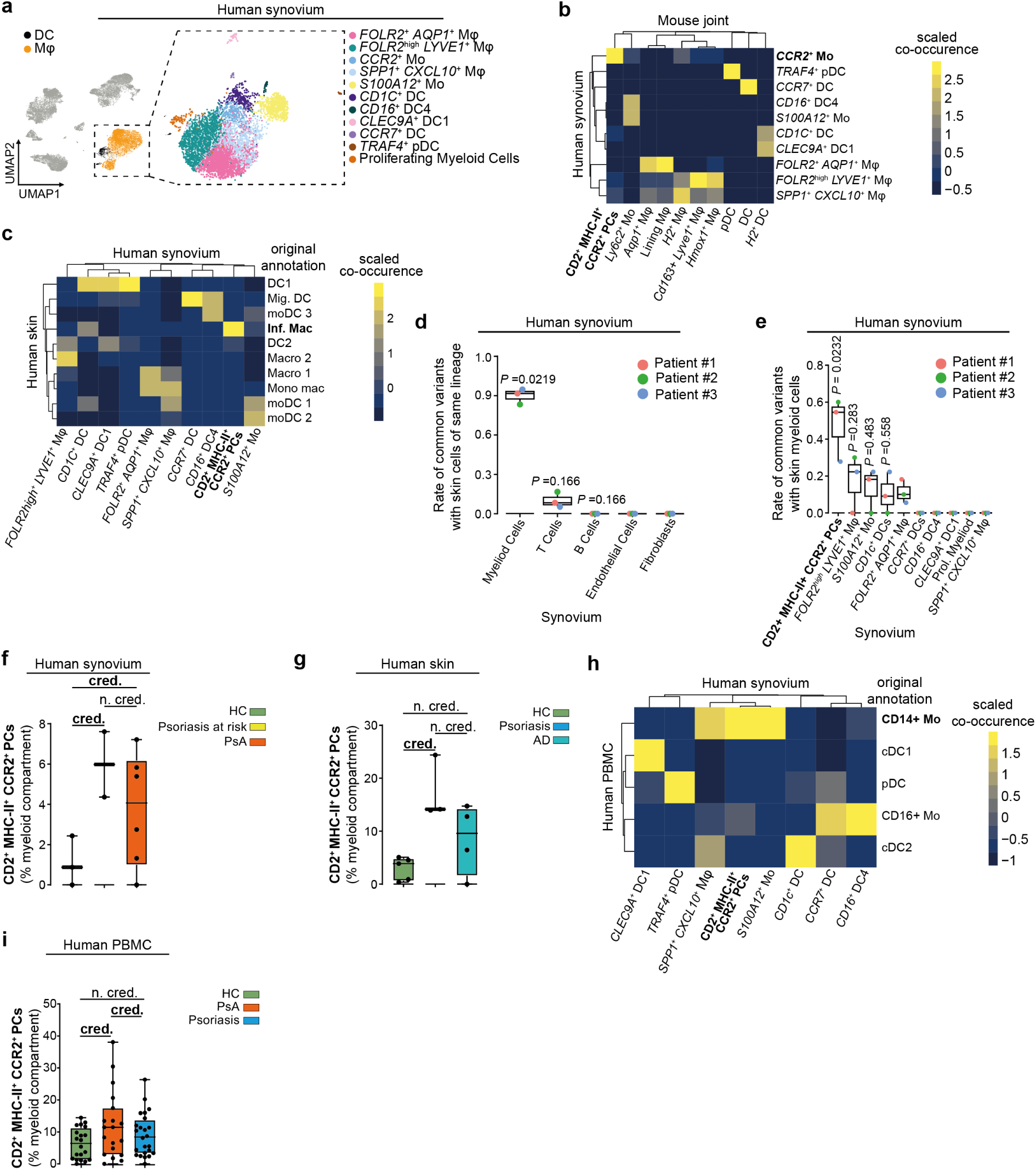
Evidence for a skin-joint axis in human. (**a**) UMAP plot of the scVI-integrated scRNAseq datasets of human psoriatic arthritis (PsA) synovium (*N* = 5, E-MTAB-11791) and healthy synovial tissue (*N* = 3). Identified clusters of myeloid cells are highlighted. (**b**) Heatmap of co-occurrence of psoriatic mouse myeloid cell clusters with human PsA myeloid cell clusters when mouse joint annotations were mapped to human cells using scANVI. (**c**) Heatmap of co-occurrence of human synovial myeloid cell clusters (PsA *N* = 5, healthy *N* = 3) with skin myeloid cell clusters from a human skin scRNAseq dataset (healthy subjects *N* = 5, psoriasis *N* = 3, atopic dermatitis (AD) *N* = 4, E-MTAB-8142) when synovial annotations were mapped to skin annotations (retrieved from metadata) using scANVI. (**d**) Proportions of shared mitochondrial variants between synovial and skin clusters of the same lineage for the major clusters in the synovia. Graph shows median, quartiles and min-max.; *N* = 3; *P*-values were calculated by one-vs-rest Wilcoxon rank-sum test with BH correction. (**e**) Boxplot corresponding to **d**. The synovial myeloid compartment is shown in detail. (**f**) Proportion of CD2^+^ MHC-II^+^ CCR2^+^ myeloid precursors in psoriasis at risk and PsA compared to healthy in the scRNAseq dataset of human synovial tissue. Graph shows median, quartiles and min-max.; psoriasis at risk *N* = 2; PsA *N* = 5, healthy subjects (HC) *N* = 3; Statistically credible (cred.) and non-credible (n. cred.) changes in abundance were identified using scCODA. (**g**) Proportion of CD2^+^ MHC-II^+^ CCR2^+^ myeloid precursors in AD and psoriasis compared to healthy in the scRNAseq dataset of human skin. Graph shows median, quartiles and min-max.; AD *N* = 4; psoriasis *N* = 3, HC *N* = 5; Statistically credible (cred.) and non-credible (n. cred.) changes in abundance were identified using scCODA. (**h**) Heatmap of co-occurrence of human myeloid cell clusters from joint (PsA *N* = 5, healthy subjects *N* = 3) with PBMC myeloid cell cluster from proteogenomic dataset (healthy subjects *N* = 20, PsA *N* = 19, psoriasis *N* = 24, GSE194315) when joint annotations were mapped to the pre-annotated PBMCs using scANVI. (**i**) Proportion of CD2^+^ MHC-II^+^ CCR2^+^ myeloid precursors in psoriasis and PsA compared to healthy in the scRNAseq dataset of human PBMCs. Graph shows median, quartiles and min-max.; psoriasis *N* = 24; PsA *N* = 19, HC *N* = 20; Statistically credible (cred.) and non-credible (n. cred.) changes in abundance were identified using scCODA.

To trace back CD2^+^ MHC-II^+^ CCR2^+^ myeloid precursors to the skin, we integrated the human synovial myeloid cell clusters from healthy and PsA donors into an existing scRNAseq dataset of skin myeloid cells (E-MTAB-8142 ^29^) containing healthy donors (*N* = 5), psoriasis (*N* = 3) and atopic dermatitis (AD, *N* = 4) patients. When the signatures of the synovial myeloid cells were mapped to the pre-annotated dermal myeloid cells using scANVI transfer, we observed a high degree of co-occurrence between synovial CD2^+^ MHC-II^+^ CCR2^+^ myeloid precursors and dermal myeloid cells that were originally termed ‘inflammatory macrophageś (**Figure 3c**) ^29^.

Having observed human orthologous populations of mouse CD2^+^ MHC-II^+^ CCR2^+^ myeloid progenitors in the skin and synovium, we sought to track their migration between these two tissues in patients. We hypothesised that cells migrating from the skin to the joint would exhibit a significantly higher prevalence of shared somatic mitochondrial mutations than either resident cells or cells originating from distinct sources within both organs. To this end, we generated three matched scRNAseq datasets of skin and joint tissue from three patients (**Figures S1a**-**d**, early PsA *N* = 1, psoriasis at risk of developing PsA *N* = 2). Additionally, we enriched and sequenced the cDNA specifically for mitochondrial transcripts to detect mitochondrial somatic mutations using the MAESTER (mitochondrial alteration enrichment from single-cell transcriptomes to establish relatedness) method ^30^, which enabled us to establish these lineage relationships. Using conservative quality and specificity thresholds for informative variant detection, we identified median 12 (IQR 3.5) shared variants between skin and synovium. Consistent with our hypothesis, the majority of these variants were found in myeloid cells, with a smaller number found in T cells (**Figure 3d**). Notably, CD2^+^ MHC-II^+^ CCR2^+^ myeloid precursors exhibited the highest enrichment of conserved somatic mitochondrial mutations (**Figure 3e**). We considered the possibility that the methodology might have favoured myeloid cells, given their presumably shorter lifespan and recent origin from bone marrow precursors, as this could result in greater preservation of somatic variants. However, to exclude this possibility, we analysed S100A12⁺ monocytes infiltrating the synovia ^18^ and found that they had relatively few shared variants with their skin counterparts comparable to T cells (**Figure**

**ED4e**). Furthermore, we used a graph-based lineage tracing approach to rank the synovial myeloid clusters based on the extent of shared variants among them, both including and excluding those shared with skin (**Figure S1e**). This approach identified CD2⁺ MHC-II⁺ CCR2⁺ myeloid cells as the earliest precursors, provided the conserved skin variants were included (**Figure ED4f**). This analysis also provided proof that the shared variants in the other joint myeloid clusters were derived from CD2⁺ MHC-II⁺ CCR2⁺ myeloid precursors, ruling out other migratory sources. Finally, to exclude the possibility that our selection criteria for informative variants had affected the results, we performed an iterative analysis, defining variable ranges for each selection criterion. As expected, myeloid cells consistently exhibited the highest number of shared variants with skin cells in 100% of the iterations (**Figures ED4g** and **ED4h**). Within the myeloid clusters, CD2⁺ MHC-II⁺ CCR2⁺ myeloid precursors were consistently ranked first in approximately 70 % of iterations (**Figures ED4i** and **ED4j**). Taken together, these results demonstrate not only the presence of a human orthologue of CD2⁺ MHC-II⁺ CCR2⁺ myeloid precursors, but also that their migration between skin and synovium is conserved in humans.

### Disease-specific expansion of myeloid precursors

We further explored whether the disease context would influence the number of CD2^+^ MHC-II^+^ CCR2^+^ myeloid precursors in affected organs in humans. Consistent with our observation in mice, we observed that CD2^+^ MHC-II^+^ CCR2^+^ myeloid precursors were significantly elevated in the synovial tissues of psoriatic arthritis (PsA) patients compared to healthy controls. We detected this increase already in psoriasis patients at risk of developing PsA (**Figures 3f** and **S2a**). In contrast, when we analysed the synovial myeloid compartment from treatment-naïve rheumatoid arthritis (RA) patients (*N* = 5, E-MTAB-8322 ^18^) and annotated it according to our healthy/PsA labels (**Figure ED4k**), we did not detect any *CD2* expression in the precursor-labelled cells in RA patients (**Figure ED4l**). This highlights the unique disease-specific and skin-specific signature of CD2^+^ MHC-II^+^ CCR2^+^ myeloid precursors.

Analysing the differential abundance of monocytic populations between psoriasis and healthy human skin, we observed that CD2^+^ MHC-II^+^ CCR2^+^ myeloid precursors were significantly more likely to be enriched in psoriasis patients compared to healthy controls (**Figures 3g** and **S2b-d**). As a proof of concept, we did not observe a significant increase of CD2^+^ MHC-II^+^ CCR2^+^ myeloid precursors in AD, which has an aetiology different from psoriatic disease (**Figures S2b** and **S2d**). These findings confirmed the co-occurrence of CD2^+^ MHC-II^+^ CCR2^+^ myeloid precursors in skin and synovial tissue, which increased under pathological conditions, consistent with our observations in mice.

Finally, we tested whether skin-disseminating recirculating precursors would be detectable in the human PBMC blood fraction to support the hypothesis of their migration along the skin-joint axis. We therefore integrated the scRNAseq datasets of myeloid cells from healthy and PsA synovial tissues with an existing dataset of PBMCs (GSE194315 ^31^) from healthy donors (*N* = 20) and patients with psoriatic disease (psoriasis: *N* = 24, PsA: *N* = 19). In addition to the gene expression data, GSE194315 partially included proteogenomic data that allowed differentiation of cell populations based on antibody stainable surface markers. When the synovial annotations were mapped to the proteogenomic-based pre-annotation of PBMCs using scANVI transfer, we observed a high degree of co-occurrence between synovial CD2^+^ MHC-II^+^ CCR2^+^ myeloid precursors and CD14^+^ blood monocytes (**Figure 3h**). The mapped synovial CD2^+^ MHC-II^+^ CCR2^+^ myeloid precursors formed a subcluster among the CD14^+^ blood monocytes, which was revealed as a distinct population of antibody-identifiable CD2^+^ MHC-II^+^ CCR2^+^ myeloid precursors (**Figures ED4m** and **ED4n**). Finally, we compared the abundance of myeloid populations between healthy controls, psoriasis and PsA patients, which showed that CD2^+^ MHC-II^+^ CCR2^+^ myeloid precursors were increased in the blood of PsA patients (**Figures 3i**, **S2e** and **S2f**). In conclusion, we have provided evidence that CD2^+^ MHC-II^+^ CCR2^+^ myeloid precursors are increased not only in the circulation but also in the skin and the synovial tissue of patients with psoriatic disease, fully mirroring the observations made in the animal models.

### Resident fibroblasts in the joint influence the activation of skin-derived mononuclear phagocytic cells

As we had observed that migrating CD2^+^ MHC-II^+^ CCR2^+^ myeloid precursors are a rather dynamic population once they arrive in the joint, giving rise to several subpopulations, we wondered whether the local tissue microenvironment would influence these differentiation processes. Therefore, we compared the change in gene expression profiles of CD2^+^ MHC-II^+^ CCR2^+^ myeloid precursors between BALB/c and C57BL/6 over pseudo-time using tradeSeq ^32^ (trajectory-based differential expression analysis for sequencing data) (**Figure 4a**). While minimal changes were observed in arthritis-resistant C57BL/6 mice, BALB/c mice showed a much greater divergence from baseline. Consequently, when we compared the enriched GO-BP among the differentially dynamic genes, we found several pro-inflammatory pathways that were activated in BALB /c mice, which develop clinical signs of arthritis, compared to C57BL/6 mice, resistant to arthritis (**Figure ED5a**). Correspondingly, when we scored the cells within the myeloid phagocyte clusters (**Figure 2a**) with UCell for pro- and anti-inflammatory responses, we observed a higher pro-inflammatory response in the Kaede^RED^ cells from BALB/c mice, but a higher anti-inflammatory response in the Kaede^RED^ cells from C57BL/6 mice (**Figure 4b**). In contrast, the Kaede^GREEN^ compartment, consisting mainly of lining layer macrophages and *Aqp1*^+^ macrophages, showed neither relative significant anti-inflammatory nor pro-inflammatory activation in both background strains. These analyses demonstrate on the one hand that skin-derived Kaede^RED^ cells are the key regulators of inflammation and on the other hand that the joint microenvironment plays a key role in priming this compartment for pro- or anti-inflammatory activation. Therefore, we next sought to unravel the interactome of precursors with the local microenvironment using CellChat ^33^. Intercellular interaction analysis revealed a robust communication probability between Kaede^RED^ phagocytes and their microenvironment in both strains, and the overall communication strength of the Kaede^RED^ migrated cells was higher than that of Kaede^GREEN^ putative resident cells (**Figure 4c**). The strongest communication probabilities were found between Kaede^RED^ phagocytes and the fibroblast compartment, and the strength of the interaction was predicted to be higher in C57BL/6 compared to BALB/c mice. To disentangle the putative interaction partners, we subclustered the fibroblast compartment in the joints and annotated seven distinct populations according to previous reports (**Figures 4d** and **4e**) ^34, 35, 36, 37^. There was a clear separation between synovial lining and sublining fibroblast subtypes, which were further stratified into *Thbs1*^+^, *Il6*^+^, *Cd200*^+^ and *Pi16*^+^ fibroblast subtypes as well as chondrocytes and osteoblasts. As we had observed a higher probability of interaction between fibroblasts and Kaede^RED^ phagocytes, we calculated the UCell scores of fibroblast-specific differentially expressed genes in C57BL/6 compared to BALB/c mice. We found the highest scores among the *Cd200*^+^ fibroblasts (**Figure 4f**), arguing that this population was the critical determinant in the microenvironment between BALB/c and C57BL/6 mice. Furthermore, MELD ^38^ analysis showed that the *Cd200*^+^ fibroblasts were more likely to be found in C57BL/6 compared to BALB/c mice exposed to IL-23OE (**Figure 4g**). Finally, we used flow cytometry to quantify the abundance of CD200^+^ fibroblasts (CD45^-^ CD31^-^ PDPN^+^ PDGFRα^+^ CD200^+^ THY1^+^ CD49f^-^) in joints before and after IL-23OE on day 7 ^34^. In consistence with the scRNAseq dataset, we observed a significant decrease in CD200^+^ fibroblasts after IL-23OE in BALB/c, but not in C57BL/6 mice (**Figure 4h**). This decrease coincided with the peak of the migration (**Figure 4i**). Therefore, we hypothesised that the onset of arthritis would be controlled not only by the migration of CD2^+^ MHC-II^+^ CCR2^+^ myeloid precursors from the skin to the joint, but also by CD200^+^ fibroblasts in the joint.

**Figure 4:**
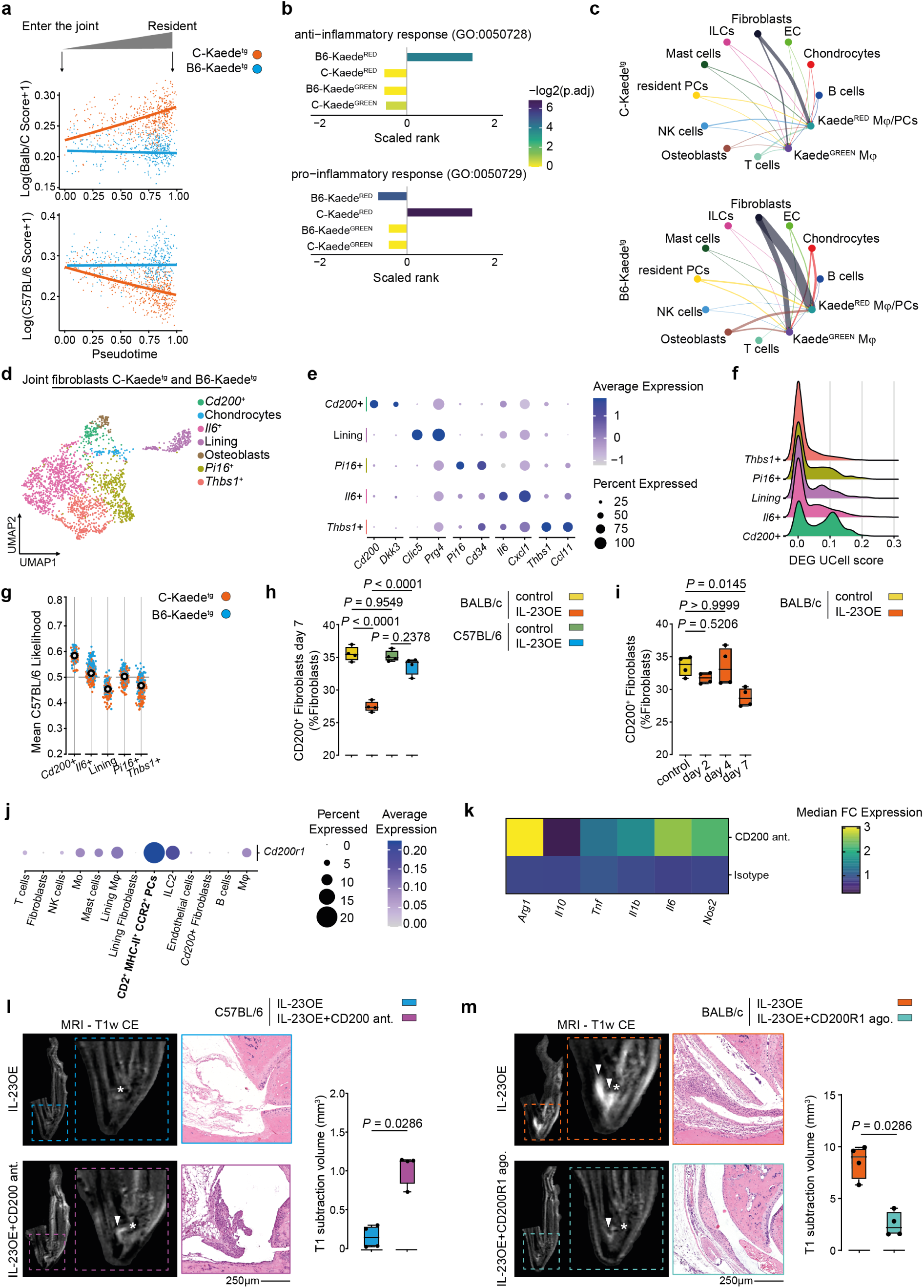
Local mesenchymal cells in the joint prime migrating precursors to a pro- or anti-inflammatory phenotype. (**a**) TradeSeq-fitted smooth expression of UCell scores of the genes dynamically associated with either BALB/c strain (top) or C57BL/6 strain (bottom) over pseudo-time from Figure 2e in the scRNAseq dataset of ankle joints from Kaede^tg^ mice (BALB/c and C57BL/6 strains) on day 7 of the IL-23 overexpression (OE) model. **(b)** UCell scores for GO-BP terms GO:0050728 - anti-inflammatory response and GO:0050729 - pro-inflammatory response on the phagocytes compartment from Figure 2a stratified by stain and photoconversion status. *P*-values were calculated with Wilcoxon signed-rank test and adjusted for multiple comparison using BH method. (**c**) Circle plot showing the communication probability between myeloid cells (Kaede^GREEN^ or Kaede^RED^) and other cell types in the joints associated with either the BALB/c (top) or the C57BL/6 (bottom) strains. Line thickness corresponds to communication probability determined using CellChat. Abbreviation: precursors (PCs). (**d**) UMAP plot of the Seurat identified sub-clusters among fibroblasts in the scRNAseq dataset of ankle joints from Kaede^tg^ mice (BALB/c and C57BL/6 strains) on day 7 of the IL-23OE model (**e**) Expression of the most relevant marker genes among each fibroblasts clusters identified in **d**. (**f**) Comparison of UCell scores of the C57BL/6 associated gene signature (fibroblasts-only differentially expressed genes in C57BL/6 compared to BALB/c) between different subclusters of joint fibroblasts. (**g**) C57BL/6 associated relative likelihood of different subclusters of fibroblasts determined by MELD. (**h**) Quantification of CD200^+^ fibroblasts in the joints of BALB/c and C57BL/6 mice on day 7. Graph shows median, quartiles and min-max.; *N* = 4 per timepoint and condition; *P*-values were calculated by one-way ANOVA with Tukeýs *post hoc* test. (**i**) Quantification of CD200^+^ fibroblasts in the joints of BALB/c mice on day 2, 4 and 7. Graph shows median, quartiles and min-max.; *N* = 4 per timepoint and condition; *P*-values were calculated by one-way ANOVA with Tukeýs *post hoc* test. (**j**) Expression of *Cd200r1* among the different cell clusters in the scRNAseq dataset of ankle joints from Kaede^tg^ mice (BALB/c and C57BL/6 strains) on day 7 of the IL-23 overexpression (OE) model. (**k**) Heatmap of median fold-change gene expression in macrophage-fibroblasts co-cultures treated with anti-CD200 antagonist (OX-90) or isotype; *N* = 7 (**l**) Representative images of MRI scans and micrographs of haematoxylin & eosin (H & E)-stained ankles of C57BL/6 animals at day 21 after IL-23OE with or without anti-CD200 antagonist (OX-90) treatment. MRI-quantification of arthritis at day 21. Arrowheads indicate inflammation. Stars indicate the talar bone. Graph shows median, quartiles and min-max; *N* = 4 per condition; *P*-values were calculated by Mann-Whitney U-test. (**m**) Representative images of MRI scans and micrographs of H & E-stained ankles of BALB/c animals at day 21 after IL-23OE with or without anti-CD200R1 agonist (OX-110) treatment. MRI-quantification of arthritis at day 21. Arrowheads indicate inflammation. Stars indicate the talar bone. Graph shows median, quartiles and min-max; *N* = 4 per condition; *P*-values were calculated by Mann-Whitney U-test.

### CD200-CD200R1 axis controls onset of arthritis

CD200 is known to interact with four different receptors ^39^, of which CD200R1 was the most abundantly expressed in our mouse scRNAseq dataset. Consistent with our previous report ^34^, we found *Cd200r1* transcripts in a few T cells and strong expression on innate lymphoid cells of type 2 (ILC2s) and myeloid cells (**Figure 4j**). In particular, the high cellular resolution of our dataset in the myeloid cells allowed us to analyse the expression in this compartment in more detail and we observed the highest abundance of *Cd200r1* transcripts in CD2^+^ MHC-II^+^ CCR2^+^ myeloid precursors. There was no significant difference in expression between the background strains. Functionally, when we cultured bone marrow monocytes (BMMs) with synovial fibroblasts and blocked the CD200-CD200R1 interaction with the monoclonal antibody OX-90 against CD200, we observed the induction of a pro-inflammatory phenotype in BMMs with higher expression of *Il1b, Il6* and *Tnf* (**Figures 4k** and **ED5b**). These findings suggested that CD200^+^ fibroblasts form a protective barrier in the joint through the CD200-CD200R1 axis.

To further explore this hypothesis, we followed four different approaches. First, we examined whether increasing the number of migrating cells could amplify inflammation. Therefore, on day 7 of IL-23OE in BALB/c mice, we isolated skin-derived CD2^+^ MHC-II^+^ CCR2^+^ myeloid precursors for adoptive transfer. We also isolated a population of CD11c⁻ CCR7⁻ CLEC10A⁺ CD11b^+^ macrophages corresponding to the Kaede^RED^ population in the joint (**Figure S3**). Then, the sorted cells were reintroduced into the ankle joint of BALB/c mice on day 7 of IL-23OE, coinciding with the decrease in CD200^+^ fibroblasts to increase the pool of migrated cells in the joint. The level of inflammation was assessed histologically and by MRI on day 14. The adoptive transfer resulted in an increased arthritis severity at this early time point (**Figure ED5c**), highlighting the critical pro-inflammatory role of CD2^+^ MHC-II^+^ CCR2^+^ myeloid precursors in an arthritis-conductive environment. Second, we used IL-23OE in the arthritis-resistant C57BL/6 inbred strain and intervened every second day with an anti-CD200 treatment (OX-90) to block the suppression of skin-derived precursors by the resident CD200^+^ fibroblasts. After treatment with blocking anti-CD200, the initially arthritis-resistant animals developed joint inflammation on day 21 (**Figure 4l**) supporting the critical gate-keeping role of CD200^+^ fibroblasts interacting with migrating CD2^+^ MHC-II^+^ CCR2^+^ myeloid precursors. Third, we induced skin psoriasis in BALB/c mice using imiquimod, which is a locally restricted model involving the topical application of imiquimod to the skin to test our hypothesis in a model that is independent of systemic IL-23OE. Using this model, we observed an increase in CD2^+^ MHC-II^+^ CCR2^+^ myeloid precursors in the joints on day 7, but no decrease in CD200^+^ fibroblasts (**Figure ED5d**). Again, we intervened every second day with an anti-CD200 treatment (OX-90) to block the suppression of skin-derived precursors by the resident CD200^+^ fibroblasts. In line with our second approach, on day 21, only animals that received the blocking antibody developed joint inflammation (**Figure ED5e**). Fourth, we used IL-23OE in the arthritis-susceptible BALB/c inbred strain and intervened every five days with an agonistic anti-CD200R1 treatment (OX-110) starting on day 7 to restore the suppression of skin-derived precursors by the resident CD200^+^ fibroblasts. After treatment with agonistic anti-CD200R1, the arthritis was significantly reduced on day 21 (**Figure 4m**) supporting the critical anti-inflammatory signalling mediated by CD200.

### Myeloid precursors activate local T cell in the synovia

To challenge our observations made in mice, we firstly inspected our human scRNAseq database. In the human synovial scRNAseq dataset, we also found substantial expression of *CD200R1* on the CD2^+^ MHC-II^+^ CCR2^+^ skin-derived joint-invading myeloid precursors (**Figure 5a**). Similarly, flow cytometric analysis of PBMCs from PsA patients showed an equally high expression of CD200R1 on the CD2^+^ MHC-II^+^ CCR2^+^ skin-derived joint-invading myeloid precursors (**Figures 5b, ED6a** and **ED6b**).

**Figure 5:**
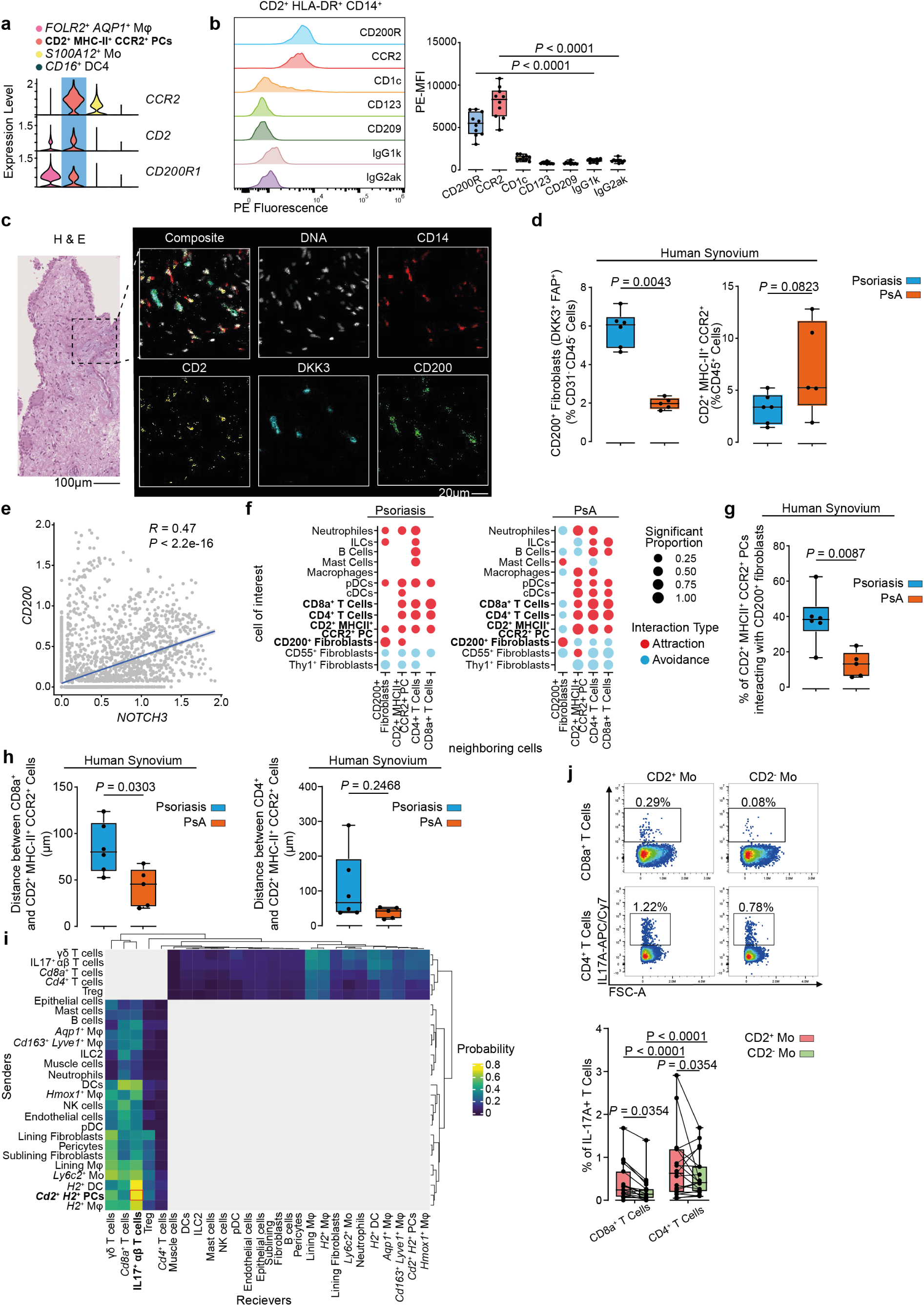
CD200-CD200R1 controls onset of joint inflammation. (**a**) Violin plots of ALRA-imputed gene expression among the identified clusters of human synovial myeloid cells from Figure 3a. Abbreviation: precursors (PCs). (**b**) Representative histogram and quantification of the median fluorescence intensity (MFI) of marker expression on CD2^+^ HLA-DR^+^ CD14^+^ circulating monocytes in PBMCs of psoriatic arthritis (PsA) patients; *N* = 10; *P*-values were calculated by Mann-Whitney *U*-test between target markers and isotype controls. (**c**) Representative micrographs of haematoxylin/eosin (H & E)-stained sections of human synovial tissue and selected imaging mass cytometry (IMC)-stained markers (**d**) IMC-based quantification of CD200^+^ fibroblasts and of CD2^+^ MHC-II^+^ CCR2^+^ myeloid precursors in synovial tissue of psoriasis and early PsA patients. Graphs show median, quartiles and min-max.; *N* = 6 and *N* = 5, respectively; *P*-values were calculated by Mann-Whitney *U*-test. (**e**) *CD200* expression correlated to *NOTCH3* expression in synovial fibroblasts (grey dots) from **Figure ED4a**. Blue line shows the linear fit, and the Gray region shows the confidence interval. Two-sided Pearson correlation and Benjamini and Hochberg (BH) adjusted *P*-value are shown. (**f**) Pairwise interaction test between CD2^+^ MHC-II^+^ CCR2^+^ myeloid PCs, CD200^+^ fibroblasts and T cells with the surrounding immune and mesenchymal cells in the IMC dataset of synovial tissue from psoriasis (*N* = 6) and PsA (*N* = 5) patients. (**g**) Proportion of CD2^+^ MHC-II^+^ CCR2^+^ PCs interacting with CD200^+^ fibroblasts in IMC dataset of **c**. (**h**-**i**) Distance between CD8a^+^ (h) and CD4^+^ (i) T cells and CD2^+^ MHC-II^+^ CCR2^+^ myeloid PCs cells in the IMC dataset of synovial tissue from psoriasis (*N* = 6) and PsA (*N* = 5) patients. (**j**) Hierarchically clustered heatmap showing the CellChat calculated interaction probability between subclusters in the scRNAseq dataset of Kaede^tg^ animals. Grey tiles are excluded interactions. (**k**) Representative flow cytometry plots for the IL-17A expression in CD8^+^ and CD4^+^ T cells after co-culture with CD2^+^ or CD2^-^ MHC-II^+^ CCR2^+^ myeloid cells. Graphs show quantification of IL-17A expression, correspondingly, as mean, quartiles and min-max; *N* = 19; *P*-values were calculated by two-way ANOVA with Tukeýs *post hoc* test.

We then used imaging mass cytometry (IMC) to detect CD2^+^ MHC-II^+^ CCR2^+^ myeloid precursors and CD200^+^ (DKK3^+^ FAP^+^ CD45^-^ CD31^-^) fibroblasts in the synovial tissues of early PsA patients, who just had developed arthritis, and psoriasis patients without arthritis (**Figure 5c**). Consistent with our previous observations in mice and humans, we observed an increase in CD2^+^ MHC-II^+^ CCR2^+^ myeloid precursors and a significant decrease in CD200^+^ fibroblasts in the synovial tissue during the onset of arthritis in humans (**Figure 5d**).

Since close cell-cell contact is essential for effective CD200 signalling, we localised CD200^+^ fibroblasts in the joint. To this end, we analysed the expression of *CD200* in relation to *NOTCH3* in fibroblasts, as *NOTCH3* has been described to form a decreasing gradient within the synovium, from the perivascular space to the lining layer ^35^. We found a significant correlation between *NOTCH3* and *CD200* in synovial tissue (**Figure 5e**), suggesting that CD200^+^ fibroblasts reside within the perivascular space and distal of the lining layer. Similarly, when we plotted the CD200 signal against the CD55 signal in fibroblasts in the IMC dataset, we found a significant negative correlation (**Figure ED6c**). In this dataset, we also examined the spatial cellular neighbourhood of CD2^+^ MHC-II^+^ CCR2^+^ myeloid precursors and CD200^+^ fibroblasts and quantified their interaction strength. While both cell populations showed a significant attraction in the non-inflamed synovia of psoriasis patients, in the inflamed synovia of PsA patients, both cell populations showed a significant avoidance (**Figure 5f**). Consequently, there was a significant reduction in the percentage of CD2^+^ MHC-II^+^ CCR2^+^ myeloid precursors interacting with CD200^+^ fibroblasts in PsA patients compared to psoriasis (**Figure 5g**). These analyses show that the priming of skin-derived recirculating CD200R1-expressing CD2^+^ MHC-II^+^ CCR2^+^ myeloid precursors by CD200^+^ fibroblasts occurs immediately upon entry into the synovial tissue and before development into subsidiary macrophage subsets. The interaction between the two cell populations is reduced in PsA due to the reduced numbers of CD200^+^ fibroblasts.

CD4^+^ and CD8^+^ T cells play a significant role in the pathogenesis of PsA ^40, 41^. To investigate the pathogenic potential of CD2^+^ MHC-II^+^ CCR2^+^ myeloid precursors, we conducted an interaction analysis with T cells. Both the CD4^+^ and CD8^+^ T cell populations demonstrated attraction to CD2^+^ MHC-II^+^ CCR2^+^ myeloid precursors in the non-inflamed synovia of psoriasis patients and the inflamed synovia of PsA patients (**Figure 5f**). Notably, we found a significant reduction in the distance between CD2^+^ MHC-II^+^ CCR2^+^ myeloid precursors and particularly CD8^+^ T cells in PsA patients, indicating enhanced interaction (**Figure 5h**). Furthermore, analysis of the scRNAseq dataset from Kaede^tg^ animals revealed that CD2^+^ MHC-II^+^ CCR2^+^ myeloid precursors exhibited the highest UCell scores for ’positive regulation of T Cell chemotaxis’ (GO:0010820) (**Figure ED6d**), and had one of the highest interaction probabilities with IL-17^+^ T cells (**Figures 5h** and **ED6e,f**). To further evaluate this interaction, we sorted CD4^+^ and CD8^+^ T cells from PBMCs of PsA patients and co-cultured them with either CD2^+^ or CD2^-^ MHC-II^+^ CCR2^+^ myeloid cells, which were correspondingly isolated from the same patients. Only CD2^+^ MHC-II^+^ CCR2^+^ myeloid precursors positively influenced IL-17 expression in both T cell subsets, contrasting with the CD2^-^ MHC-II^+^ CCR2^+^ myeloid cells (**Figure 5j**). In conclusion, the greater spatial proximity and functional interaction between myeloid precursors and CD8⁺ T cells in the synovium of PsA patients likely induces IL-17 expression in local T cells. This finding corroborates the concept that skin-derived CD2^+^ MHC-II^+^ CCR2^+^ myeloid precursors play a key role in the pathogenesis of PsA.

## DISCUSSION

Herein, we show a two-step process that orchestrates the progression from psoriatic skin to joint disease. Psoriasis induced a population of pro-inflammatory CD2^+^ MHC-II^+^ CCR2^+^ myeloid precursors to migrate from the skin to the joints, but their migration was not sufficient to induce arthritis. Rather, the local microenvironment determined the effects of CD2^+^ MHC-II^+^ CCR2^+^ myeloid precursors in the joints, with CD200^+^ fibroblasts acting as gatekeepers for arthritis by engaging the checkpoint receptor CD200R1 on migrating CD2^+^ MHC-II^+^ CCR2^+^ myeloid precursors. As a result, intervention with an anti-CD200 antibody polarized CD2^+^ MHC-II^+^ CCR2^+^ myeloid precursors towards a pro-inflammatory phenotype *in vitro* and allowed the *in vivo* spreading of psoriatic disease from the skin to the joints even in initially arthritis-resistant mice.

Our finding of myeloid precursor migration between skin and joint sheds light on a previously unexplored aspect of the pathogenesis of psoriatic disease and inflammation in general. Tracking cell migration between tissues in humans is technically limited due to a lack of labelling options. The introduction of next-generation single cell sequencing has focused attention on lymphocytes, whose propagation between tissues can be tracked due to their unique clonal expansion with inherited receptor rearrangements. For example, shared T cell repertoires have been identified between synovial fluid and blood T cells in patients with PsA and spondylarthritis ^42^ or between skin and blood in healthy individuals ^9^. In addition, skin-resident T cells express high levels of cutaneous lymphocyte-associated antigen (CLA), allowing them to be tracked in the peripheral blood. However, it has been described that CLA^+^ T cells preferentially (re)migrate to the skin of PsA patients rather than the synovial tissues ^43^. As these methods do not capture the full spectrum of leukocytes, we first took a translational approach to investigate the migration of any leukocyte population from the skin in mouse models of psoriasis with or without concomitant development of arthritis, using stable photoconversion of a fluorescent reporter as a robust labelling method. We found that the majority of skin-derived cells within the joint belonged to distinct myeloid populations, most of which are typically characterized as tissue-resident and non-migratory ^17^. However, we identified a migrating myeloid precursor subset, CD2⁺ MHC-II⁺ CCR2⁺, which gives rise to the broader pool of skin-derived myeloid cells observed in the joint. Secondly, by extrapolating our results from mice to humans, we identified conserved CD2^+^ MHC-II^+^ CCR2^+^ myeloid precursors in the skin and synovial tissue across species. To test whether these precursors migrate between skin and joint, we used the MAESTER method ^30^ to track mitochondrial variants based on mRNA. This method relies on identifying somatic mutations, the rate of which is 10- to 100-fold higher in the mitochondrial genome than in the nuclear genome ^44^. MAESTER provides a significant advantage over originally mtDNA-based methods, resulting in a full mRNA library allowing for its detailed analysis. Using mitochondrial variant tracking, we could corroborate the skin-joint-axis in humans. Furthermore, we showed an increased abundance of precursors in psoriasis and PsA patients compared to healthy controls suggests that the migration of these precursors is an event that occurs independently of joint involvement but constitutes a pre-requisite to spread inflammation from the skin to the joints. The consistency of our results across species highlights the conserved nature of the migration of myeloid cells in the context of the skin-joint axis in psoriatic disease. This opens up new perspectives for the development of novel diagnostic and therapeutic strategies targeting immune cell trafficking in psoriasis disease even before development of arthritis.

Our study demonstrated that the development of arthritis is induced by the differentiation of CD2^+^ MHC-II^+^ CCR2^+^ myeloid precursors into highly pro-inflammatory mononuclear phagocytes. Using computational analyses such as pseudo-time, RNA velocity and interactome, we uncovered the dynamic behaviour of CD2^+^ MHC-II^+^ CCR2^+^ myeloid precursors after entering the synovial tissue, which is under tight control of the articular microenvironment. We found that synovial fibroblasts, particularly CD200^+^ fibroblasts, played a crucial role in modulating precursor activation and differentiation by activating the checkpoint receptor CD200R1 on migrating CD2^+^ MHC-II^+^ CCR2^+^ myeloid precursors after entry into the joint, thereby exerting a suppressive effect on inflammation. Conversely, blocking CD200 in different arthritis-resistant models resulted in arthritis, highlighting the inflammatory potential of skin-derived CD2^+^ MHC-II^+^ CCR2^+^ myeloid precursors. Additionally, we demonstrated that CD2^+^ MHC-II^+^ CCR2^+^ myeloid precursors are highly effective at inducing IL-17 expression in T cells. Interestingly, we did not find evidence of a cutaneous priming of the CD2^+^ MHC-II^+^ CCR2^+^ myeloid precursors, emphasising the influence of the synovial microenvironment on disease progression from the skin to the joint. We have recently identified a CD200^+^ fibroblast subtype that contributes to resolution of inflammation in arthritis ^34^ and have now demonstrated its involvement in suppressing the development of arthritis and pinpointing the CD200-CD200R1 axis as a promising target for preventive medicine.

In conclusion, our study provides first insights of the cellular and molecular mechanisms involved in the spreading of psoriatic skin inflammation to the joints. Similar as in tumour metastasis, this process requires a migrating cell population and a microenvironment that supports the activity of the migrated cells at their target tissue. By elucidating the role of CD2^+^ MHC-II^+^ CCR2^+^ myeloid precursors, synovial fibroblasts and the CD200-CD200R1 axis, we have identified several components of this process that represent potential new targets for therapeutic intervention at various stages of disease development. The identification of cellular key players and signalling pathways opens up exciting opportunities for the improvement of patient outcomes and advances in personalised medicine in the treatment of this debilitating disease.

## Acknowledgements

We thank Stefan Uderhardt and Steffen Jung for critical discussion of the data. We thank J. Friedrich, M. Rose, C. Pfaff, J. Tu, B. Happich and K-T. Yang for excellent technical assistance. We thank U. Appelt and M. Mroz from the FAU ‘Core Unit für Zellsortierung und Immunomonitoring’ for cell sorting. We thank I. Gadjalova, from the ‘Core Facility Cell Analysis’ of TranslaTUM - Central Institute for Translational Cancer Research at the Technical University of Munich for providing ImageStream imaging cytometry. We acknowledge the FAU NGS core facility for sequencing. We acknowledge the Optical Imaging Centre Erlangen (OICE) for microscopy. We acknowledge the Preclinical Imaging Platform Erlangen (PIPE) for providing small animal MRI. The IMC analysis was supported by the de.NBI Cloud within the German Network for Bioinformatics Infrastructure (de.NBI) and ELIXIR-DE (Forschungszentrum Jülich and W-de.NBI-001, W-de.NBI-004, W-de.NBI-008, W-de.NBI-010, W-de.NBI-013, W-de.NBI-014, W-de.NBI-016, W-de.NBI-022). The work was supported by the German Research Foundation (DFG) to A.R. (project numbers: 320379231 (RA 2506/4-1, RA 2506/4-2), 423477573 (RA 2506/6-1), 493624887 (RA 2506/7-1)), to M.G.R. (project number: 493624887 Clinician Scientist Program NOTICE) and to A.S. (project number: 422621469 (SO 1735/2-1)); CRC1181 (project number: 261193037) to G.S. (projects A01/Z03) and A.R. (project C06); TRR369 (project number: 501752319) to G.S.(projects C06/Z01) and A.R. (project A04); Gottfried Wilhelm Leibniz Prize 2023 to G.S; Major research instrumentation funding for light sheet microscopy, small animal MRI and CyTOF (DFG project numbers 391371888, 456565718 and 424726560, respectively). The work was supported by the European Research Council (project number 853508 - BARRIER BREAK) to A.R. and (project number 810316 - 4-D nanoScope) to G.S. The work was supported by the Federal Ministry of Education and Research (BMBF, 01EC1903A - MASCARA). This project has received funding from the Innovative Medicines Initiative 2 Joint Undertaking (JU) under grant agreement No 101007757 (HIPPOCRATES) and No 777357 (RTCure). The JU receives support from the European Union’s Horizon 2020 research and innovation program and EFPIA. Any dissemination of results must indicate that it reflects only the author’s view and that the JU is not responsible for any use that may be made of the information it contains. The work was supported by the PARTNER Fellowship program to M.G.R., S.A. and M.A.D.A. The work was supported by Novartis Pharma to A.R. The work was supported by the Interdisciplinary Centre for Clinical Research (IZKF) Erlangen (D034 to A.R., P049 and J106 to M.G.R., J107 to S.R.). The present work was performed in partial fulfilment of the requirements for obtaining the degree Dr. rer. biol. hum. at the FAU Erlangen-Nürnberg.

## Author contributions

Lead contact: Further information and requests for resources and reagents should be directed to and will be fulfilled by the lead contact, Andreas Ramming.

These authors contributed equally: MGR, HM These authors share senior authorship: SR, AR Design of the study: MGR, HM, SR, AR

Acquisition of data: MGR, HM, MRA, SA, VF, KH, RD, PR, CX, RM, ZW, ARR, CAG, AS, ML, HL, JDC, SR

Interpretation of data: MGR, HM, SA, VF, KH, RD, PR, ZW, CX, CAG, TB, GS, SR, AR

Support of material: SA, CG, UF, DJV, FC, JR, MS, JHWD, MSKS, MM, ABE, APC, OD, CO, JDC, MADA, GS

Manuscript preparation: MGR, HM, VF, GS, SR, AR

## Declaration of interest

Peter Rhein is employed by Cytek Biosciences, the maker of the Amnis brand ImageStream®, which was used in this study. Oliver Distler has/had consultancy relationship with and/or has received research funding from and/or has served as a speaker for the following companies in the area of rheumatology in the last three calendar years: 4P-Pharma, Abbvie, Acepodia, Aera, AnaMar, Anaveon, Argenx, AstraZeneca, Boehringer Ingelheim, BMS, Calluna, Cantargia AB, CSL Behring, EMD Serono, Galderma, Galapagos, Gossamer, Hemetron, Innovaderm, Janssen, Lilly, MSD Merck, Nkarta, Novartis, Oorja Bio, Orion, Pilan, Prometheus, Quell, Redxpharma, Scleroderma Research Foundation, Sumitomo, Topadur, UCB and Umlaut bio. Patent issued “mir-29 for the treatment of systemic sclerosis” (US8247389, EP2331143). Co-founder of CITUS AG. The rest of the authors declare no competing interests.

## Extended Data Figures

**Figure ED1:**
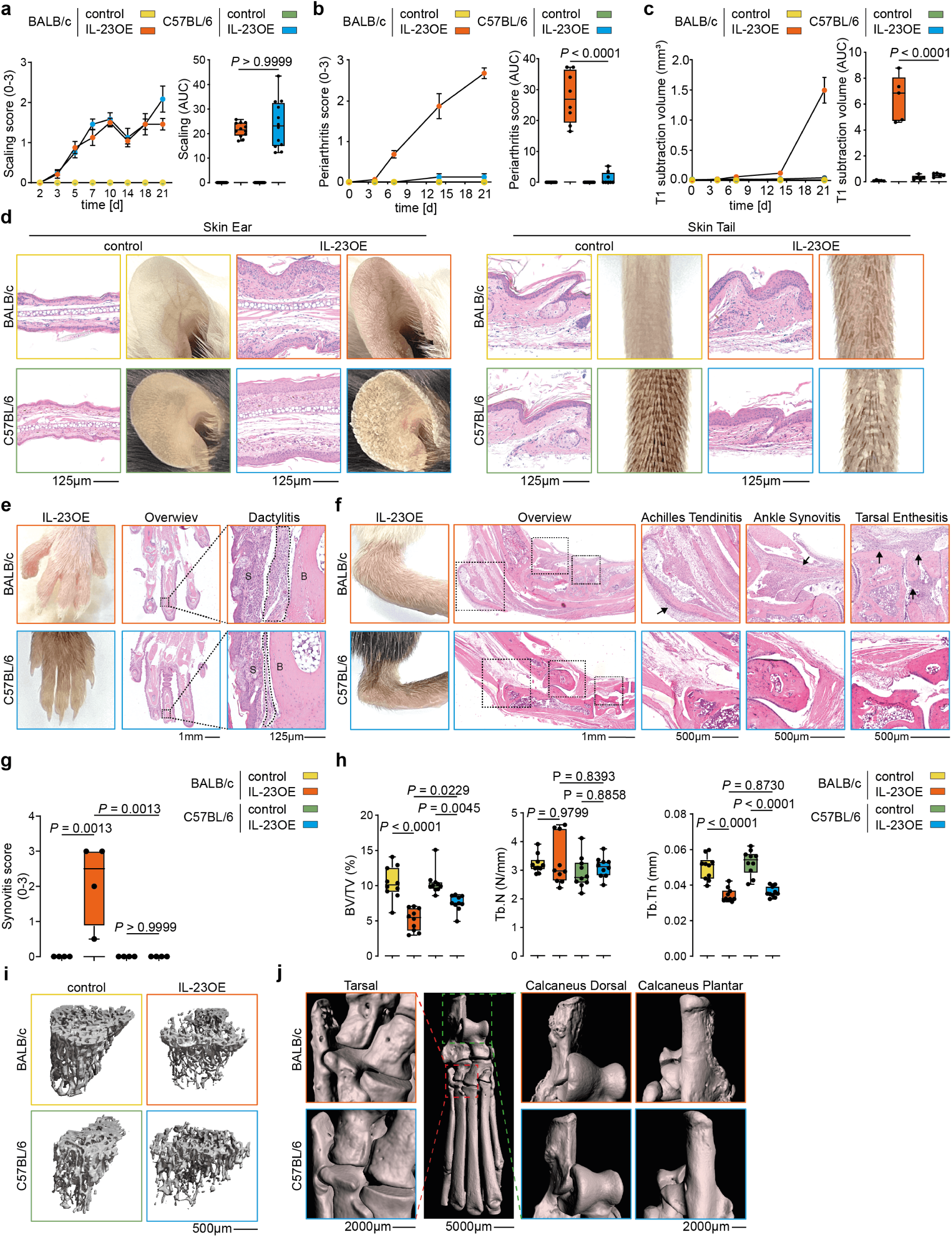
Assessment of psoriasis and arthritis over time. (**a**) Macroscopic evaluation of skin-scaling in BALB/c and C57BL/6 mice with and without IL-23 overexpression (OE) for 21 days (left). Quantification of total scaling as area under the curve (right). Graphs show mean and standard error of the mean (left) and median, quartiles and min-max (right); *N* = 12 per condition. *P*-values were calculated by one-way ANOVA with Tukeýs *post hoc* test. (**b**) Histological inflammation score of the ankle area based on haematoxylin & eosin (H & E) stained sections from BALB/c and C57BL/6 mice with and without IL-23OE for 21 days (left). Quantification of arthritis as area under the curve (right). Graphs show mean and standard error of the mean (left) and median, quartiles and min-max (right); *N* = 8 per time point and condition. *P*-values were calculated by one-way ANOVA with Tukeýs *post hoc* test. (**c**) MRI ankle measurements over time in BALB/c and C57BL/6 mice with and without IL-23OE for 21 days (left). Quantification of arthritis as area under the curve (right). Graphs show mean and standard error of the mean (left) and median, quartiles and min-max (right); *N* = 5 per condition. *P*-values were calculated by one-way ANOVA with Tukeýs test. (**d**) Representative micrographs of H & E-stained skin sections from ear and tail of BALB/c and C57BL/6 mice at day 21 with and without IL-23 OE together with representative pictures of the same regions; (**e**) Representative micrographs of H & E-stained paw sections and pictures of distal hind paws of BALB/c and C57BL/6 mice at day 21 after IL-23 OE. Dashed lines highlight dactylitis. Abbreviations: S = skin, B = bone. (**f**) Representative micrographs of H & E-stained ankle sections and representative pictures of ankle joints of BALB/c and C57BL/6 at day 21 with IL-23OE. Dashed squares in the overview identify the areas of corresponding magnifications. Black arrows indicate sites of inflammation. (**g**) Histological inflammation score of the synovia in ankle and tarsal joints based on H & E-stained sections from BALB/c and C57BL/6 mice with and without IL-23OE for 21 days. (**h**) Micro-CT quantification of tibia from BALB/c and C57BL/6 mice at day 21 with IL-23 OE. Abbreviations: BV/TV = bone volume/total volume; Tb. N = trabecular number; Tb. Th = trabecular thickness; *N* = 10 per condition; *P*-values were calculated by one-way ANOVA with Tukeýs *post hoc* test. (**i**) Representative micro-CT images of the tibial trabecular network of from BALB/c and C57BL/6 mice at day 21 with or without IL-23 OE. (**j**) Representative micro-CT images from hind paws of BALB/c and C57BL/6 mice at day 21 with or without IL-23OE. Red and green dashed squares indicating the corresponding magnification of metatarsal bone and calcaneus.

**Figure ED2:**
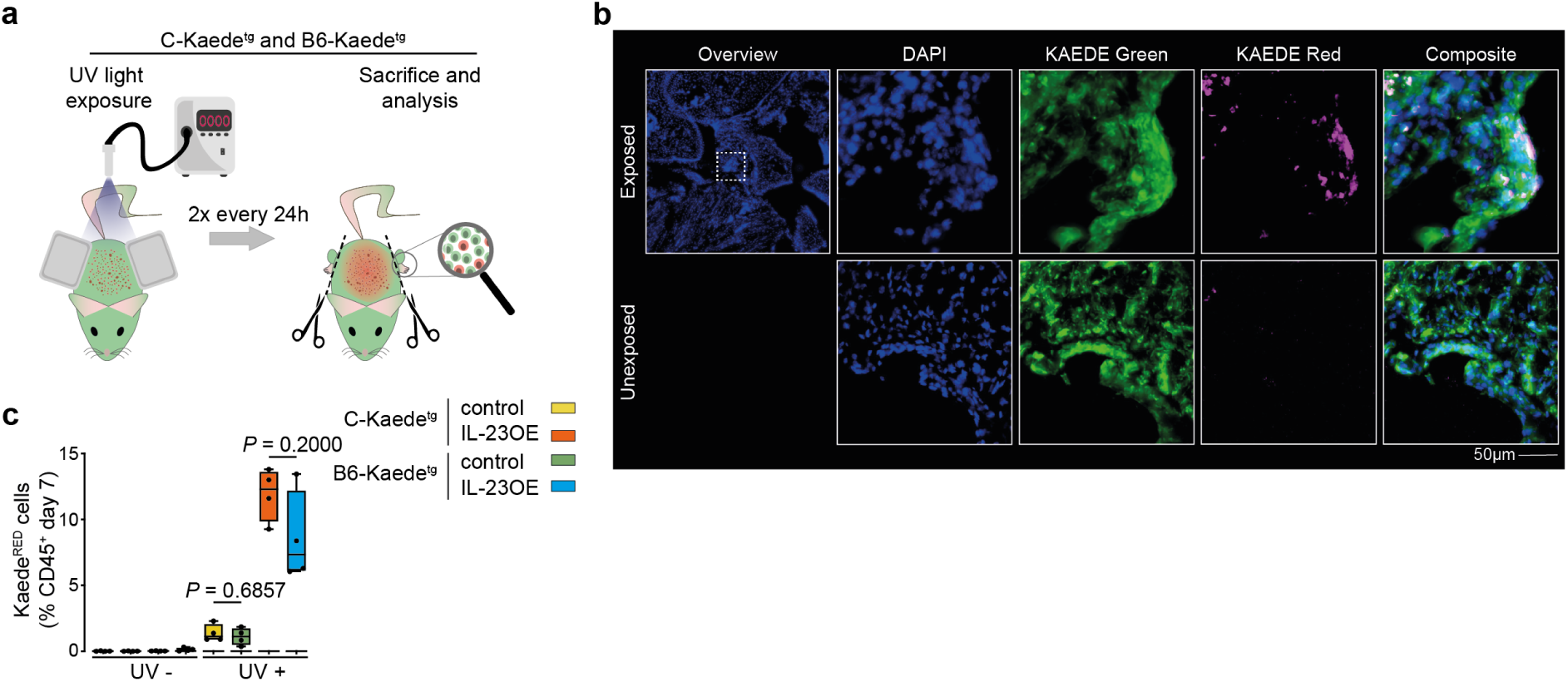
Validation of photoconversion. (**a**) Graphical cartoon explaining skin-photoconversion of the Kaede^tg^ mice upon ultraviolet (UV) exposure. (**b**) Representative micrographs of immunofluorescence microscopy of synovial tissue from hind paw joints of photoconverted (exposed) or not photoconverted (unexposed) Kaede^tg^ at day 7 with IL-23 overexpression (IL-23 OE). (**c**) Quantification of Kaede^RED^ skin-derived cells in the joints at day 7. Graph shows median, quartiles and min-max; *N* = 4 per condition; *P*-values were calculated by Mann-Whitney U-test.

**Figure ED3:**
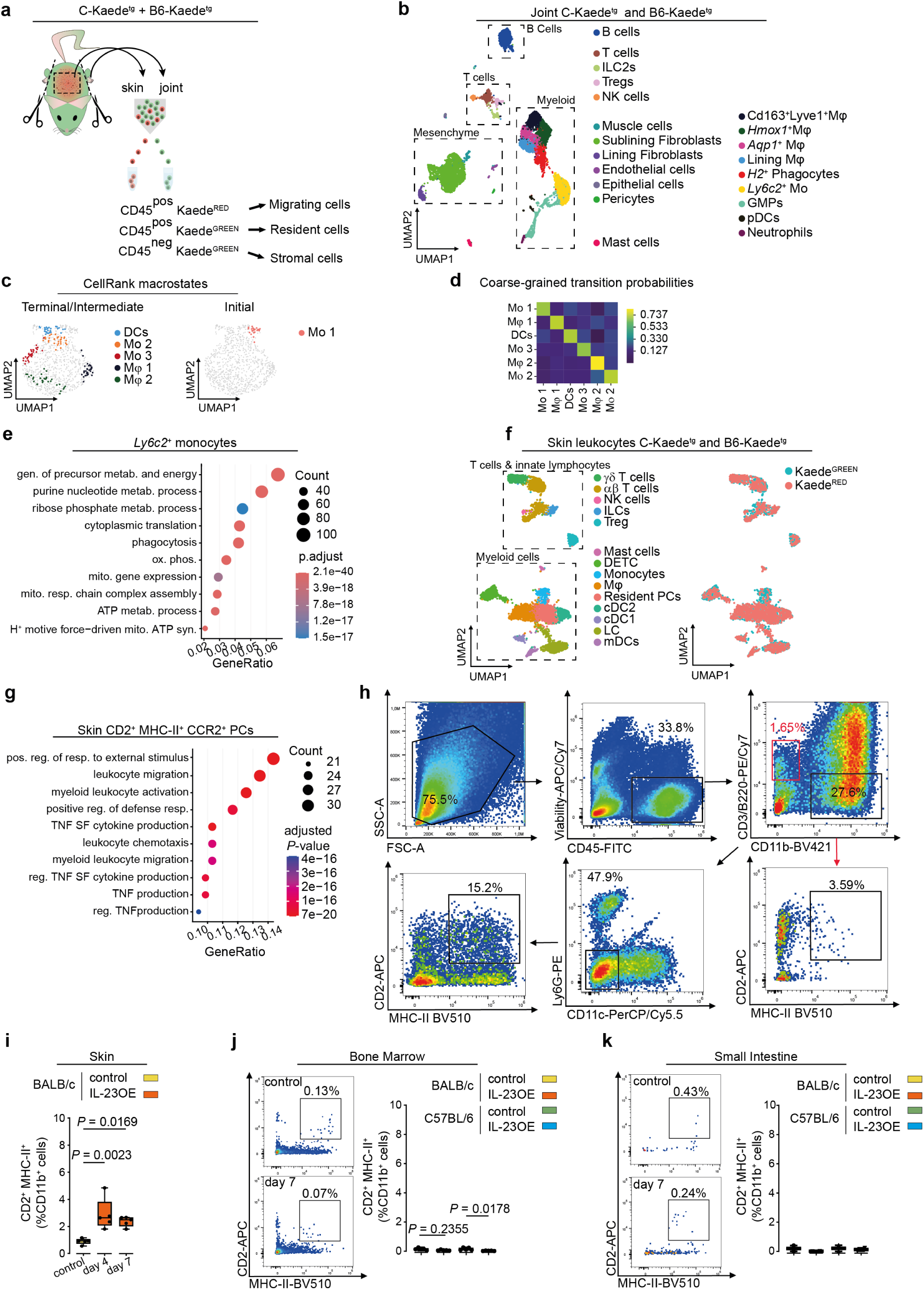
Mouse single-cell landscape in the joints and the skin at day 7 after IL-23OE. (a) Graphical cartoon showing the strategy to cell sort target populations used for scRNAseq. Target populations were individually hash-tagged before generating the scRNAseq library. (**b**) UMAP plot of the hash-tag identified ankle joint cells in the scRNAseq dataset from Kaede^tg^ mice (BALB/c and C57BL/6 strains) on day 7 of the IL-23OE model. Detailed annotation with Seurat-identified clusters is shown. (**c**) CellRank identification of terminal (left) and initial states (right) in the Kaede^RED^ myeloid cells. (**d**) Coarse-grained matrix of transition probabilities between states calculated by CellRank. (**e**) Significantly enriched GO-BP terms in the *Ly6c2*^+^ monocyte cluster, when compared to all other cell types in the joint. Selection criteria: BH-adjusted *P*-value < 0.05. (**f**) UMAP plot of the hash-tag identified skin leukocytes in the scRNAseq dataset from Kaede^tg^ mice (BALB/c and C57BL/6 strains) on day 7 of the IL-23OE model. Annotated Seurat identified clusters are shown (left). Kaede^GREEN^ and Kaede^RED^ cells are highlighted (right). (**g**) Significantly enriched GO-BP terms in the skin CD2^+^ MHC-II^+^ CCR2^+^ myeloid precursors (PCs) cluster, when compared to other cell types. Selection criteria: BH-adjusted *P*-value < 0.05. (**h**) Representative flow cytometry plots showing the gating strategy for the identification of CD2^+^ MHC-II^+^ CCR2^+^ skin-derived precursors in the joint. Red gate on T and B lymphocytes as internal control for CD2 and MHC-II. (**i**) Quantification of CD2^+^ MHC-II^+^ myeloid cells in the skin of BALB/c animals after IL-23OE over time. Graph shows median, quartiles and min-max.; *N* = 5, per timepoint and condition; *P*-values were calculated by one-way ANOVA with Tukeýs *post hoc* test. (**j**) Representative flow cytometry plots showing the gating of CD2^+^ MHC-II^+^ skin-derived myeloid precursors in the bone marrow (left). Quantification of CD2^+^ MHC-II^+^ skin-derived myeloid precursors in the bone marrow after IL-23OE on day 7 in BALB/c and C57BL/6 mice (right). Graph shows median, quartiles and min-max.; *N* = 5, per condition; *P*-values were calculated by unpaired Man-Whitney *U*-test. (**k**) Representative flow cytometry plots showing the gating of CD2^+^ MHC-II^+^ skin-derived myeloid precursors in the small intestine (left). Quantification of CD2^+^ MHC-II^+^ skin-derived myeloid precursors in the small intestine after IL-23OE on day 7 in BALB/c and C57BL/6 mice. Graph shows median, quartiles and min-max.; *N* = 2, per condition; no *P*-values were calculated.

**Figure ED4:**
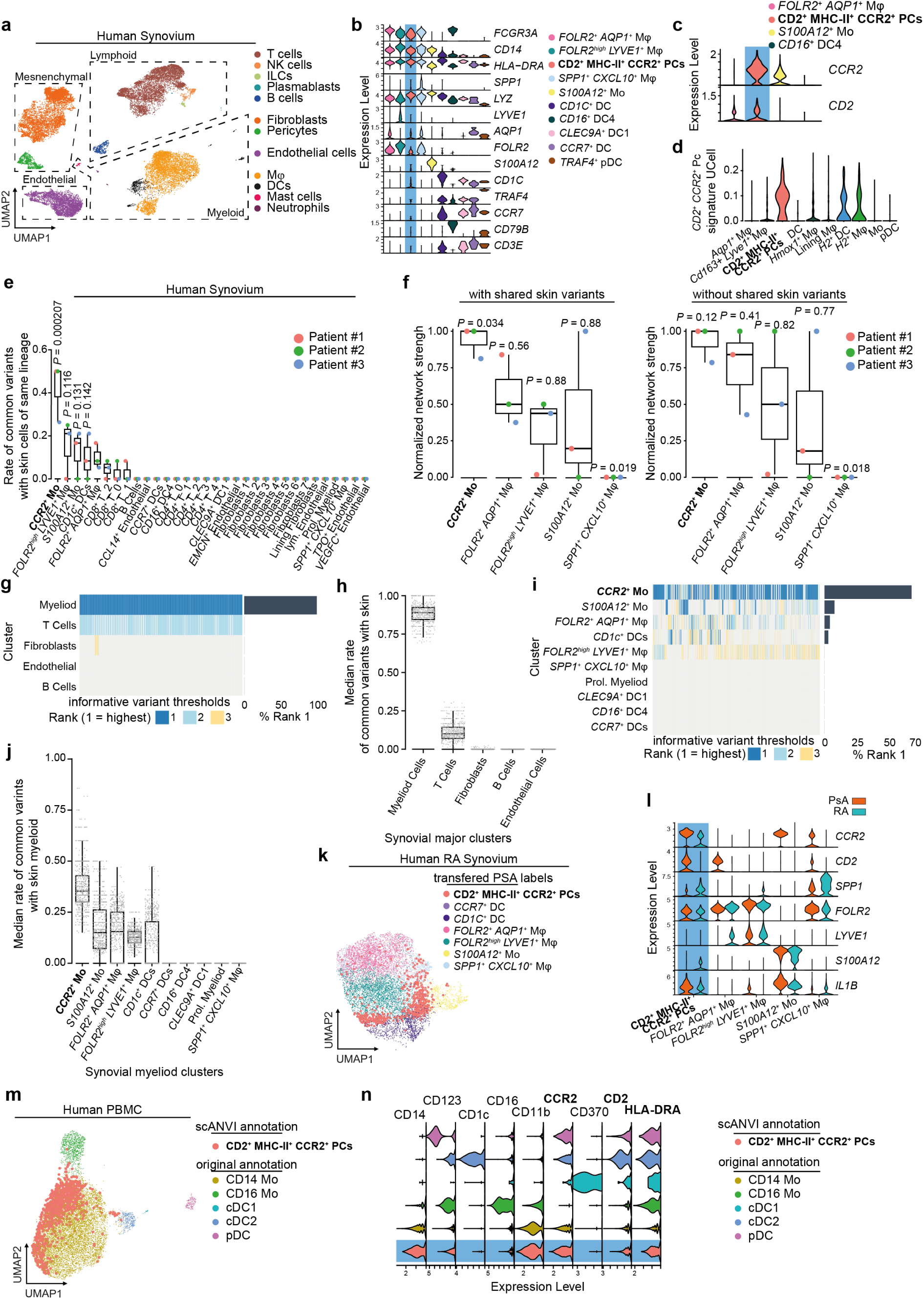
Human single cell landscape of the myeloid compartment in synovium, skin and PBMC of psoriatic disease. (**a**) UMAP plot of the identified clusters of major cell types in the scVI-integrated scRNAseq datasets of human (PsA, *N* = 5, E-MTAB-11791) and healthy synovial tissue (*N* = 3). (**b**) Violin plots of relevant marker genes based on ALRA-imputed gene expression among the identified clusters of human synovial myeloid cells from Figure 3a. Abbreviation: precursors (PCs). (**c**) Violin plots of ALRA-imputed gene expression of selectively *CD2* and *CCR2* among the identified clusters of human synovial myeloid cells from Figure 3a. (**d**) Comparison of the UCell score of the gene signature for human joint CD2^+^ MHC-II^+^ CCR2^+^ myeloid precursors in the myeloid phagocytes of the scRNAseq dataset of mouse ankle joint among dataset from Figure 2a. (**e**) Proportions of shared mitochondrial variants between synovial and skin clusters of the same lineage for all myeloid, T and B cell, fibroblast and endothelial clusters in the synovia. Graph shows median, quartiles and min-max.; *N* = 3; *P*-values were calculated by one-vs-rest Wilcoxon rank-sum test with BH correction. (**f**) Normalized network strength for the shared mitochondrial variants among subclusters of myeloid cells in the synovia, when all shared variants are included (left) or shared variants with skin are excluded (right). Graph shows median, quartiles and min-max.; *N* = 3; *P*-values were calculated by one-vs-rest Wilcoxon rank-sum test with BH correction. (**g-h**) Median rank (*N* = 3) in each iteration of synovia-skin shared variant identification among major cell clusters shown as heatmap in **g** or boxplot in **h**. (**i-j**) Median rank (*N* = 3) in each iteration of synovia-skin shared variant identification among myeloid cell clusters shown as heatmap in **i** or boxplot in **j**. (**k**) UMAP plot of a scRNAseq dataset of synovial myeloid cells of treatment naive rheumatoid arthritis patients (RA, *N* = 5, E-MTAB-8322). (**l**) Violin plots of the ALRA-imputed signature genes for each cluster identified in the RA and PsA synovial myeloid cell scRNAseq datasets. (**m**) UMAP plot of integrated CITEseq dataset of human PBMC myeloid cells. Publication annotations along with the scANVI identified clusters of CD2^+^ MHC-II^+^ CCR2^+^ myeloid precursors are shown. (**n**) Expression levels of the most relevant CITEseq antibodies among the publication clusters of human PBMC dataset along with the scANVI identified clusters of CD2^+^ MHC-II^+^ CCR2^+^ myeloid precursors.

**Figure ED5:**
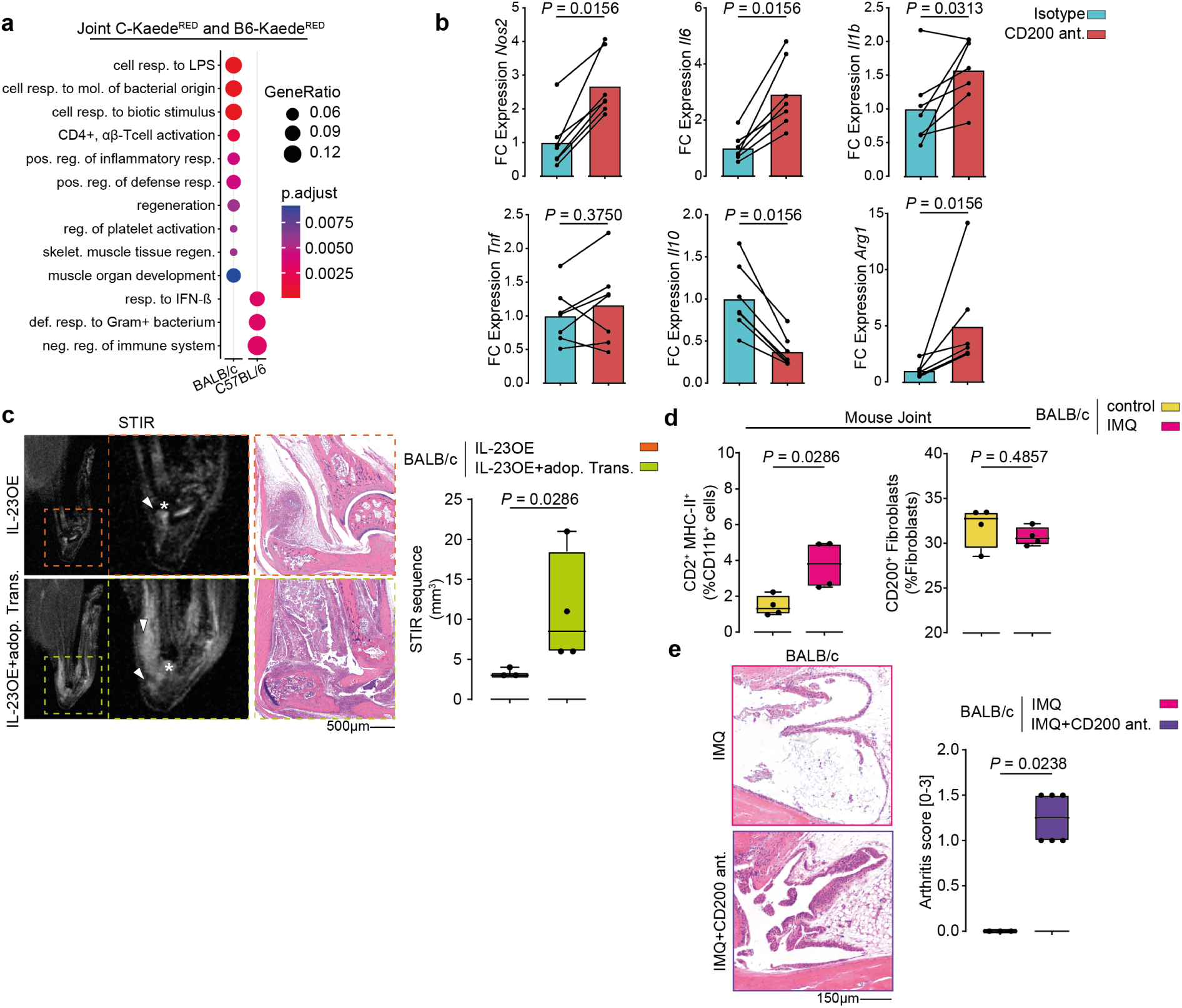
Pro-inflammatory responses in BALB/c ankle joins. Comparison of the significantly enriched GO-BP terms in the genes dynamically associated with either BALB/c or C57BL/6 strain over pseudo-time as shown in Figure 4a. Selection criteria: BH-adjusted *P*-value < 0.05. (**b**) Mean fold change (FC) gene expression in macrophage-fibroblasts co-cultures treated with CD200 antagonist (OX-90) or isotype; *N* = 7 per condition; *P*-values were calculated by Wilcoxon matched-pairs signed-rank test. (**c**) Representative images of MRI-scans and haematoxylin & eosin (H & E)-stained ankles of BALB/c mice at day 14 after IL-23 OE with or without adoptive transfer of skin-derived myeloid cells at day 7 with IL-23 OE. MRI-quantification of arthritis at day 21. Arrowheads indicate inflammation. Stars indicate the talar bone. Graph shows median, quartiles and min-max; *N* = 4 per condition; *P*-values were calculated by Mann-Whitney U-test (**d**) Quantification of CD2^+^ MHC-II^+^ CCR2^+^ skin-derived myeloid precursors and of CD200^+^ fibroblasts in the joint of BALB/c on day 7 with or without imiquimod (IMQ). Graph shows median, quartiles and min-max.; *N* = 4 per condition; *P*-values were calculated by Mann-Whitney U-test. (**e**) Representative micrographs of haematoxylin & eosin (H & E)-stained ankles of BALB/c mice on day 21 after IMQ with or without anti-CD200 antagonist (OX-90) treatment and corresponding quantification of arthritis score; *N* = 3 mice treated with IMQ and N = 6 mice with IMQ and CD200 antagonist; *P*-values were calculated by Mann-Whitney U-test.

**Figure ED6:**
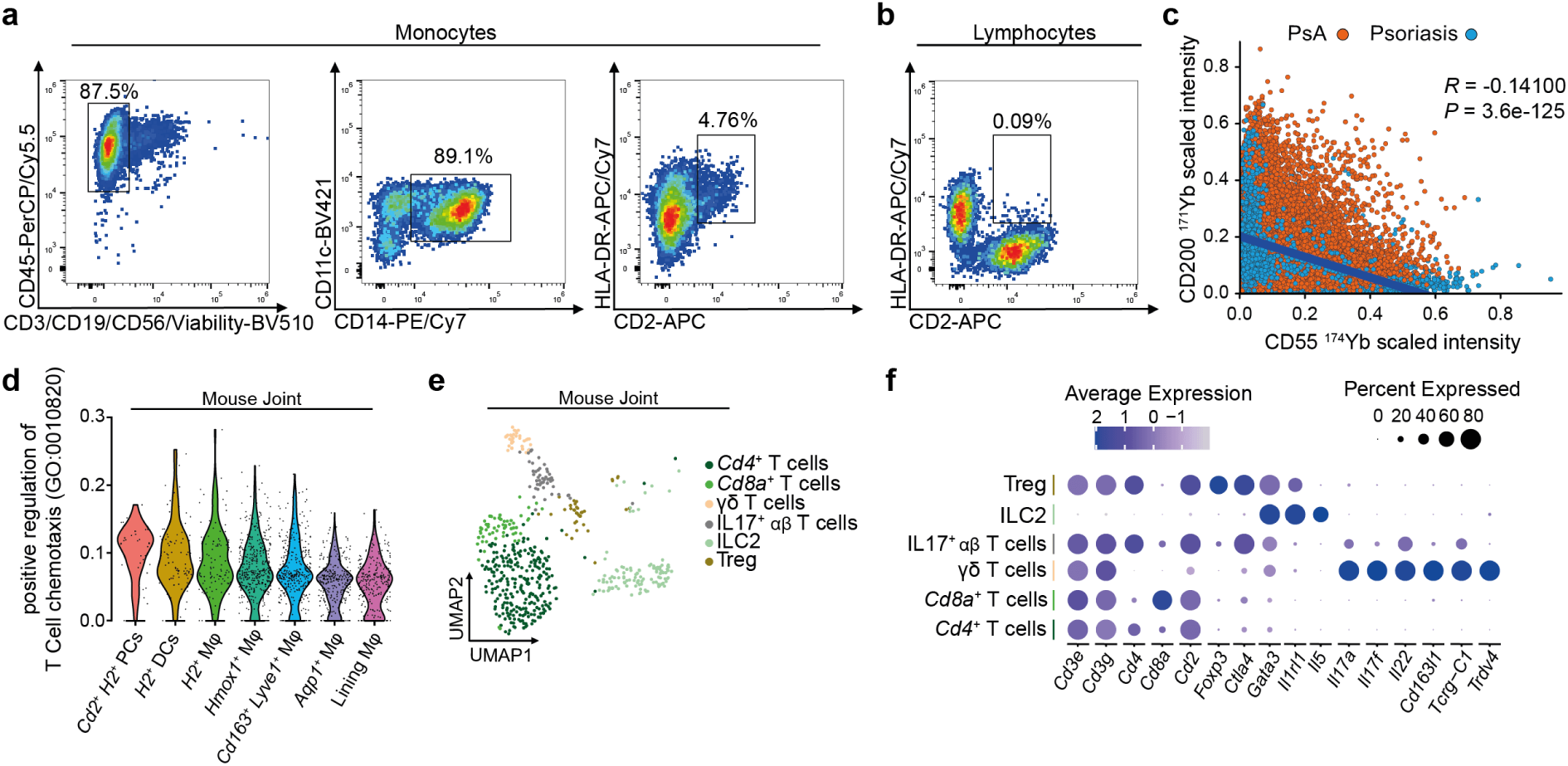
CD2^+^ MHC-II^+^ CCR2^+^ myeloid precursors trigger IL-17 expression in T cells. (**a**) Representative flow cytometry plots showing the gating strategy for the identification of CD2^+^ HLA-DR^+^ CD14^+^ circulating monocytes (pre-gated on FSC/SSC) in PBMCs of PsA patients. (**b**) Representative flow cytometry plot showing CD2^+^ HLA-DR^+^ gate on lymphocytes. (**c**) Scaled CD200 signal intensity correlated to scaled CD55 signal intensity in synovial of fibroblasts from psoriasis (yellow dots) or PsA patients (orange dots) from Figure 4i. Blue line shows the linear fit. Two-sided Pearson correlation and *P*-value are shown. (**d**) Mean fold change (FC) gene expression in macrophage-fibroblasts co-cultures treated with CD200 antagonist (OX-90) or isotype; *N* = 7 per condition; *P*-values were calculated by Wilcoxon matched-pairs signed-rank test.

## Supplementary Figures

**Supplementary Figure 1:**
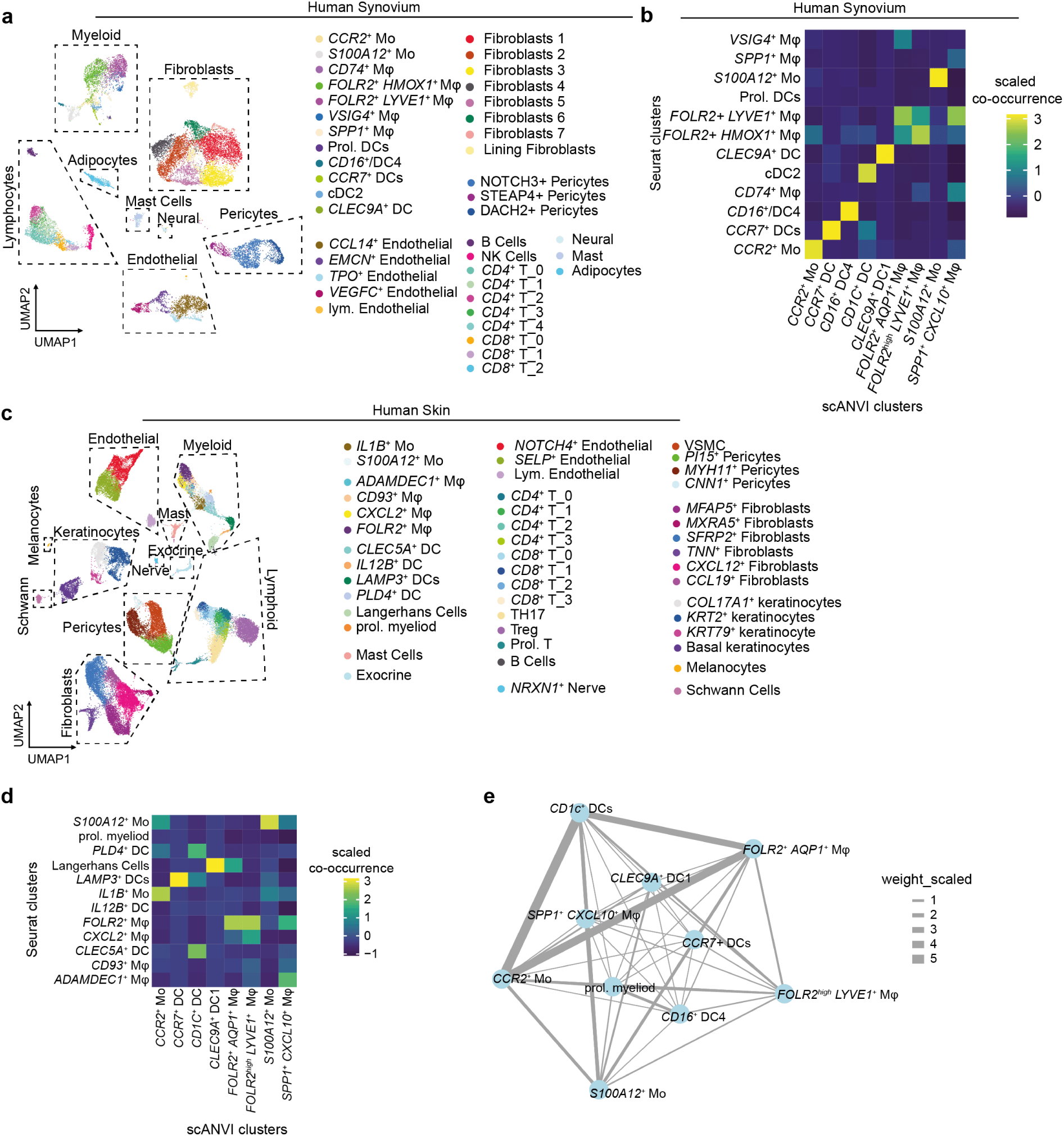
(**a**) UMAP plot of the identified clusters in the integrated dataset of synovia from psoriasis at risk and PsA patients used in the mitochondrial variant enrichment experiment. Psoriasis at risk *N* = 2, PsA *N* = 1. (**b**) Heatmap of co-occurrence of Seurat identified clusters and scANVI mapped labels in the scRNAseq datasets of synovia used in the mitochondrial enrichment analysis. scANVI reference mapping was done based on the dataset from Figure 3a. (**c**) UMAP plot of the identified clusters in the integrated dataset of skin from psoriasis at risk and PsA patients used in the mitochondrial variant enrichment experiment. Psoriasis at risk *N* = 2, PsA *N* = 1. (**d**) Heatmap of co-occurrence of Seurat identified clusters and scANVI mapped labels in the scRNAseq datasets of skin used in the mitochondrial enrichment analysis. scANVI reference mapping was done based on the dataset from Figure 3a. (**e**) Visualization of the shared variant graph among the myeloid subclusters in the synovium. Nodes represent each myeloid subcluster. Edge thickness corresponds to the scaled mean number of shared variants between two nodes.

**Supplementary Figure 2:**
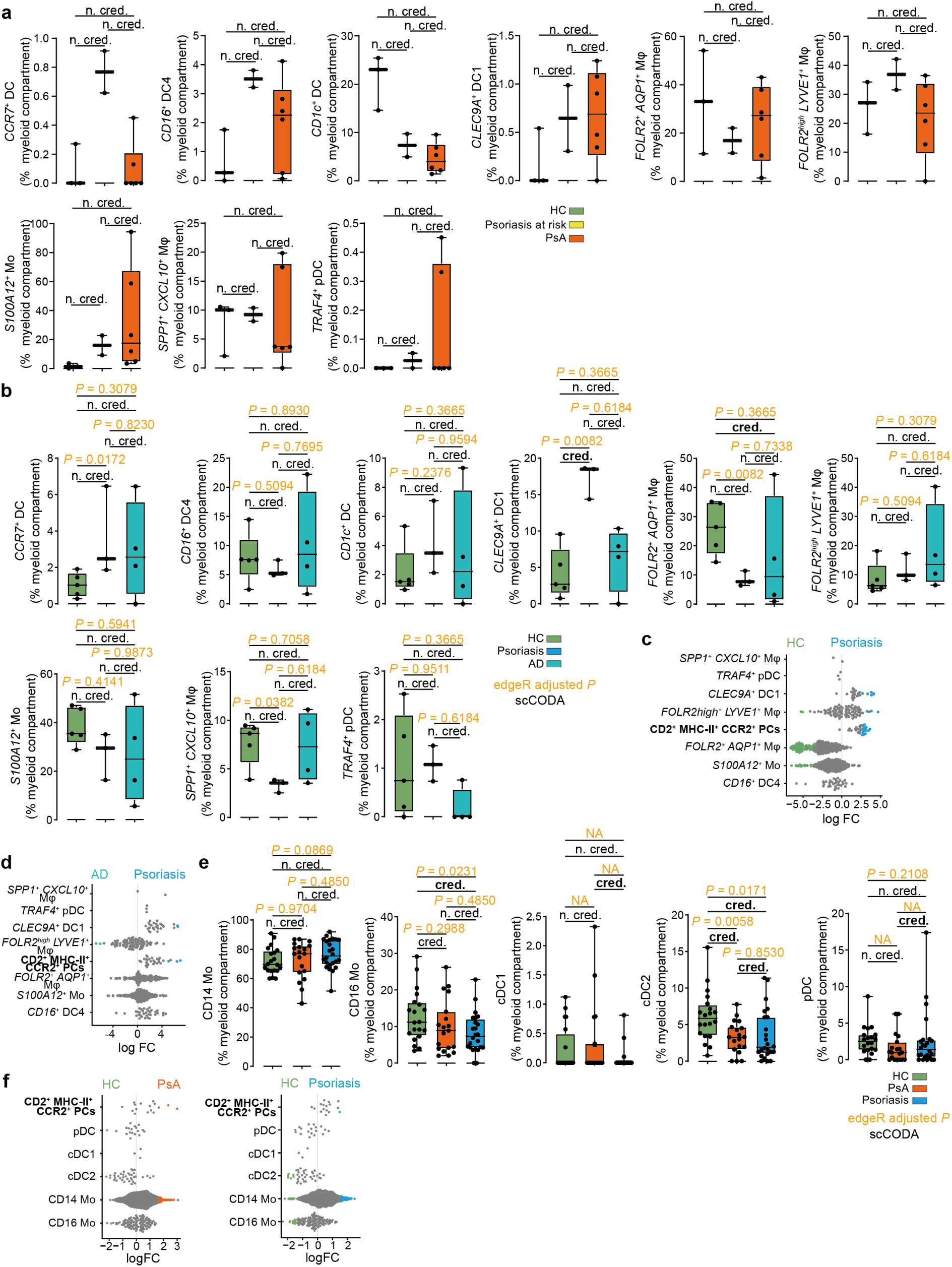
(**a**) Proportion of different myeloid cell populations in psoriasis at risk and PsA compared to healthy in the scRNAseq dataset of human synovial tissue. D2^+^ MHC-II^+^ CCR2^+^ myeloid precursors are shown in Figure 3 **f**. Graph shows median, quartiles and min-max.; psoriasis at risk *N* = 2; PsA *N* = 5, healthy subjects (HC) *N* = 3; Statistically credible (cred.) and non-credible (n. cred.) changes in abundance were identified using scCODA. (**b**) Proportion of different myeloid cell populations in healthy (HC), psoriasis and AD in scRNAseq dataset of human skin (E-MTAB-8142). CD2^+^ MHC-II^+^ CCR2^+^ myeloid precursors are shown in Figure 3 **g**. Graph shows median, quartiles and min-max.; healthy *N* = 5; psoriasis *N* = 3, AD *N* = 4; *P*-values were calculated by edgeR’s differential abundance test and corrected for multiple testing using the BH-method (orange); statistically credible (cred.) and non-credible (n. cred.) changes in abundance were identified using scCODA (black). (**c**) Differential abundance of Milo neighbourhoods in psoriatic versus healthy human skin shown as a bee swarm plot. Significantly enriched (spatial false discovery rate (FDR) < 0.05, positive fold changes (FCs), blue) or depleted (FDR < 0.05, negative FCs, green) neighbourhoods among the scANVI mapped clusters from the human synovium are highlighted. (**d**) Differential abundance of Milo neighbourhoods in atopic dermatitis (AD) versus psoriasis in the skin scRNAseq shown as a bee swarm plot. Significantly enriched (spatial false discovery rate (FDR) < 0.05, positive FCs, blue) or depleted (FDR < 0.05, negative FCs, cyan) neighbourhoods among the scANVI mapped clusters from the human synovium are highlighted. (**e**) Proportion of subclusters of myeloid cells along in healthy compared to PsA and psoriasis in CITEseq dataset of human PBMC (GSE194315). scANVI identified CD2^+^ MHC-II^+^ CCR2^+^ myeloid precursors are shown in Figure 3i. Graphs show median, quartiles and min–max; healthy subjects *N* = 29, PsA *N* = 20, psoriasis *N* = 33; *P*-values were calculated by edgeR differential abundance test and corrected for multiple testing using the BH-method (orange); statistically credible (cred.) and non-credible (n. cred.) changes in abundance were identified using scCODA (black). (**f**) Differential abundance of Milo neighbourhoods in PBMCs from healthy donors or patients with PsA (left) or psoriasis (right) shown as a bee swarm plot. Significantly enriched (FDR < 0.05, positive FCs, orange / blue) or depleted (FDR < 0.05, negative FCs, green) neighbourhoods among the original annotation as well as the scANVI identified CD2^+^ MHC-II^+^ CCR2^+^ myeloid precursors are highlighted.

**Supplementary Figure 3:**
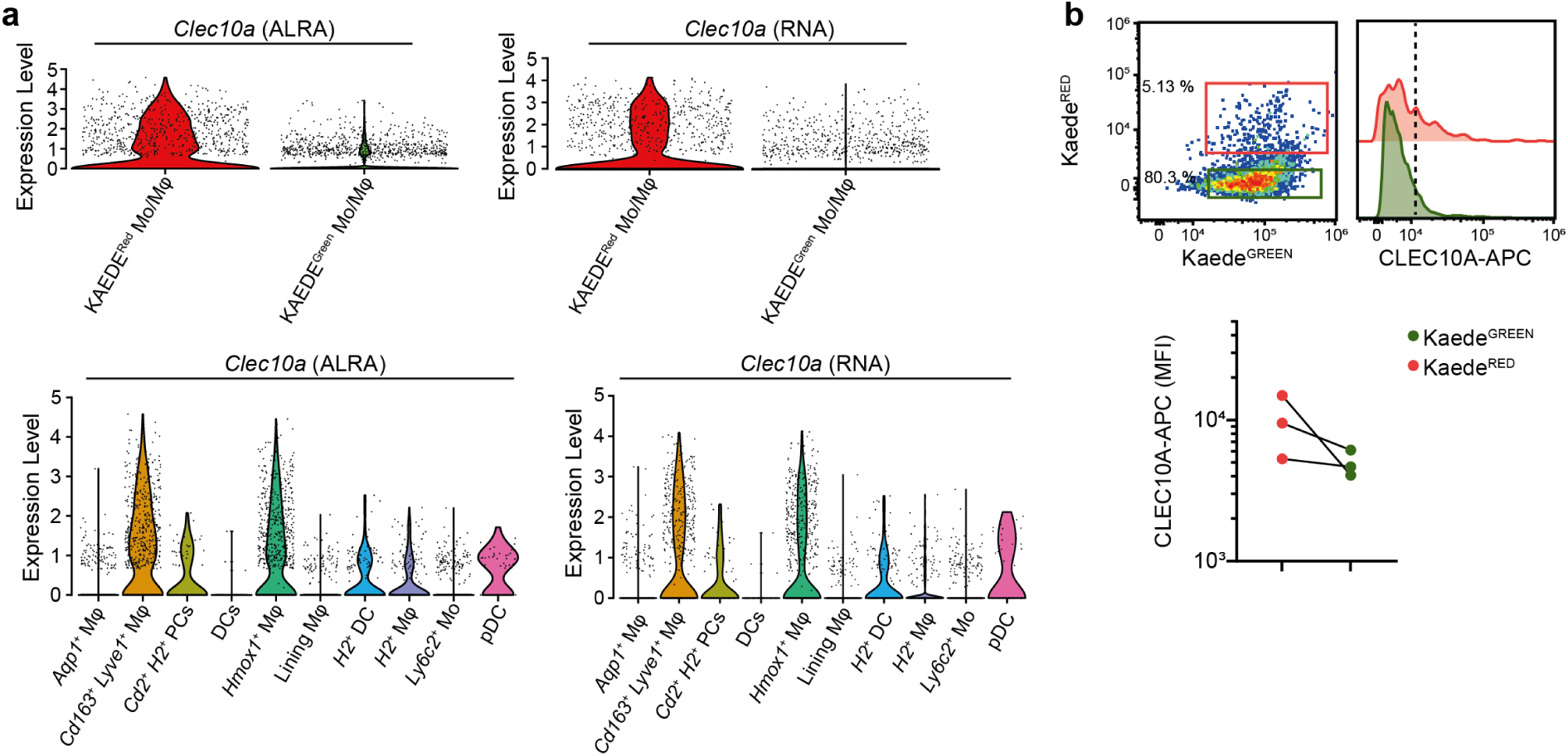
(**a**) ALRA-imputed (left) and RNA (right) expression of *Clec10a* in Kaede^RED^ and Kaede^GREEN^ myeloid cells (top) and among the subclusters of myeloid cells (bottom) the joint of the scRNAseq dataset from Kaede^tg^ mice on BALB/c and C57BL/6 background on day 7 of the IL-23OE. (**b**) Representative flow cytometry plots of CLEC10A expression on Kaede^RED^ versus Kaede^GREEN^ viable CD45^+^ CD11b^+^ Ly6G^-^ CD11c^-^ CCR7^-^ CD3^-^ B220^-^ cells in the joint on day 7 of the IL-23OE in Kaede^tg^ animals. Quantification of geometric median fluorescence intensity in Kaede^RED^ versus Kaede^GREEN^.

**Supplementary Figure 4:**
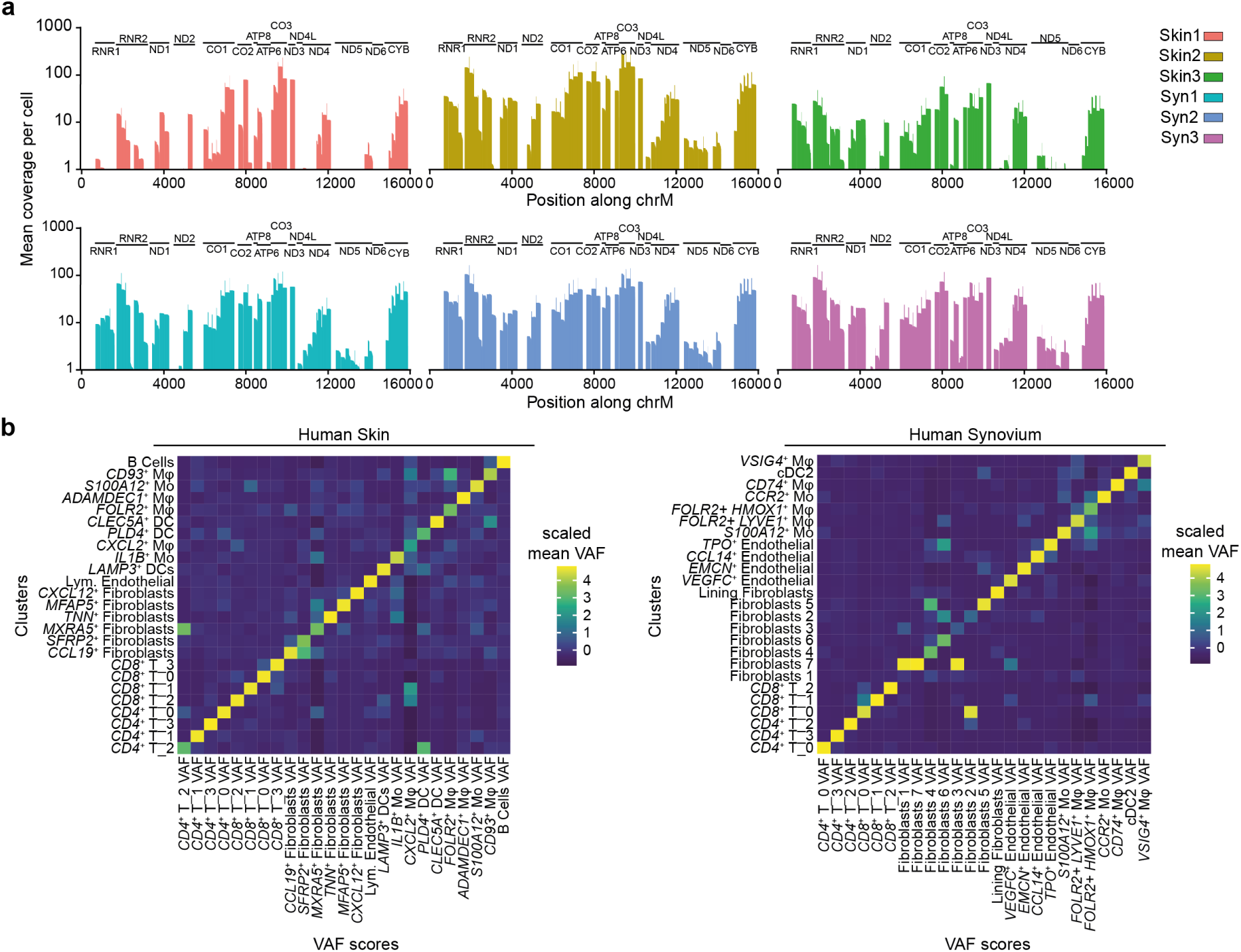
(**a**) Mean coverage over the mitochondrial genome in the MAESTER mitochondrial sequencing libraries for each sample. (**b**) Heatmap of mean variant allele frequencies (VAF) of the informative mitochondrial variants across subclusters for the skin (left) and the synovia (right) dataset.

## Supplementary Tables

**Supplementary Table 1.** Synovial tissue patient characteristics.

|  | **Psoriasis (*N* = 8)** | **PsA (*N* = 6)** | ***P*-value** |
| --- | --- | --- | --- |
| **Female, n (%)** | 4 (50) | 2 (33) | 0.6270 |
| **Age, mean (SD)** | 45 (12) | 59.7 (8) | 0.0216 |
| **Psoriasis disease duration years, mean (SD)** | 18.1 (13.7) | 11 (9.6) | 0.4908 |
| **PsA disease duration years, mean (SD)** | N/A | 1.2 (0.5) | N/A |
| **DAPSA, mean (SD)** | N/A | 22.7 (6.3) | N/A |
| **Krenn Score, mean (SD)** | 1.3 (0.7) | 4.2 (1.6) | 0.0087 |

N/A = not applicable; PsA = Psoriatic arthritis; SD = standard deviation. *P*-values were calculated by non-parametric t-test.

**Supplementary Table S2.** PBMCs patient characteristics.

|  |  |  |
| --- | --- | --- |
|  | **PsA (N = 29)** | 936 |
| **Female, n (%)** | 15 (51.7) |  |
| **Age, mean (SD)** | 53.2 (12) | 937 |
| **Disease duration years, mean (SD)** | 6.4 (4.9) |  |

PBMCs = peripheral blood mononuclear cells; PsA = Psoriatic arthritis; SD = standard deviation.

**Supplementary Table S3:**
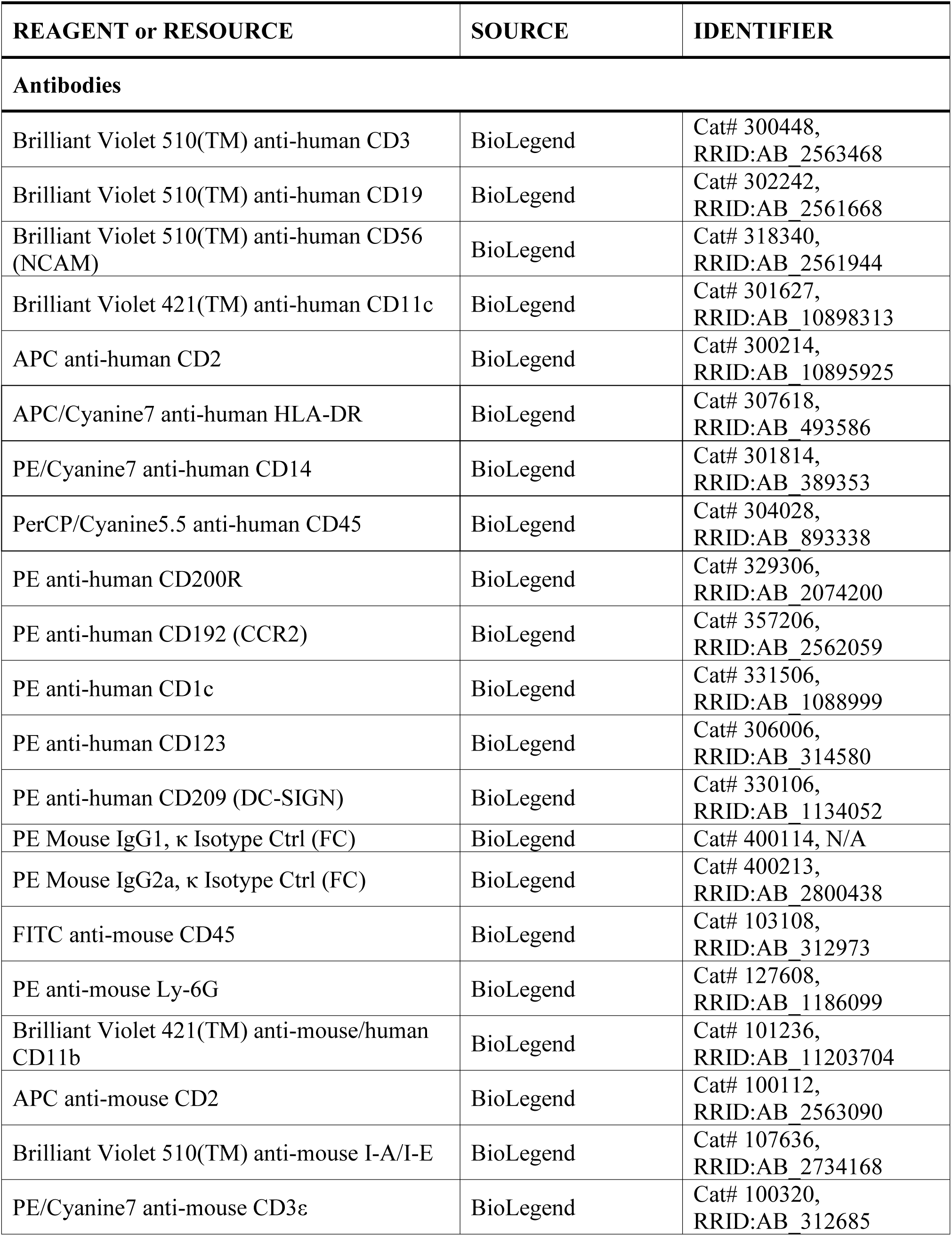
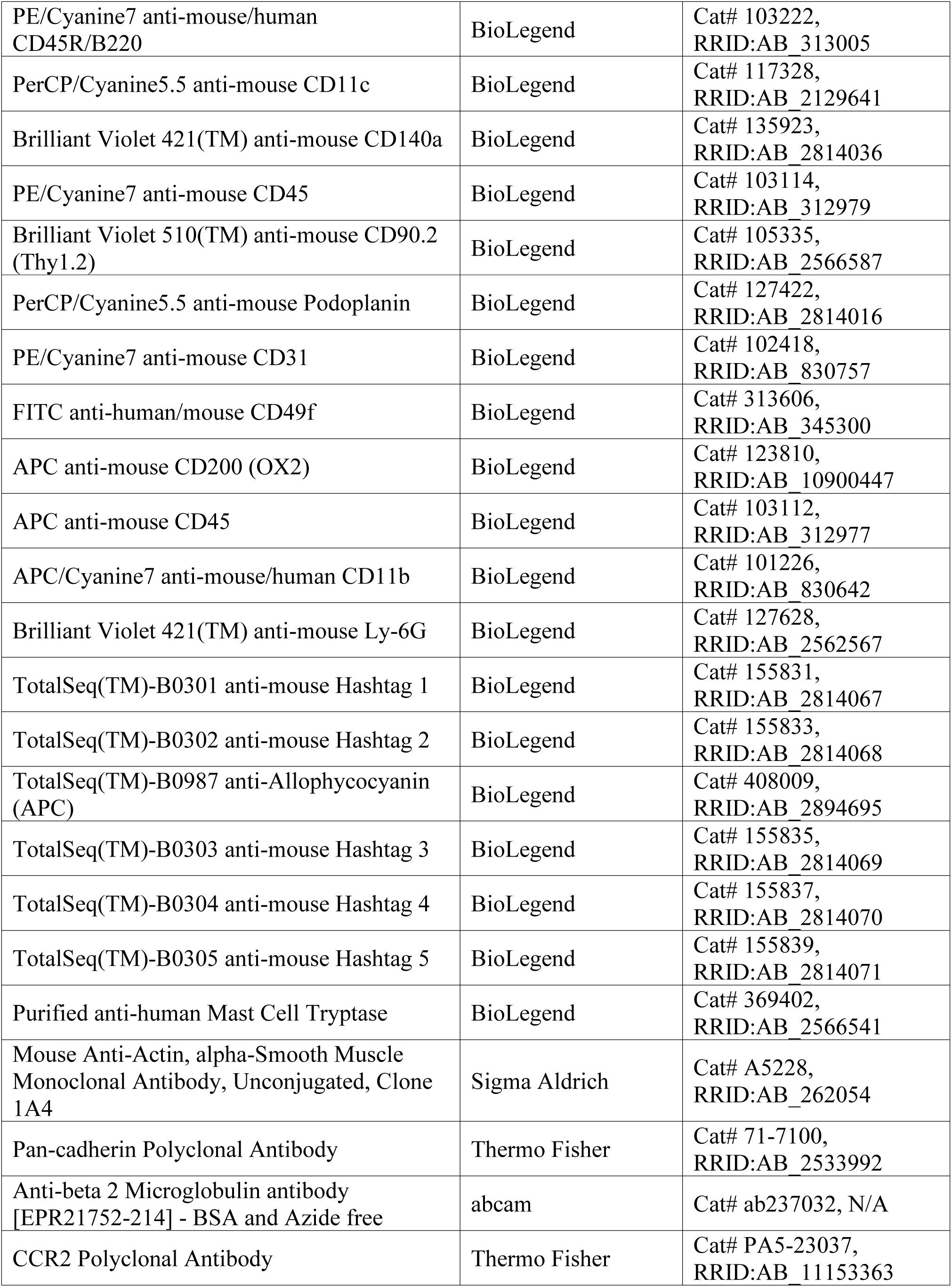
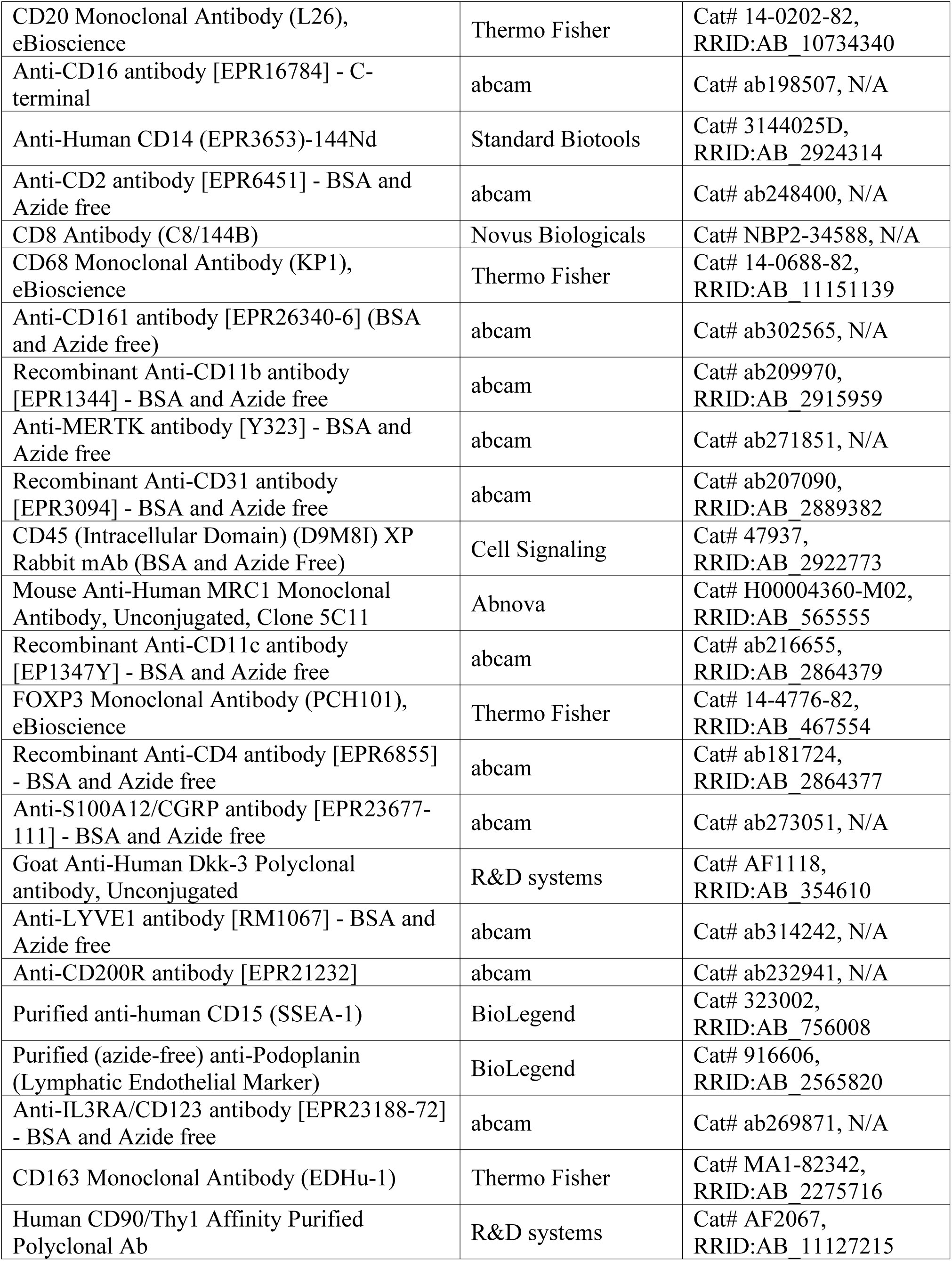
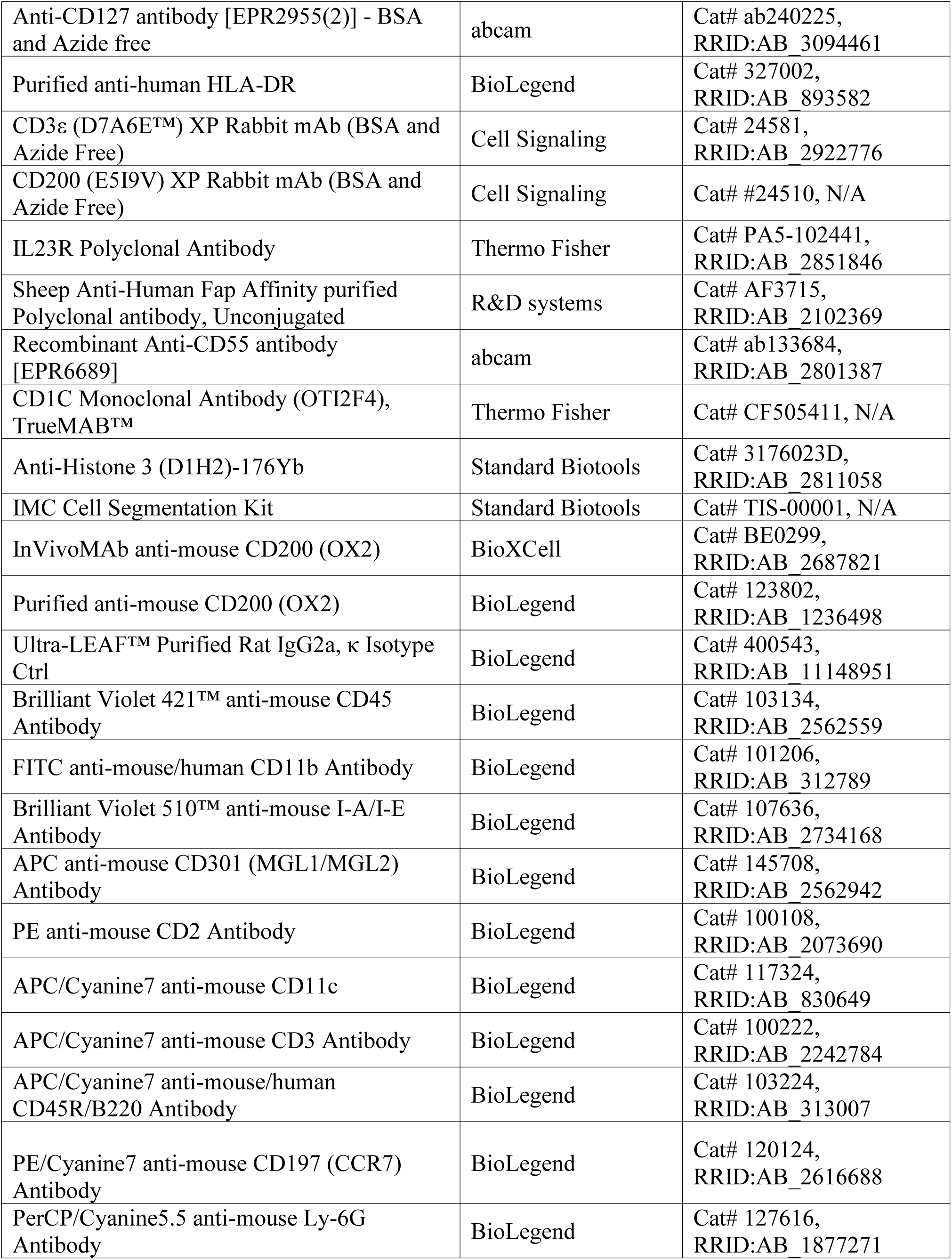
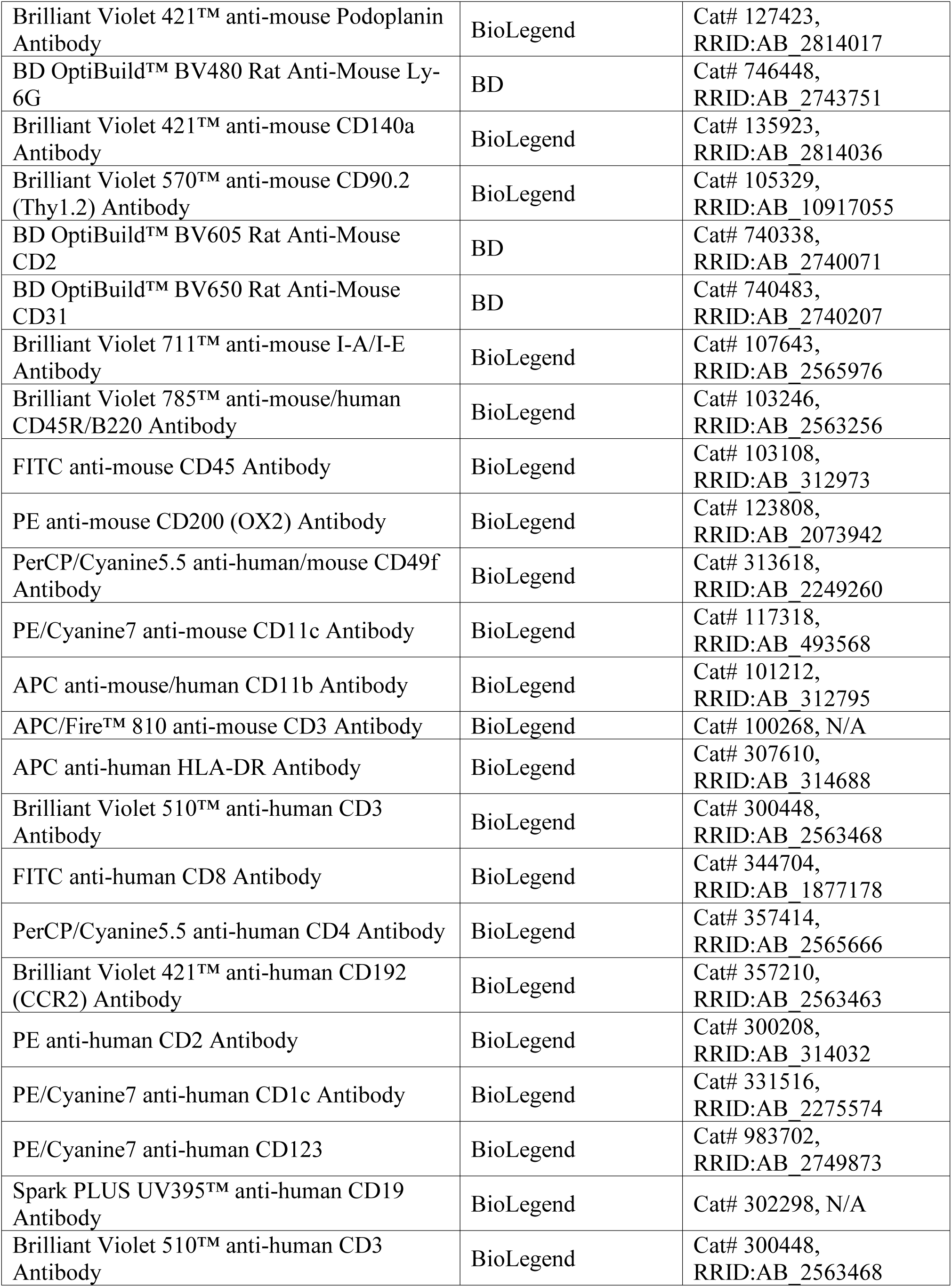
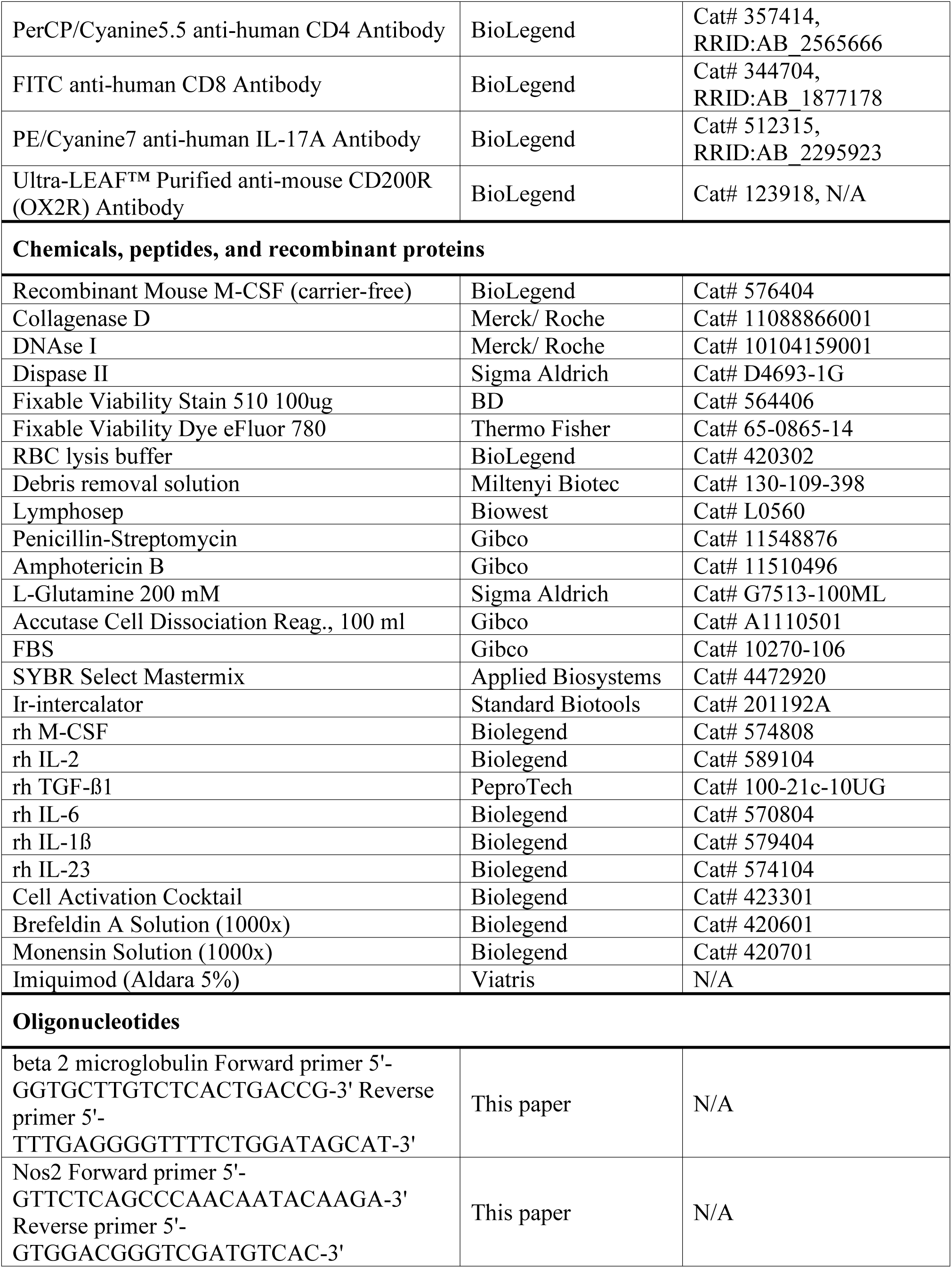
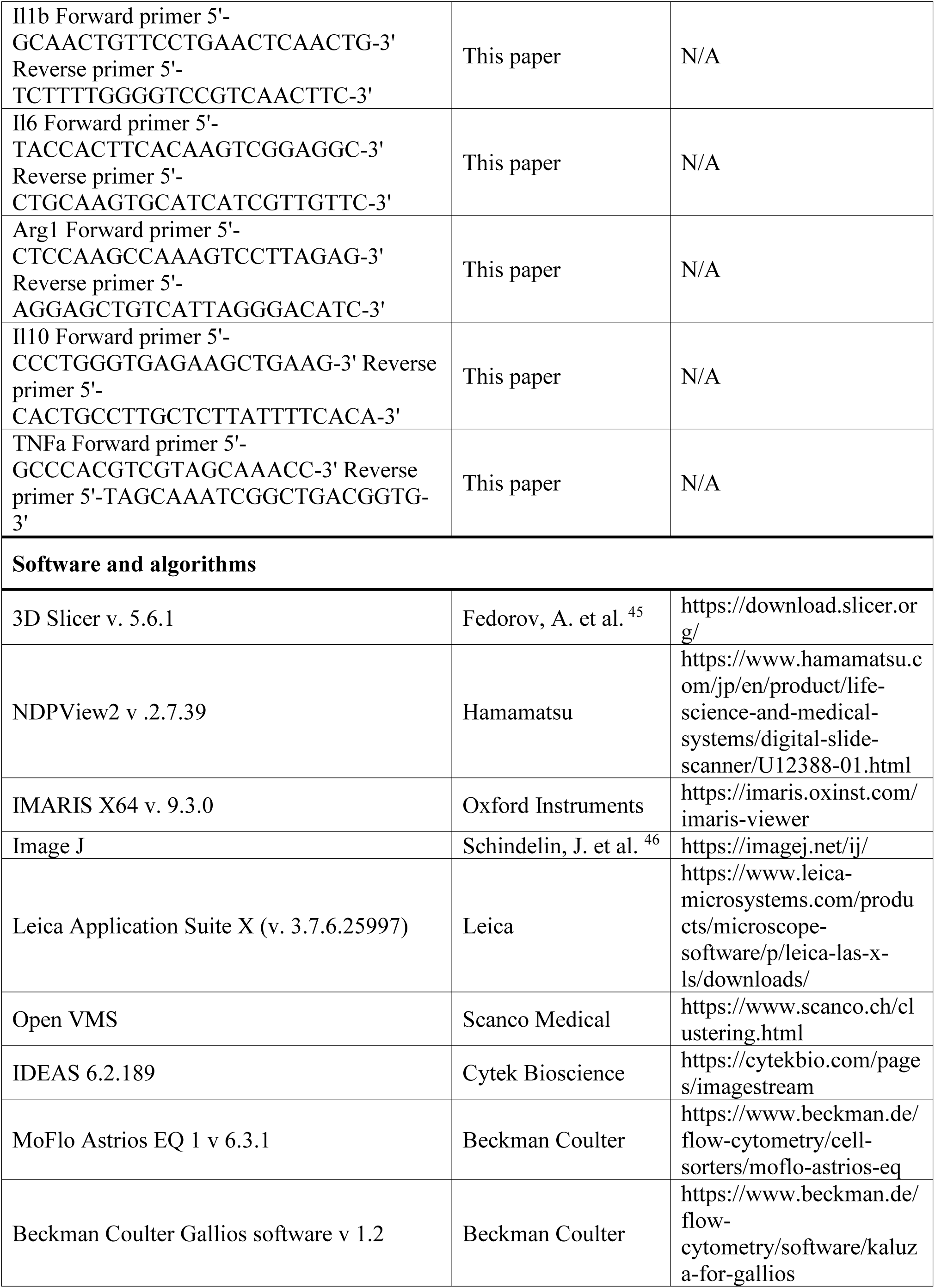
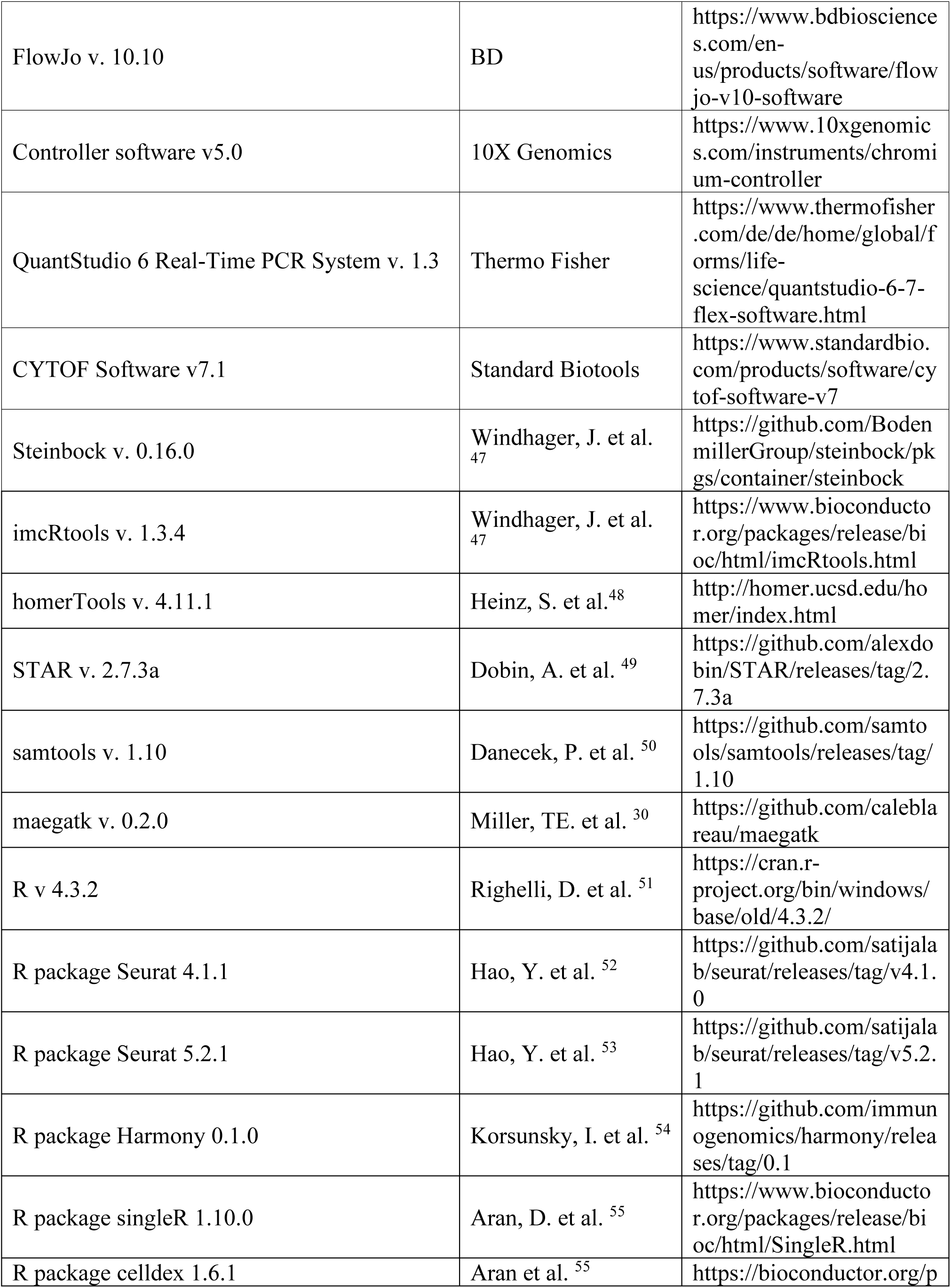
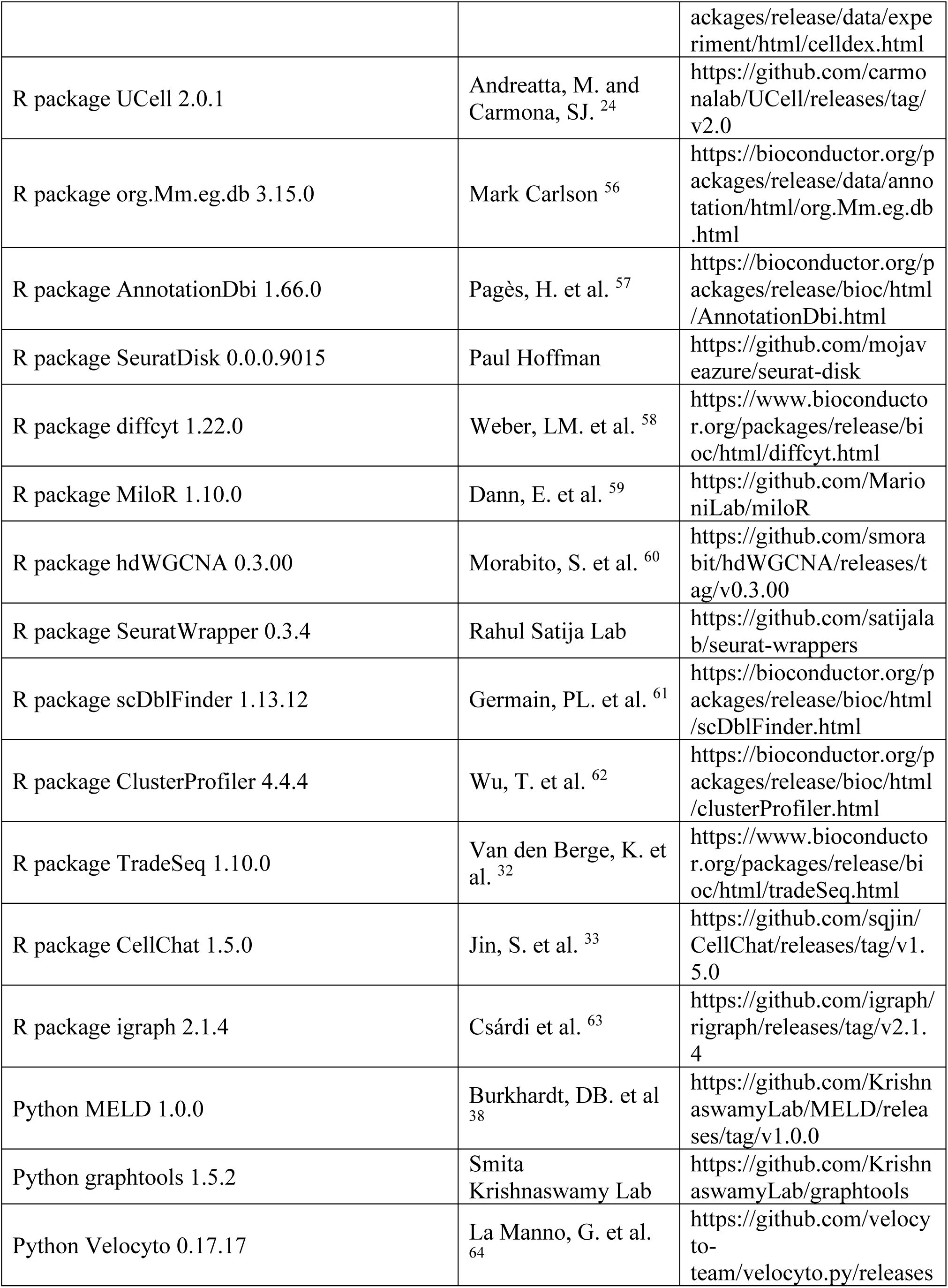
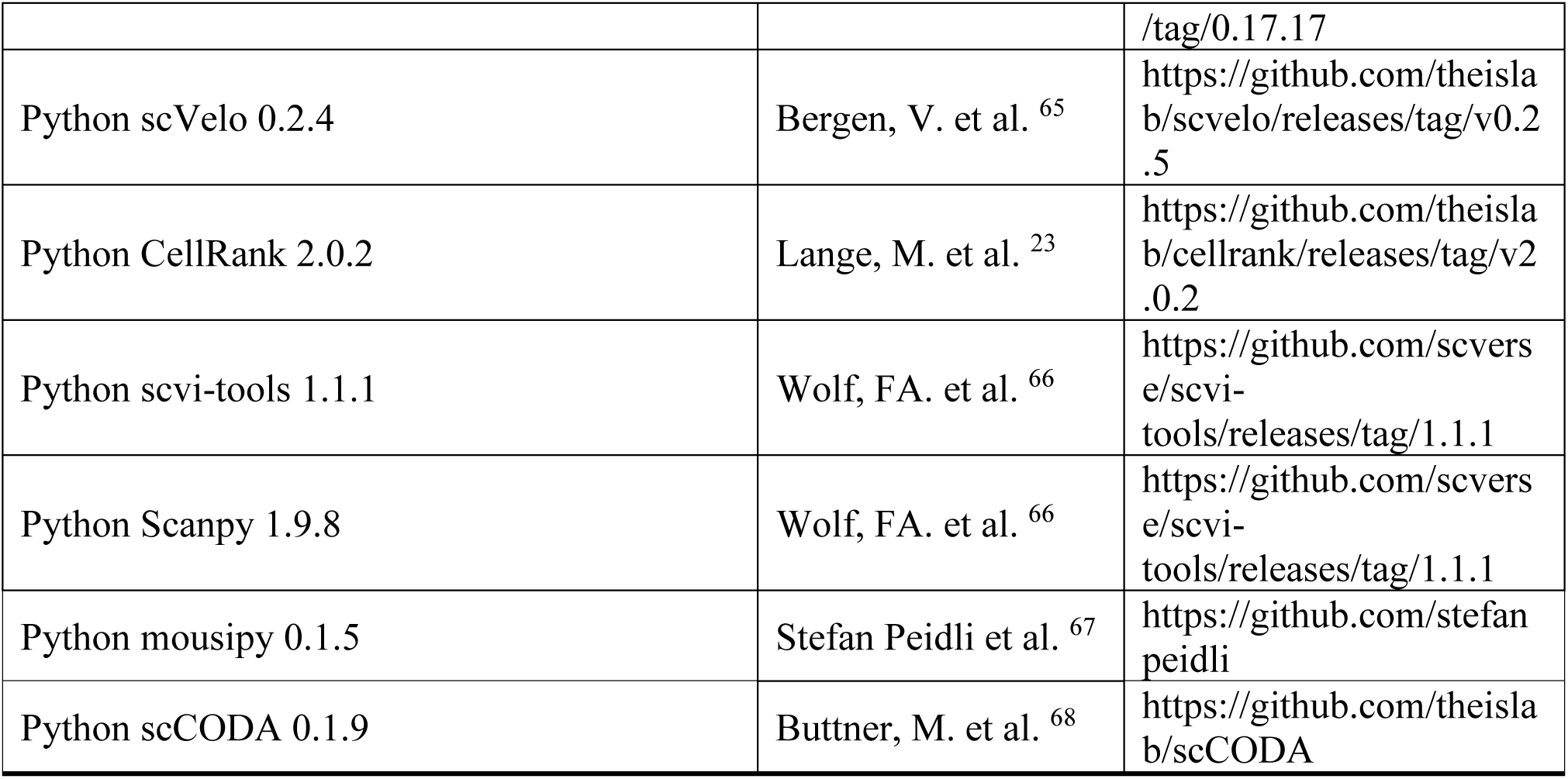
Resource table.

## Material and Methods

### Experimental approaches

Experiments were not performed in a blinded fashion, except where specifically stated. There were no exclusion criteria for human or animal experimentation. Mice were stratified by sex and then randomised to the different groups. Cells from human donors were also randomised.

### Human studies and patient characteristics

Human research was conducted in accordance with approved protocols by the institutional review boards of the Friedrich-Alexander-University (FAU) of Erlangen-Nürnberg, the Fondazione Policlinico Gemelli IRCCS and the Hospital Clinic de Barcelona. Ultrasound guided-minimally invasive synovial tissue biopsies were collected from early PsA (*N* = 6) and patients with psoriasis (*N* = 8) at the SYNGem Biopsy Unit of the Fondazione Policlinico Universitario A. Gemelli RCCS (ID: 4951-2022) and at the Uniklinikum Erlangen (ID: 18-334_4-Bio). Additional synovial tissue from healthy donors (HC; *N* = 3) was obtained at the Hospital Clinic de Barcelona (ID: HCB/2 020/0100). Punch biopsies (3 mm) from lesional skin were collected from psoriasis patients (*N* = 3) in Erlangen. Blood samples were collected from individuals with PsA (*N* = 29) in Erlangen. Patient information is provided in **Table S1** and **S2**. All patients included fulfilled the 2006 CASPAR classification criteria for PsA ^69^. Additional human datasets were obtained from online databases listed in the data availability section and referenced in the text and figure legends.

### Animals

BALB/c JRj and C57BL/6 NRj mice were obtained from the Janvier Laboratories. C-Kaede^tg^ (C.Cg-Tg(CAG-tdKaede)15Utr) and B6-Kaede^tg^ (B6.Cg-Tg(CAG-tdKaede)15Utr) were described in ^16^ and were kindly provided by Yoshihiro Miwa and Michio Tomura from the RIKEN BRC through the National BioResource Project of the MEXT/AMED, Japan. C-Kaede^tg^ and B6-Kaede^tg^ were backcrossed to BALB/c JRj and C57BL/6 NRj, respectively, and only heterozygous Kaede^tg^ mice were used in the experiments. All animals were maintained under specific-pathogen-free (SPF)-conditions with a 12-h day/night cycle, fed chow (sniff Spezialdiäten, V1534-000) and given water *ad libitum*. Room air temperature and cage climate were standardised at 20 - 24 °C, 45 - 65 % relative humidity, and 15 - 20 air changes per hour. Mice were housed in groups of three to five animals per cage. A period of one week was observed between delivery and the start of the study for test animals that did not come from our own breeding programme. All animals were 8-10 weeks old at the start of the study. The animal experiments were conducted in accordance with local regulations and approved by the government of Lower Franconia (protocol 55.2-2532-2-1061, 55.2-2532-2-1886 and 55.2-2532-2-2157).

### Il-23-induced animal model of psoriatic arthritis

*Il23* treatment (3 µg / mouse) was administered on the first day of scoring. Scoring was continued until day 21, the end of the model. The development of skin symptoms (scaling) was monitored and reported as follows: grade 0 (no scaling), 1 (mild scaling), 2 (moderate scaling) and 3 (severe scaling). C-Kaede^tg^ and B6-Kaede^tg^ heterozygous mice were treated with 6 µg / mouse. The *Il23* vector, which encodes the two subunits IL-12b (IL-12p40) and IL-23a (IL-23p19) linked by a flexible region under the control of the albumin promoter for efficient expression in hepatocytes, followed by a secretion sequence for efficient release into the circulation, was kindly provided by Stefan Wirtz (Department of Medicine 1, Friedrich-Alexander-Universität (FAU) Erlangen-Nürnberg). The vector was produced in *Escherichia coli* DH5α grown in terrific broth medium without removal of vector backbone. Plasmids were purified using the PureLink HiPure Plasmid Maxiprep Kit (Thermo Fisher) followed by the MiraCLEAN Endotoxin Removal Kit (Mirus) to ensure efficient removal of endotoxin. Next, 3 µg of naked plasmid DNA in Ringer’s solution was administered into the lateral tail vein by hydrodynamic gene transfer at a volume equivalent to 10 % of body weight ^70, 71^. Treatment with an antibody (clone OX-90, BioXCell) blocking CD200 (OX-2) was administered intraperitoneally every other day (100 µg/dose) in C57BL/6 animals and started on the same day as *Il23* treatment. Treatment with the agonistic anti-CD200R1 (OX-2R) antibody (clone OX-110, BioLegend) was administrated subcutaneously every 5 days in BALB/c animals and started 7 days after *Il23* treatment.

### Animal model imiquimod induced psoriasis

A 2 × 3 cm area on the dorsal skin of female BALB/c mice was shaved using an electric shaver, and any remaining hair was removed using a depilatory cream (Veet). A total of 100 mg of Aldara cream (Viatris), containing 5% IMQ, was applied daily for either 7 or 21 consecutive days. Mice treated for 21 days concomitantly received intraperitoneal injections of a CD200 (OX-2)-blocking antibody (clone OX-90; Bio X Cell) or a rat IgG2a isotype control (clone RTK2758; BioLegend) every other day (100 µg/dose).

### UV light exposure

Mice were anesthetized with O2/isoflurane (3 % (v/v) isoflurane, 1.2 l · min^-1^ O2) and placed on a heating pad (37 °C). The psoriatic skin of the lower back and of the proximal parts of the legs was shaved, while the rest of the body, including the distal parts of the legs and hind paws, was shielded with aluminium foil. The skin was exposed to a UV light source for 10 min according to a previous protocol ^16^. Briefly, a BlueWave LED PrimeCure UVA QX4 (Dymax) with a 3 mm diameter lens was used at 51 % power and a distance of 16 cm was maintained between the skin and the lenses. The exposure was repeated for two consecutive days (-48 h and -24 h) before sacrifice.

### MRI imaging

*In vivo* MRI of the hind paws was performed with a preclinical 7T MRI (BioSpec, Bruker BioSpin) using a volume resonator: RF RES 300 1H 075/040 QSN TR. The imaging protocol included pre- and post-contrast T1-weighted spin echo sequences and a T2 TurboRARE short tau inversion recovery (STIR) sequence. During the imaging, the mice received an intravenous bolus injection of a low molecular weight gadolinium chelate agent (0.15 mmol · kg^-1^ Gadobutrol, Bayer Vital) over a time period of 10 s via a tail vein catheter. Mice were laid on a heating pad (37 °C), were anesthetised with O2/isoflurane (3 % (v/v) isoflurane, 0.5 l · min^-1^ O2) and eye ointment was applied. The volume of joint inflammation was evaluated on either STIR or T1-weighted images after subtraction of pre- from post-gadolinium was performed, and the volume of contrast-enhancing areas of the ankle joint was quantified by manual segmentation (3D Slicer ^45^, version 5.6.1).

### Histological processing of mouse samples

Hind paw joints were fixed in PBS containing 4 % (w/v) formaldehyde for 16 - 24 h. After removal of the skin, the bone tissue was decalcified in 0.5 M EDTA pH 7.4 before embedding and sectioning. Tissues were dehydrated, infiltrated and embedded in paraffin, cut into 1 µm thin sections and mounted on standard histological slides. Thin sections were deparaffinized by heating the slides at 65 °C for 30 min, washed in Histo Clear (National Diagnostics) and rehydrated through a series of 100 % (v/v) ethanol, 95 % (v/v) ethanol, 80 % (v/v) ethanol, 60 % (v/v) ethanol and water. Mayer’s haematoxylin solution (Merck) was applied for 10 min. Excess haematoxylin was washed off and the stain was blued by rinsing the slides for 10 min in tap water. Eosin counterstain (0.3 % (wt/v) Eosin Y (Sigma), 0.0 1 % (v/v) acetic acid) was applied for 3 min and excess stain was washed off with deionized water. The sections were again dehydrated in isopropanol, mounted with Roti Mount mounting medium (Carl Roth). Slides were digitalized with a NanoZoomer S60 (Hamamatsu) and regions of interest were exported as *.TIFF files. Epidermal thickness was measured using the NDPView2 (2.7.39) program (Hamamatsu). A periarthritis score was established to assess the inflammatory infiltrate in the joints: grade 0 (no infiltrate), 1 (mild infiltration), 2 (moderate infiltration) and 3 (severe infiltration).

### Microscopy image analysis

Immediately after sacrifice, the hind paws were dissected and the skin was removed. Samples were immersed in -20 °C cooled acetone and stored at -20 °C overnight for fixation. For light sheet microscopy ethyl cinnamate was used to clear the tissue and imaging was performed using UltraMicroscope II (LaVision). Kaede^GREEN^ and Kaede^RED^ were detected in the 488 and 594 channel. For fluorescence microscopy, the hind paws were decalcified in decalcification buffer for 2 days after fixation. Tissues were cryoprotected in 30% sucrose and 10% PVP in PBS, embedded in 30% gelatine, and sectioned at 40 µm. Imaging was conducted using a Leica Thunder Imager 3D Assay with a 63x oil immersion objective (NA 1.4-0.6) with subsequent large volume computational clearing and maximum intensity projection. The imaging analysis was performed with IMARIS X64 (Oxford Instruments) software (v. 9.3.0) and Fiji (v. 1.52)^46^.

### Microcomputed tomography imaging and analysis

Structures of tibial bones and paws were measured with a SCANCO Medical μCT 40 or µCT45 scanner and analysed with SCANCO Application Center evaluation software for segmentation, 3D morphometric analysis, density and distance parameters (SCANCO Medical). The three-dimensional modelling of the bone was performed with optimised greyscale thresholds of the operating system Open VMS (SCANCO Medical) and with XamFlow-Workflow ^72^.

### Preparation of single cell suspensions from mouse tissues

After sacrifice, the skin covering both legs was shaved and isolated, the subcutaneous fat was removed and the tissue was minced with scalpels. The legs with intact ankles were dissected and the nails were cut-off. The tibia was removed by dislocation and the tendons and joint capsules were cut to facilitate enzymatic dissociation. Skin and joint tissues were digested in RPMI 1640 containing 1.2 mg · ml^−1^ collagenase D (Roche), 0.6 (skin) and 0.2 (joint) mg · ml^−1^ Dispase II (Sigma-Aldrich) and 0.2 mg · ml^−1^ DNase I (Roche). Samples were incubated three times at 37 °C for 20 min with constant shaking (2,000 rpm) on a thermal shaker (Eppendorf). After 20 min, the supernatant was collected, filtered (70 µm) and fresh digestion medium was added. The digestion was repeated for a total of 3 times. Red blood cell (RBC) lysis was performed using RBC lysis buffer (BioLegend) according to the manufacturer’s recommendations. Debris removal solution (Miltenyi Biotec) was used to remove impurities according to the manufacturer’s protocol. Small intestine samples were collected and, after removal of the mesentery, pre-soaked and rinsed in cold DMEM (Gibco) containing 20 % (v/v) FBS (Gibco). The tissue was then cut longitudinally and incubated with agitation in HBSS (without Ca^2+^) (Gibco) containing 2 mM EDTA. The mucosal epithelium was isolated and the subepithelial tissue was further washed with HBSS (without Ca^2+^) (Gibco) and cut into 5 mm^2^ pieces for enzymatic digestion with HBSS (Gibco) (with Ca^2+^ and 2 % (v/v) FBS) containing 3.0 mg · ml^−1^ collagenase D (Roche) and 0.2 mg · ml^−1^ DNase I (Roche). Samples were incubated three times for 10 minutes at 37 °C with constant shaking (800 rpm) on a thermal shaker (Eppendorf). The resulting cell suspension was filtered (100 µm), washed with DMEM (Gibco) containing 10% (v/v) FBS (Gibco) and pooled with the mucosal isolated cells. Cell counts were determined by trypan blue staining using an automated cell counter (Bio-Rad). For the isolation of bone marrow cells, the femur and tibia were collected, cleaned of muscle and tendons and cut at both ends. After centrifugation at 10,000 g for 15 sec, the resulting pellet was treated with RBC lysis buffer (BioLegend) and washed. The resulting cell suspensions from all tissues were filtered (40 µm) and cell counts were determined by trypan blue staining using an automated cell counter (Bio-Rad).

### Preparation of single cell suspensions from human tissue

Fresh human skin and synovial tissue or fresh frozen human synovial tissues were thawed, thoroughly minced and digested in RPMI 1640 containing 2 mg · ml^−1^ collagenase D (Roche), 0.2 mg · ml^−1^ Dispase II (Sigma-Aldrich) and 0.2 mg · ml^−1^ DNase I (Roche) for synovial tissue or 7.5 mg · ml^−1^ collagenase D (Roche), 2.5 mg · ml^−1^ Dispase II (Sigma-Aldrich) and 0.5 mg · ml^−1^ DNase I (Roche) for skin. Samples were incubated three times at 37 °C for 20 min with constant shaking (2,000 rpm) on a thermal shaker (Eppendorf). After 20 min, the supernatant was collected, filtered (70 µm) and fresh digestion medium was added. The digestion was repeated for a total of 3 times. RBC lysis was performed using RBC lysis buffer (BioLegend) according to the manufacturer’s recommendations. The resulting cell suspensions were filtered (30 µm) and counted in trypan blue using a Neubauer counting chamber (Brand).

### PBMC isolation

Human EDTA blood samples were collected from patients with PsA at the outpatient clinic of the University Hospital Erlangen, as part of routine diagnostics. Written informed consent was obtained from all patients. Peripheral blood mononuclear cells (PBMCs) were purified using Lymphosep (Biowest). Briefly, 6 ml of EDTA blood was mixed with 6 ml of DPBS (Gibco), underlaid with 4 ml of Lymphosep and centrifuged at 1,000 g for 30 min without break. The plasma component was discarded and the PBMC ring was collected, washed in 50 ml PBS by centrifugation at 200 g for 30 min. Erylysis (RBC Lysis buffer, BioLegend) and a second centrifugation at 100 g for 30 min was performed to remove RBCs and thrombocytes, respectively.

### Flow cytometry and cell sorting

Cells were stained for 30 min on ice in PBS containing 2 % (v/v) FBS and 5 mM EDTA and then fixed with 1 % formaldehyde for 20 min. Single dye staining and fluorescence minus one (FMO) controls were used to compensate and set up the gating, respectively. Dead cells were excluded using FVS510 (1:200, BD) or eFluor 780 Fixable Viability Dye (1:4000, eBioscience). Detailed information about all antibodies is listed in the **Table S3**. Anti-CD45 (30-F11; 1:1000 dilution), anti-CD3ε (145-2C11, 1:100 dilution), anti-CD45R/B220 (RA3-6B2, 1:200 dilution), anti-Ly-6G (1A8, 1:100 dilution) and anti-CD11b (M1/70, 1:1000 dilution) were used to identify/sort Kaede^RED^ skin-derived migrating cells in the hind paws. Antibodies used for identification of migrating myeloid precursor in the skin and in the hind paws included anti-CD45 (30-F11; 1:1000 dilution), anti-Ly6G (1A8; 1:1000 dilution), anti-CD11b (M1/70, 1:1000 dilution), anti-CD2 (RM2-5; 1:1000 dilution), anti-I-A/I-E (M5/114.15.2; 1:500 dilution), anti-CD3ε (145-2C11, 1:100 dilution), anti-CD45R/B220 (RA3-6B2, 1:200), anti-CD11c (N418; 1:500 dilution). Antibodies used to analyse the CD200 expression in fibroblast compartment were anti-CD140a (APA5; 1:500 dilution), anti-CD45 (30-F11; 1:1000 dilution), anti-CD90.2 (30-H12; 1:500 dilution), anti-Podoplanin (8.1.1; 1:200 dilution), anti-CD31 (390; 1:500 dilution), anti-CD49f (GoH3; 1:200 dilution), anti-CD200 (OX-90; 1:100 dilution). Antibodies used for adoptive cell transfer experiment included anti-CD45 (30-F11; 1:500 dilution), anti-Ly6G (1A8; 1:500 dilution), anti-CD11b (M1/70, 1:200 dilution), anti-CD2 (RM2-5; 1:500 dilution), anti-I-A/I-E (M5/114.15.2; 1:100 dilution), anti-CD3ε (145-2C11, 1:100 dilution), anti-CD45R/B220 (RA3-6B2, 1:200 dilution), anti-CD11c (N418; 1:500 dilution), anti-CCR7 (4B12, 1:100 dilution) and anti-CLEC10A (LOM-14, 1:100 dilution).

PBMCs were freshly analysed by flow cytometry for anti-CD2 (RPA-2.10; 1:1000 dilution), anti-CCR2/CD192 (K036C2; 1:500 dilution), anti-CD200R1 (OX-108; 1:100 dilution), anti-HLA-DR (L243; 1:1000 dilution), anti-CD14 (M5E2; 1:100 dilution), anti-CD16 (3G8; 1:100 dilution), anti-CD45 (HI30; 1:100 dilution), anti-CD3 (UCHT1; 1:100 dilution), anti-CD19 (HIB19; 1:200 dilution), anti-CD56 (HCD56; 1:200 dilution), anti-CD11c (3,9; 1:100 dilution), anti-CD1C (L161; 1:1000 dilution), anti-CD123 (6H6; 1:1000 dilution), anti-CD209 (9E9A8; 1:200 dilution) expression in the myeloid fraction in human together with isotype IgG1, κ (MOPC-21; 1:100 dilution) and IgG2a, κ (MOPC-173; 1:200 dilution). PBMCs used for sorting and analysis of co-culture experiments were stained for anti-CD3 (UCHT1; 1:200 dilution), anti-CD19 (HIB19; 1:200 dilution), anti-CD8a (SK1, 1:1000 dilution), anti-CD4 (A161A1, 1:200 dilution), anti-CCR2 (K036C2, 1:500 dilution), anti-HLA-DR (L243, 1:200 dilution), anti-CD2 (RPA-2.10; 1:200 dilution), anti-CD1C (L161; 1:500 dilution), anti-CD123 (6H6; 1:500 dilution), anti-IL17A (BL168, 1:500 dilution). IL-17A was stained intracellularly using the IC fixation kit (Thermo Fisher) according to the manufacturer’s recommendations. All antibodies were purchased from BioLegend unless stated otherwise. The acquisition of cytometry data was performed with Beckman Coulter Gallios software v 1.2, whilst Beckman Coulter MoFlo Astrios EQ (running with BD Astrios Summit v6.3.1) was used for cell sorting. Sorted populations were reanalysed to determine target cell purity after sorting (>98% purity). Flow cytometry data were analysed with FlowJo (v. 10.10, BD Biosciences).

### Adoptive cell transfer approach

Donor BALB/c mice were sacrificed 7 days after *Il23* treatment. Hind paws were isolated, digested and resulting cell suspension was stained with a cocktails of flow cytometry antibodies as described above. Skin-derived myeloid cells were sorted from hind paws of donors and adoptively transfer in hind paws of recipient BALB/c mice via intra-articular injection of the tarsal joints (50.000 cells/20 µl /paw) 7 days after *Il23* treatment.

### Imaging cytometry

All samples were acquired using a 12-channel Amnis ImageStreamX Mark II (Cytek Biosciences) imaging flow cytometer equipped with three excitation lasers: 405nm (120 mW), 488nm (200 mW), and 642 nm (150 mW), and a MultiMag with three objectives lenses (20 x, 40 x, and 60 x magnification). Samples were acquired at 60 x magnification, and the excitation lasers (405 nm, 488 nm and 642 nm) were used at 80 mW, 5 mW and 150 mW, respectively.

Image analysis was performed using image-based algorithms in the ImageStream Data Exploration and Analysis Software (IDEAS 6.2.189, Cytek). Typical files contained imagery for 50,000 to 100,000 cells. Analysis was restricted to single cells in best focus. Single cells were identified by their intermediate size (area) and high aspect ratio (minor axis divided by the major axis) in comparison to debris (small area and a range of aspect ratios depending on the shape of the debris) and doublets (large area and small aspect ratio). Out-of-focus events were excluded by using the feature bright-field gradient RMS, a measurement of image contrast.

### Cell Culture

#### Mouse fibroblast – macrophage co-culture

Single cell suspensions form digested hind paws of healthy C57BL/6 mice was seeded in complete DMEM/F12 containing 10 % FBS (Gibco) 100U · ml^-1^ Penicillin-Streptomycin (Gibco), 0.5ug · ml^-1^ Amphotericin B (Gibco) and 20 mM L-Glutamine (Sigma) for fibroblast expansion. The medium was changed on day 3 and the cells were passaged on day 6 using trypsin-EDTA (0.25%, Gibco). 1 x 10^5^ cells of the second passage were seeded per well in a 6-well plate. In parallel, 20 x 10^6^ bone marrow cells obtained from the same mice were seeded in IMDM (Gibco) containing 10% FBS (Gibco), 100U · ml^-1^ Penicillin-Streptomycin (Gibco), 0.5ug · ml^-1^ Amphotericin B (Gibco), 20 mM L-Glutamine (Sigma) and 20 ng · ml^-1^ rmM-CSF (BioLegend) in a 100 mm petri-dish. Medium was changed on day 4 and on day 6 macrophages were detached by accutase (Gibco). 5 x 10^5^ cells were seeded with the fibroblasts in complete IMDM medium enriched with 20 ng · ml^-1^ rmM-CSF. The following day CD200 (OX2) antagonist (clone OX-90, BioLegend) 10µg ml^-1^ or isotype (Clone RTK2758, BioLegend) were added to the medium. After 24 hours, cells were harvested and RNA isolation was performed as described below.

#### Human monocyte – T cell co-culture

Flow cytometry-based cell sorting was used to isolate CD2^+^ and CD2^−^ myeloid cells as well as autologous CD8^+^ or CD4^+^ T cells from PBMCs of PsA patients. Cells were cultured in CD3 coated (1 μg/mL anti-human CD3 (clone UCHT1) in DPBS (gibco), 4°C overnight) 96-F-well plates at a ratio of 1:5 (myeloid cells : T cells) in RPMI-1640 medium supplemented with 10% heat-inactivated FBS, 2 mM L-glutamine, 100 U/mL penicillin, 100 µg/mL streptomycin, 1× nonessential amino acid solution and 50 µM β-mercaptoethanol. Human M-CSF (50 ng/ml, BioLegend), rhIL-2 (10 ng/mL, BioLegend), rhTGF-β (5 ng/ml, BioLegend), rhIL-6 (25 ng/ml, BioLegend), rhIL-1β (10 ng/ml, BioLegend), rhIL-23 (25 ng/ml, BioLegend) were added to the culture. Cells were maintained at 37°C, 5% CO2 for 6 days, with medium and cytokine replenishment every 48 hours. On day 6, cells were restimulated with cell activation cocktail (PMA/Ionomycin) without Brefeldin A (1:1000 dilution, BioLegend) for 1 hour, followed by monensin and brefeldin A (1:1000 dilution, BioLegend) for 3 hours. Then cells were harvested for analysis by flow cytometry as described above.

### Gene expression analysis

RNA isolation was performed using the Nucleo Spin RNA isolation kit according to the manufacturers’ protocol (Macherey-Nagel). Immediately following extraction, the total RNA concentration and A260:A280 ratio of each sample was determined via NanoDrop 2000 (Thermo Scientific). 8.2 μL of RNA was used to transcribe mRNA to cDNA following the manufacturer’s protocol (Applied Biosystems). The cDNA synthesis reaction was performed in a total volume of 20 μL, containing 8.2 μL RNA, 2 μL 10X PCR buffer Ⅱ, 4.4 μL MgCl2 solution 25 mM, 4 μL DNTP Mix, 0.5 μL 10X RT Random primers, 0.4 μL RNase Inhibitor and 0.5 μL Multiscribe Reverse Transcriptase. The reaction mixture was incubated at 20°C for 10 min, followed by 48°C for 30 min and 95°C for 5 min to inactivate the enzyme. Real-time PCR was performed in triplicate using SYBR Green master mix (Applied Biosystems) and QuantStudio 6 Real-Time PCR System v. 1.3. Fold change (FC) expression of target genes was calculated by the ΔCt, ΔΔCt and 2^-ΔΔCt^ comparative method for relative quantification after normalization. The endogenous control was β2 microglobulin. A list of primers is provided in the **Table S3**.

### Generation and sequencing of droplet-based single cell RNA sequencing libraries

Skin and joint cells from six C-Kaede^tg^ and nine B6-Kaede^tg^ mice with equal sex distribution were pooled and sorted for Ly-6G^-^ viable Kaede^RED^ CD45^+^, Kaede^GREEN^ CD45^+^ and Kaede^GREEN^ CD45^-^ fractions on day 7 of IL-23OE. Purified cells were stained for 30 min on ice in PBS containing 1 % (w/v) BSA with hashtag antibodies (TotalSeq-B0301/B0302/B0303/B0304B0305 anti-mouse hashtag 1/2/3/4/5, respectively, and anti-APC-B0987 (customized), all from BioLegend) to distinguish all six fractions per inbred strain. Cells were washed, counted and concentrated to 1,000 cells / µl prior to pooling. 24,000 cells (hyper-loading) equally pooled from each strain were loaded into a single well of a Chromium chip G (10x Genomics). Human synovial cells were loaded immediately after tissue dissociation with up to 25,000 cells in a single well of a Chromium chip G (10x Genomics), as described above. 3’ gene expression libraries were generated using Chromium Next GEM Single Cell 3’ Kit 3.1 with 3’ Feature Barcode Kit and dual indexing (10x Genomics protocol CG000316 Rev C). Libraries were sequenced as paired end (PE) 150 bp by Illumina sequencing to 65-80% saturation. Reads were mapped to the mm10 mouse genome (GENCODE vM23/Ensembl 98) or the GRCh38 human genome (GENCODE v32/Ensembl98) using the 10x Genomics Cell Ranger pipeline (6.0.0 and 7.2.0 respectively) with default settings.

### Generation of MAESTER libraries

To trace the lineage of specific cell populations and investigate potential migration from skin to joint in psoriatic disease, we applied the Mitochondrial Alteration Enrichment and Single-cell Transcriptome for Enhanced Resolution (MAESTER) protocol ^30^. In brief, single-cell suspensions were generated from paired lesioned skin and synovial biopsies obtained from PsA or psoriasis at risk patients and loaded onto a 10x Genomics Chromium controller using Single Cell 3’ v3.1 chemistry. Following droplet encapsulation and barcoding, cDNA synthesis and amplification were performed according to the manufacturer’s protocol. 25% of cDNA was processed for standard scRNAseq as described above, while a separate part of the cDNA was processed using the MAESTER workflow. This included an initial amplification, followed by a targeted PCR to enrich mitochondrial DNA (mtDNA) amplicons, and a final PCR to append sequencing adapters and sample indices. Sequences of mtDNA enriching primers and sequencing adapters are listed in the original publication ^30^.

Libraries were sequenced on an Illumina NovaSeq X system using standard 10x Genomics settings for scRNAseq and a custom configuration for MAESTER libraries (28 bp Read 1, 8 bp i7, 8 bp i5, 256 bp Read 2). Sequencing data were demultiplexed and aligned to both nuclear and mitochondrial genomes to enable integrated transcriptomic and lineage analysis at single-cell resolution. Reads were mapped to the GRCh38 human genome (GENCODE v32/Ensembl98) using the 10x Genomics Cell Ranger pipeline (9.0.1) with default settings.

### Quality control and annotation of single cell RNA sequencing libraries (mouse)

The two scRNAseq libraries from Kaede^tg^ animals after IL-23OE were first subjected to hashtag antibody oligonucleotide (HTO) demultiplexing using Seurat (4.1.1) ^52^ in R (4.2.1). The “HTOdemux” function with 0.95 positive quantile resulted in the separation of six libraries encoding for skin and Kaede^RED^ CD45^+^, Kaede^GREEN^ CD45^+^ and Kaede^GREEN^ CD45^-^ fractions as well as the exclusion of doublets and negative single cells. Quality control was performed to exclude low quality events by individually examining the distribution of quality criteria including the number of unique molecular identifiers (UMIs) >= 1,000, number of genes detected >= 550, complexity (log10(genes/UMI) > 0.8 and the ratio of mitochondrial genes

(mitoRatio) < 0.09. Genes expressed in fewer than 10 cells were excluded. From this stage, skin and joint datasets were analysed separately. Unwanted variations after log-normalisation in Seurat (“NormalizeData”, “FindVariableFeatures”, “ScaleData”, “RunPCA”) were cycle phase scores and mitoRatio, which were unevenly distributed in the Principal Component (PC) Analysis (PCA) space and regressed out before further analysis. For normalization, “SCTransform” (v2) ^73^ was applied individually to the raw UMI counts of each library, selecting the top 3,000 variable genes from each dataset. Technical variations resulting from library processing and sequencing were eliminated using Harmony (0.1.0) ^54^ integration on the merged SCT-transformed counts using the top 50 PCs. The top 50 Harmony components were used for clustering and visualisation. Seurat’s default clustering (“FindNeighbors” and “FindClusters”) with resolution = 1.5 and 2 resulted in 29 clusters in the skin and 32 clusters in the joint dataset. A single cluster in the joint was identified as debris due to its high mitoRatio and high background gene expression and was therefore removed before repeating harmony integration. In both tissues singleR (1.10.0) ^55^ using the global Immgen dataset ^74^ from celldex (1.6.0) ^55^ with “tune.thresh” = 0.001 was used for reference based annotation. UMAP visualization was generated using “RunUMAP”. Characteristic gene signatures of each cluster were identified using “PrepSCTFindMarkers” on SCT assays followed by “FindAllMarkers” with default parameters. To label the clusters identified by Seurat, both reference-based annotation and manual annotation based on the identified markers ^17, 18, 19, 20^ were used.

### Post-processing of myeloid clusters (mouse)

Harmony integration was repeated for Kaede^RED^ myeloid cells as previously described. Using the Immgen mononuclear phagocytes dataset (GSE122108) ^21, 22^, a second reference-based annotation was performed. After generating the UMAP visualisation in Seurat, “FindSubCluster” was used to identify subsets of DCs, monocytes and macrophages among the *H2*^+^ myeloid cells in the joint. To annotate *Cd2*^+^ *H2*^+^ myeloid precursors in the skin, their gene signature in the joint (“FindConservedMarker” with adjusted *P*-values < 0.01, average log FC > 0.5, average pct.2 < 0.2 while excluding ribosomal genes) were used in UCell (2.0.1) ^24^. Cells with a UCell score > 0.995 quantile were identified as *Cd2*^+^ *H2*^+^ myeloid precursors. To differentiate between the inflammatory and anti-inflammatory potential of the Kaede^GREEN^ and Kaede^RED^ phagocytes, UCell scores for the GO-BP gene sets GO:0050728 - anti-inflammatory response and GO:0050729 - pro-inflammatory response were calculated. Gene sets were extracted from org.Mm.eg.db (3.15.0) ^56^ using the AnnotationDbi’s (version 1.66.0) ^57^ “select” function. UCell scores were then compared separately between each cluster using the Wilcoxon rank-sum test and *P*-values were corrected for multiple testing using the Benjamini and Hochberg (BH) method. For visualisation logFC was scaled using the “scale” function in R without centring. Data were exported as *.h5ad for subsequent analyses in python using seuratDisk’s (0.0.0.9015, https://github.com/mojaveazure/seurat-disk) “SaveH5Seurat” and “convert” functions. Kaede^RED^ enriched subclusters were selected by modelling the Kaede^RED^ ratios using a quasibinomial generalised linear model GLM (“glm” function in stats R) due to overdispersion in the ratios (dispersion > 3,5; calculated by first fitting a binomial model and using the following formula: “sum(residuals(fit, type = “deviance”)^2) / fit$df.residual”). To account for the sensitivity to small total cell numbers, which can artificially inflate ratios, log(total) cell population in each cluster was included as a fixed effect (final formula: “cluster + strain + log(total)”). Sublining fibroblasts were selected as the baseline for comparison due to their stable Kaede^RED^ ratio and high total cell number. Resulted *P*-values were corrected for multiple testing using BH method.

### Post-processing of mesenchymal clusters (mouse)

Harmony integration over both strains was performed for fibroblasts after log-normalisation and regression of unwanted sources of variation such as mitochondrial gene distribution and cell cycle. The top 30 Harmony components were used for clustering and visualisation. The clusters identified by Seurat (top 30 Harmony components, resolution = 1) were manually annotated based on their marker gene expression. The subset of fibroblasts with the highest transcriptional differences between the strains was identified by first identifying the fibroblast-specific differentially expressed genes (DEGs) using “FindMarkers” in Seurat between strains on the entire joint dataset excluding fibroblasts, as well as on fibroblasts only, and then by subtracting the non-fibroblast significant (adjusted *P*-value < 0.01) genes from the significant fibroblasts-specific genes (adjusted *P*-value < 0.01) upregulated in C57BL/6. Finally, UCell score was calculated for these DEGs and compared among different fibroblast clusters by Seurat’s “RidgePlot”.

### Differential abundance analysis

scCODA (0.1.9) ^68^ was used for differential abundance analysis to account for the compositionality issue in scRNAseq cluster frequencies ^75^. In all cases scCODA “CompositionalAnalysis” was performed using default parameters in python (3.8.0). In human synovial myeloid scRNAseq datasets FOLR2^high^ LYVE1^+^ macrophage cluster was chosen as the reference clusters as we expected no major change in abundance for these resident cell population ^18^. Sex and age (Z-normalized) was accounted for, and the FDR threshold was increased gradually to 0.4 as first credible effect was observed. In human skin dataset reference was chosen automatically by the scCODA and sex was accounted for, and FDR threshold was set to 0.05. In human PBMC dataset reference was set automatically by scCODA and FDR threshold was set 0.05. Additionally edgeR ^76^ differential abundance test on cluster quantities was performed using diffcyt’s (1.22.0) ^58^ “testDA_edgeR” with default parameters. Neighbourhood-based differential abundance was performed using MiloR (1.10.0) ^59^ by creating the neighbourhood graph using “buildGraph” (“k” = 20 and “d” = 30, with Harmony components or in case of scVI integration using scVI embeddings), “makeNhoods” with “prop” = 0.1, “countCells” with the biological replicate variable as the samples, and “testNhoods” with “fdr.weighting” set to “neighbour-distance”. For visualisation using “plotDAbeeswarm”, Spatial FDR α was set to α < 0.05. For mouse fibroblasts, MELD (1.0.0 on Python 3.8.8) ^38^ was used for differential composition analysis. Harmony components were used to generate the graph using the “Graph” function of graphtools (1.5.2, https://github.com/KrishnaswamyLab/graphtools), which was the input to the “fit_transform” function of MELD to calculate the strain-associated densities. These densities were L1 normalised to calculate the strain-associated relative likelihoods for each individual cell, and visualised per fibroblast subcluster.

### RNA velocity and CellRank

To generate the spliced and unspliced RNA counts, Velocyto workflow (0.17.17 on Python 3.10.2) ^64^ for droplet-based 10x Chromium libraries was used. RNA velocities were estimated using scVelo (0.2.4 on Python 3.9.10) ^65^ in the dynamic mode with default parameters with the following sequence of functions: “scv.pp.moments”, “scv.tl.recover_dynamics”, “scv.tl.velocity” and “scv.tl.velocity_graph”. CellRank (2.0.2) ^23^ was used to create a transition matrix by equally combining the velocity vector and the connectivity kernel. After initialising the generalised Perron cluster analysis (GPCCA) ^77^ estimator, the “fit” function was applied to the combined kernel with n_states between 5 and 13, using the reference-based clustering annotations to identify the optimal macrostates. The “predict_initial_states” function was used default parameters to determine the initial macrostates. To predict the terminal states, the “predict_terminal_states” function was used with the method set to “top_n” and n_states set to 1. The “recover_latent_time” function was used for pseudotime estimation. To visualise the coarse-grained transition matrix, “plot_coarse_T” was used with “order” = “incoming”.

### Trajectory differential expression analysis

Trajectory differential expression analysis was performed on the Kaede^RED^ myeloid subset using TradeSeq (1.10.0) ^32^. Using the latent time estimated by CellRank as the pseudotime, with the binarized strain considered as the “cellWeights” parameter, the “fitGam’’ function was applied to the SCT-corrected counts of the highly variable genes to produce negative binomial generalised additive model (NB-GAM) smooth expression values for each strain along the differentiation. To find the differentially expressed genes between strains along the pseudotime “diffEndTest” was implemented. Genes with *P*-value < 0.01 and according to the “cellWeights” parameters, a positive logFC, were assigned to BALB/c, while genes with a negative logFC were assigned to C57BL/6. To generate a smoothed expression of the overall gene signature for each strain, UCell scores were generated with these strain-specific genes and fitted to NB-GAM using TradeSeq’s “fitGam”. “PlotSmoothers” was used to plot the smoothed expression over pseudotime. The strain-specific genes were also used as input to gene ontology functional overrepresentation analysis (see below).

### Gene Ontology Overrepresentation analysis

For all functional enrichment analysis the same setting was used for the overrepresentation analysis using the ClusterProfiler (4.4.4) ^62^ “enrichGO” function with background set to NULL, using the org.Mm.eg.db (3.15.0) ^56^ dataset, ontology set to GO-BP and with “pAdjustMethod” = BH, “pvalueCutoff” = 0.01 and “qvalueCutoff” = 0.05. If two or more clusters are compared, then “compareCluster” was used with “fun” set to “enrichGO”. The selection of genes of interest was performed using either Seurat’s “FindConservedMarkers”, when conserved markers were considered among strains, or “FindMarker”. The selection criteria for significant genes were adjusted *P*-value < 0.01 and average |logFC| > 0.5.

### Ligand-receptor interaction analysis

Ligand-receptor interaction analysis was performed using CellChat (1.5.0) ^33^ on the SCT normalised assay including all annotated cell types separately for each strain. To detect enriched signalling pathways, the “rankNet” function was used after merging the CellChat objects, followed by row normalisation. Signalling pathways with more than 50% relative contribution in each strain were selected for downstream visualisation. Interaction probabilities were calculated for all annotated subclusters, but for visualisation their probability was summarised by grouping major cell types. In case of T cell interactions with other cell types, the interaction analysis was performed on BALB/c Kaede^tg^ mice scRNAseq dataset and all pathways were included.

### Quality control and processing of single cell RNA sequencing libraries (human)

A public dataset of human synovial scRNAseq dataset (E-MTAB-11791 ^26^) and the in-house healthy human synovial libraries (E-MTAB-14339) were subjected to a quality control to exclude low quality events by individually examining the distribution of quality criteria for each dataset, including the number of UMIs >= 1,400 / 1,000 number of genes detected >= 700 / 800, and mitoRatio < 0.4 / 0.3 and complexity (log10(genes/UMI) > 0.80 / 0.83, respectively. Genes expressed in fewer than 10 cells were excluded. Doublets were detected and removed using scDblFinder (1.13.12) ^61^ with default parameters. To overcome batch effects, the scVI (1.1.1, Python 3.9.18) ^78, 79^ integration was used. Scanpy (1.9.8) ^66^ log-normalisation and variable gene detection (seurat_v3) were performed on the top 3,000 variable genes. The scVI model was defined with categorical batch variables (library, dataset origin) and their continuous covariates (mitoRatio, cell cycle score) and trained with four layers and 30 latent layers. Clustering was performed using the Leiden algorithm in scanpy and clusters were manually annotated based on their marker gene expression. Myeloid cells were subset and subjected to a repeated scVI workflow with manual annotation to identify subclusters. Alra imputation ^80^ (as part of SeuratWrapper 0.3.4, https://github.com/satijalab/seurat-wrappers) was performed and the absence of T, B and DC markers in the *CD2-*expressing myeloid cluster was confirmed to exclude any *CD2*-expressing contaminating events originating from previously undetected DCs or lymphocyte doublets. Myeloid subclusters from PsA and healthy samples were subjected to scANVI reference mapping as described below.

A second public dataset of human synovial scRNAseq dataset (E-MTAB-8322 ^18^) was analysed separately. Quality control was performed in Seurat by removing cells with number of UMIs < 800, number of detected genes < 300, complexity < 0.8, mitoRatio > 0.01 and < 0.2. Genes expressed in fewer than 10 cells were excluded. Doublets were detected and removed using scDblFinder as described above. Harmony integration was performed across libraries after log-normalisation and regression of cell cycle scores followed by Seurat clustering (resolution = 0.3). Alra imputation was performed for gene expression visualisation as described above. E-MTAB-8322 only captured *CD1C*^+^ and *CCR7*^+^ DCs due to technical restrictions in cell isolation. Myeloid cells were subset and subjected to scANVI reference mapping together with the myeloid subclusters from PsA and healthy synovia as described below, while excluding the undetected subsets of DCs.

Filtered and annotated data were retrieved from a public scRNAseq dataset of human skin from healthy individuals and patients with psoriasis and AD (E-MTAB-8142 ^29^). Myeloid cells were subset and subjected to scANVI reference mapping together with the myeloid subclusters from PsA and healthy synovia as described below. Differential abundance analysis on the human skin scRNAseq dataset was performed as described above.

Filtered and annotated data were retrieved from a public scRNAseq/HTOseq dataset of human PBMCs from healthy individuals and patients with psoriasis, PsA and PsX (GSE194315 ^31^).

Datasets without HTO immunophenotyping were excluded. Myeloid cells were separately subjected to scVI integration as described above (2,000 variables genes, batch variable (library), continuous covariates (mitoRatio, heat shock gene score). Clustering using the Leiden algorithm in scanpy identified the original annotated clusters together with two doublet clusters co-expressing typical myeloid cell markers and either those for T and NK cells or B cells, which were excluded. Myeloid cells were down-sampled to 25 %, subset and subjected to scANVI reference mapping together with the myeloid subclusters from PsA and healthy synovia as described below. Differential abundance analysis on the human PBMC scRNAseq dataset was performed as described above.

scRNAseq datasets from matched skin and joint biopsies of three patients were analysed using Seurat (5.2.1) ^53^. Cells were filtered based on the following QC criteria: number of UMIs > 1800, number of genes > 1000, mitochondrial gene ratio between 0.008 and 0.22, and erythrocyte gene ratio (HBA and HBB) < 0.002. Genes expressed in fewer than 10 cells or those without annotation were removed. Harmony (1.2.3) ^54^ was used for integration. Clustering was done using Seurat “FindCluster” (**Figures S1a and c**). Within each major cluster, subclusters were identified using Seurat’s “FindSubClusters” function. Subclusters of myeloid cells were annotated based on their top upregulated genes. Reference mapping of myeloid cells in skin and joint was performed using scANVI, following steps described in “Cross-tissue reference mapping” section below and based on our previously annotated synovial myeloid scRNAseq dataset (**Figures S1b and d**). Plasmacytoid dendritic cells (pDCs) were not included in the skin scANVI analysis due to their absence in the skin samples.

### Cross-tissue and -species integration and reference mapping (scANVI)

Reference mapping was performed using scANVI according to the scvi-tools (1.1.1) ^66^ guidelines to map annotations between the pairs of reference/query datasets described above. The workflow included merging the query and reference datasets, Scanpy log-normalisation and variable feature selection (3,000 variable features using “seurat_v3”), scVI model construction (4 layers, 30 hidden layers, with sample and organism or tissue as categorical batch variables and predetermined continuous covariates), preparation of missing labels on the merged object by setting the query datasets cluster labels to “unknown”, scANVI model setup and training with 100 max epochs and 100 samples per label, and finally the scANVI “predict” function. In order to create a co-occurrence heat map, a matrix was created by evaluating the ratio of mapped annotations in each of the originally annotated clusters of the query dataset using the following script: query_metadata.groupby([‘origninal_label’, ‘scANVI_label’]).size().reset_index(name=’total’).groupby(‘origninal_label’)[’total’].transfor m(lambda x: x / x.sum()). A scaling was performed on the ratios using “sklearn.preprocessing” “scale” function for visualisation. In the case of cross-species mapping, genes in the mouse dataset were translated into human orthologs using the “translate” function in mousipy (0.1.5, https://github.com/stefanpeidli).

### Single cell correlation analysis

To asses gene correlations in scRNAseq datasets of human PsA and healthy fibroblasts, we created meta cells using hdWGCNA (0.3.00) ^60^ “MetacellsByGroups” function on the Alra imputed expressions. Correlation was measured between expression levels in the meta cells with Pearson analysis.

### Analysis of MAESTER libraries

#### Mitochondrial variant calling

Mitochondrial variant calling was performed following the steps in the MAESTER protocol ^81^ and scripts provided in the original publication ^30^. The raw sequencing files were first assembled using the “Assemble_fastq.R” script. Then, 24 nucleotides were trimmed from the 5′ end using homerTools (v4.11.1) ^48^. Reads were aligned to the GRCh38-2024-A reference genome using STAR (v2.7.3a) ^49^, and mitochondrial chromosome reads were extracted using samtools (v1.10)^50^. Barcode and UMI sequences were moved from the read ID into the BAM file using the “Tag_CB_UMI.sh” script. Variant calling was performed using maegatk (v0.2.0 https://github.com/caleblareau/maegatk) with the command: “maegatk bcall -i SAMPLE_chrM.10x.bam -z --snake-stdout -mr 3”, as recommended by the protocol. The skin sample from patient 2 was randomly down sampled to 30 percent to improve processing speed. Mitochondrial DNA coverage was visualized using the script provided in the original MAESTER manuscript ^30^ “MT_coverage.R” (**Figures S4a**).

#### Identification of informative mitochondrial variants

Following the MAESTER protocol, we identified informative mitochondrial variants specific to each subcluster of cells in the skin and joint samples, separately for each patient. Clusters with fewer than 50 cells were excluded due to imbalance between samples and possible instability in variant identification. For each variant and subcluster, we calculated quality metrics using provided scripts “8.1_MGH252_voi.R”, including mean quality (mean base quality score across all cells where a variant was observed on either strand), mean and 99th percentile allele frequency (AF) or heteroplasmy, and mean coverage.

Variants with quality scores below 30 were filtered out. Variants were considered informative if they had mean coverage >5, 99th percentile heteroplasmy >5% (i.e., at least 1% of the cells have a heteroplasmy level exceeding 5% for this variant) and if their 99th percentile heteroplasmy was 5 times higher than in all other clusters, except those of the same lineage (allowing us to keep shared variants that might be present across related cell types). To confirm the cluster specificity of these variants, we calculated the mean variant allele frequency (VAF) over all the variants for each cluster and visualized the results using a heatmap (**Figures S4b**). High values on the diagonal indicated that the variants were specific to their assigned clusters.

#### Cross-tissue mitochondrial variant comparison

We intersected informative variants between each pair of subclusters across skin and joint samples. For each major cell type shared variants were defined as the union of common variants found across all associated subclusters. In the joint myeloid subclusters, variant identification was performed using the manual annotation in order to avoid potential bias introduced by label transfer as opposed to manual clustering. However, for the final visualization, we used the scANVI labels. For each scANVI-defined cluster, variants were aggregated based on their corresponding manual annotations (**Figures S1b**). Statistical significance was tested using a one-vs-rest Wilcoxon rank-sum test, with p-values adjusted using the BH correction.

#### Mitochondrial variants identification robustness analysis

Given the sensitivity of informative variant identification to threshold settings, as well as the overall differences in data quality between skin and joint samples, possibly due to ongoing inflammation in the skin, we repeated the common variant identification process across 1,000 iterations using varying thresholds. In each iteration, we changed the thresholds for mean coverage (from 1 to 10), the skin 99th percentile heteroplasmy (from 1 % to 10 %), and the joint 99th percentile heteroplasmy (from 1 % to 10 %). For each cluster or subcluster in each iteration, we recorded the median rate of shared variants across the three patients, as well as the rank of this rate compared to other clusters. The results were visualized using boxplots (for medians) and heatmaps (for ranks).

#### Graph-based analysis of mitochondrial variant sharing

Graph analysis within the myeloid subclusters in the joint was performed using igraph (2.1.4) ^63^, where each subcluster was represented as a node. Edges between nodes were weighted based on the number of shared variants between each pair of subclusters (**Figures S1e)**. This was done both including and excluding skin-joint common variants. Graph strength for each node was calculated using the “strength” function from the igraph. Strength values were then compared using a one-vs-rest Wilcoxon rank-sum test, and *P*-values were adjusted using the BH method.

### Imaging mass cytometry

Synovial tissue slides were deparaffinised by heating the slides for 30 min at 65 °C and washing in Histo Clear (National Diagnostics) and rehydrated through a series of 100 % (v/v) ethanol, 95 % (v/v) ethanol, 80 % (v/v) ethanol, 60 % (v/v) ethanol, water. Heat-induced epitope retrieval was performed for 30 min at 95°C in Tris-EDTA buffer (10 mM Tris, 1 mM EDTA, pH 9.2). After cooling down for 15 min, blocking was performed in TBS-T supplemented with 3 % (w/v) BSA and 1 % rabbit serum for 1 h at ambient temperature. A pre-composed panel of metal labelled antibodies (**Table S3**) was applied to the sections. Antibodies were pre-labelled with metals or conjugated according to the manufacturer’s instructions of the Maxpar X8 Multimetal Labeling Kit (Standard Biotools). Indium chloride was purchased from Sigma-Aldrich (natural abundance, 96% of ^115^In), ^113^Indium was purchased from Trace Science. Incubation with antibodies was performed over for 16 h at 4 °C in the presence of 0.3 µM Ir-intercalator (Standard Biotools). Slides were washed twice in TBS-T for 10 min, followed by a brief wash in deionized water, air dried and stored in a desiccated environment until analysis. Acquisition was performed at 200 Hz on a Hyperion Imaging System (Standard Biotools, CYTOF software v. 7.1). MCD files were processed using Steinbock (v. 0.16.0) ^47^. imcRtools^47^ (v. 1.3.4) and R (v. 4.3.2) was used to load the data and construct a SpatialExperiment ^51^. An inverse hyperbolic sine (arcsinh) transformation with a coefficient of 1 was applied to all the datasets. The data were normalized by computing a z-score, which used the entire dataset as reference. All data was exported as FCS and further analysed with FlowJo Software (v. 10.10, BD Biosciences).

### Spatial analysis of imaging mass cytometry data

Cells were annotated by gating on the known markers using FlowJo software. After annotation, the cellular neighbours were identified by using the imcRtools R package ^47^. Cellular neighbours were defined by a centroid extension of 40 µm. To quantify interacting cells, neighbours of the cell population of interest were first retrieved and interacting cells were defined as those that had at least one neighbour of the target cell population. Cellular pairwise interactions were analysed using the imcRtools R package. The statistical significance of each pairwise interactions was assessed by permutation testing ^82^. This approach quantified the average number of neighbouring cells for each subpopulation and compared the results to a null distribution generated by randomly reassigning cell type labels over 1,000 permutations. Pairwise interactions with *P*-values below 0.05 were considered as statistically significant. Attraction was defined as a higher number of neighbouring cells of a specific type than would be expected in a random distribution, whereas avoidance was defined as a lower number of such neighbours than would be expected in the sample. The results are visualized by dot plots generated by using the R package ggplot2. The dot size represents the median proportion of statistically significant interactions found across samples.

### Statistical analysis

Statistical analysis of non-sequencing data was performed using Prism 9. Unless otherwise stated, all data are reported as median, interquartile range and minimum-maximum. No statistical methods were used to pre-specify sample sizes, but our sample sizes are similar to those reported in previous publications. Parametric and non-parametric analyses were used where appropriate. Differences were considered significant when *P* < 0.05. Corrections for multiple testing were made as appropriate.

## Data availability

Single cell sequencing data supporting the results of this study have been deposited in the Gene Expression Omnibus (GEO) under accession GSE228629 and in the European Nucleotide Archive (ENA) under the ArrayExpress accession E-MTAB-14339. Publicly available datasets reanalysed in the study can be accessed under accession numbers: ENA-ArrayExpress E-MTAB-11791 and E-MTAB-8322 (human synovia), ENA-ArrayExpress E-MTAB-8142 (human skin), GEO GSE194315 (human PBMC) and GEO GSE122108 (ImmGen mouse mononuclear phagocytes).

## Code availability

All the methods and algorithms used in this manuscript are from previously published studies and are cited in the methods section. A comprehensive list about all software used can be found in the **Table S3**. Selection criteria, thresholds and other essential parameters are stated in the methods section. No new method for data analysis was developed. Additional scripts to reproduce the analyses are available from the authors upon request.

